# Ecological Stability Emerges at the Level of Strains in the Human Gut Microbiome

**DOI:** 10.1101/2021.09.30.462616

**Authors:** Richard Wolff, William Shoemaker, Nandita Garud

## Abstract

The human gut microbiome harbors substantial ecological diversity at the species level, as well as at the strain level within species. In healthy hosts, species abundance fluctuations in the microbiome are thought to be stable, and these fluctuations can be described by macroecological laws. However, it is less clear how strain abundances change over time. An open question is whether individual strains behave like species themselves, exhibiting stability and following the macroecological relationships known to hold at the species level, or whether strains have different dynamics, perhaps due to the relatively close phylogenetic relatedness of co-colonizing lineages. Here, we analyze the daily dynamics of intra-specific genetic variation in the gut microbiomes of four healthy, densely longitudinally sampled hosts. First, we find that overall genetic diversity in a large majority of species is stationary over time, despite short-term fluctuations. Next, we show that fluctuations in abundances in approximately 80% of strains analyzed can be predicted with a stochastic logistic model (SLM)—an ecological model of a population experiencing environmental fluctuations around a fixed carrying capacity which has previously been shown to capture statistical properties of species abundance fluctuations. The success of this model indicates that strain abundances typically fluctuate around a fixed carrying capacity, suggesting that most strains are dynamically stable. Finally, we find that the strain abundances follow several empirical macroecological laws known to hold at the species level. Together, our results suggest that macroecological properties of the human gut microbiome, including its stability, emerge at the level of strains.

## Introduction

The human gut microbiome is a complex ecological community, composed of tens of trillions of cells that interact directly and indirectly with one another and the host [20, 44, 45]. Though the precise species composition of the gut microbiome differs between hosts, healthy adult guts tend to be both ecologically diverse and temporally stable at the species level under normal circumstances [21, 23, 29, 40]. The ecological stability of the gut community is critical to the preservation of its functional capacity through time, and periods of instability and heightened variability are often associated with environmental perturbation or disease states [9, 22, 24].

Much as the gut community as a whole is made up of a diverse array of species, within species, populations of gut microbes harbor many genetic variants [38]. A growing body of literature highlights the importance of inter-host differences in microbiome genetic composition for various aspects human health, with specific microbial genotypes and strains associated with the digestion of certain foods [25], a range of host disease risk factors [7], bile and lipid composition [8], and antibiotic resistance [10]. Recent studies have begun to characterize how this genotypic diversity changes through time within hosts [15, 46, 38, 40], and have linked longitudinal changes in genetic composition to specific host phenotypes and metabolite levels [6].

Broadly, dynamic changes in genetic variation in the gut microbiome occur at two distinct levels. First, there are changes in the frequencies of lineages that have clonally diverged since their common ancestor colonized the host due to the evolutionary forces of mutation, drift, selection, and recombination. Typically, such lineages differ from one another at a small number 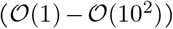 of single nucleotide variants (SNVs) [46, 15]. Second, there are fluctuations in the relative abundances of con-specific strains that do not share a common ancestor within the host. When levels of recombination between such strains are sufficiently low, clonal descendants of the initial colonizers may persist within the host as genetically distinguishable populations differing from one another at a similar number of sites in their shared, core genome 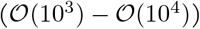 as strains drawn from unrelated hosts [79, 38, 73, 78, 15]. Multiple colonization by con-specific strains is evidently under some degree of ecological constraint, as only a small handful of strains (typically between one and four) are ever observed within a host at any one time—a phenomenon dubbed “oligo-colonization” [36, 15, 69, 14]. The mechanisms enabling a small number of strains to colonize a host and rise to high frequency, but preventing a large number of exogenous strains from doing the same, are not known as yet. Interestingly, similar colonization patterns have been observed in a number of other host-associated microbiota, both at different human body sites [57], and in other organisms [27, 13].

In healthy, adult hosts, a large majority of strains persist over periods of months to years [38, 40, 31, 6, 26, 15]. Moreover, strains within the gut can remain resilient in the face of large perturbations, such as antibiotics [36] and fecal microbiome transplants [16]. However, little is known about the magnitude of daily fluctuations in genetic composition at either the strain or lineage level under ordinary conditions, or how such fluctuations ultimately affect the stability properties of the gut community, as longitudinal studies of genetic diversity in the gut have tended to focus on samples collected at multi-month intervals.

In this work, we seek to understand how the genetic composition of the gut changes through time, from daily to multi-year timescales in four healthy, adult hosts sampled over the course of six to 18 months [34]. To do so, we leverage concepts from macroecology to examine the dynamics of strains in the these four hosts. Macroecology focuses on characterizing statistical regularities in patterns of abundance and diversity within and between ecological communities. A growing body of work has demonstrated that species-level patterns of diversity in a variety of natural microbial communities are well-described by macroecological laws [1, 60, 19, 11]. Many of these macroecological laws can be recapitulated through intuitive ecological models containing few if any free parameters [1, 11, 59]. Among these successful models is the Stochastic Logistic Model (SLM), which describes the dynamics of a population experiencing rapid stochastic fluctuations induced by environmental noise around a fixed carrying capacity [80]. Whether the populations making up a community exhibit regular, statistically quantifiable dynamics, and if so, whether these dynamics can be explained using simple models, are fundamentally macroecological questions. In this work, we find not only that a large majority of strains in these healthy hosts exhibit abundance dynamics consistent with an SLM, but also that strain abundance fluctuations follow several macroecological laws known to hold among species [1, 11, 59, 60].

Together, our results indicate that daily fluctuations in overall genetic composition within the gut microbiome are largely stationary, and that these fluctuations follow broad macroecological patterns. Thus, several macroecological properties of the human gut known to hold at higher levels of taxonomic organization, including its stability, appear to emerge at the level of strains.

## Results

To explore how daily fluctuations in nucleotide diversity and strain abundances translate into stability over periods of months to years, we used high-resolution temporal data from four hosts sampled in the BIO-ML project [34] (see Materials and Methods for further details on sampling).

### Temporal stability of intraspecific genetic variation

Within hosts, allele frequencies change through time in gut microbial populations due to mutation, drift, selection, and fluctuations in the relative abundances of strains. While studies examining broad cohorts of sparsely longitudinally sampled individuals indicate that the magnitude of intra-host fluctuations only infrequently approaches that of inter-host differences over timescales of months to years [38, 40, 15], more finely-resolved temporal trends are less well characterized. To evaluate the stability of the gut community, it is crucial to determine whether temporal fluctuations are stationary or directional.

To assess the stability of intra-specific genetic variation through time, we examined temporal trends in patterns of nucleotide diversity within our four hosts using *F_ST_*, a standard measure of genetic differentiation between populations (see Materials and Methods for further details on *F_ST_* calculations). If the genetic composition of a species changes directionally, we expect that samples drawn later in the timecourse will have greater *F_ST_* relative to the initial timepoint than earlier samples. If, by contrast, fluctuations are stationary, then later timepoints should on average be no more diverged from the initial sample than earlier timepoints.

To contextualize the magnitude of variation in genetic composition through time, we normalized our longitudinal measurements of *F_ST_* within hosts by the mean *F_ST_* value of the species across hosts. We calculated this species-wide mean *F_ST_* using shotgun metagenomic data from 250 North American hosts sampled in the Human Microbiome Project [28, 29], allowing us to better capture the extent of inter-host diversity. We refer to the resulting normalized *F_ST_* statistic, obtained by dividing each intra-host *F_ST_* measurement by the mean *F_ST_* across hosts, as 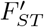. 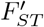 will approach or exceed one when intra-host fluctuations are of the same magnitude as inter-host differences, and will remain close to zero when the genetic composition of the population is constant over time.

In line with previous work [38, 40], we observe that changes in genetic composition within the hosts we examine (*am, ao, an*, and *ae*) rarely approach the magnitude of inter-host differences, as 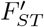 remained well below one at all timepoints for all but one species examined (Figure S5 in S1 Text). For the one aberrant species, *Faecalibacterium prausnitzii* in host *ao*, 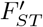 increases steadily before a rapid rise above one around timepoint 60 (Figure 1B). More typical, however, is the example of *Bacteroides vulgatus* in host *am*, for which 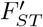 fluctuates but appears to remain near a long-term steady state (Figure 1A).

**Figure 1:**
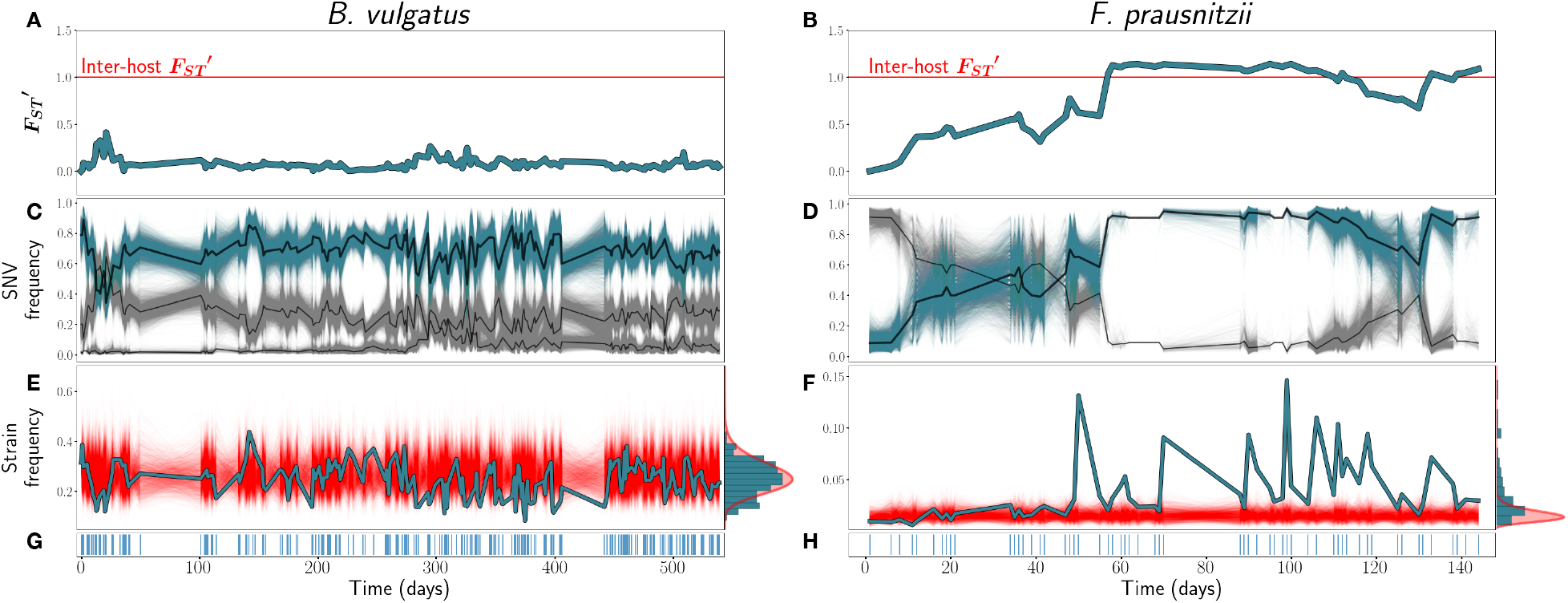
**A & B**) *F_ST_′* trajectories for *B. vulgatus* (*am*) and *F. prausnitzii* (*ao*), respectively. **C & D**): SNV frequencies of three inferred strains for B. vulgatus (C) and two inferred strains for F. prauznitzii (D). In black are the inferred strain trajectories. Highlighted in blue are example strains featured further in (E) and (F). **E&F**) Frequencies of the example strains in blue, with simulations of the corresponding SLM overlaid in red. At right, the empirical distribution of strain abundances is plotted in blue, and the stationary Gamma distribution of abundances (see Equation 2) predicted by the SLM in red. **G & H**) Sampling time points. Blue lines indicate that a sample was taken on that day.

To determine quantitatively whether fluctuations in genetic composition were stationary or directional, we implemented an Augmented Dickey-Fuller test (ADF) for each 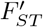 time series [77]. The ADF tests the null hypothesis that a time series is non-stationary against the alternate hypothesis that the time series is stationary. Rejecting the null hypothesis with the ADF is thus evidence that the mean and variance in a temporally varying quantity are time invariant. For 34 of the 45 species (76%) considered, we rejected the null hypothesis at a significance level of *p* = 0.05, indicating that the majority of species exhibit stationary 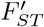 trends.

Our results suggest that fluctuations in allele frequencies are typically stable, at least when coarse-grained across the whole genome, in these hosts. However, large-scale, rapid changes in allele frequency, such as observed in *F. prausnitzii*, also occur.

### Strain frequencies

Recent empirical studies using isolate, single-cell, and shotgun sequencing data have demonstrated that at any one time, the human gut microbiome is colonized by at most a handful of distinct conspecific strains (between one and four) which do not share a common clonal ancestor within the host [31, 34, 15, 78, 69]. Much less is known, however, about the dynamics of strains once they have colonized the host.

To investigate these dynamics, we infer strain genotypes and frequencies using an algorithm adapted from by Roodgar *et al*. This algorithm identifies large clusters of SNVs with tightly correlated frequency trajectories [36], indicative of linkage on a common genomic background. By identifying large clusters, we expect to distinguish the trajectories of deeply diverged strains (for further information on our strain phasing, see S1 Text section 3.2). When no large cluster of tightly linked SNVs was detected within a species, we inferred that only a single strain was present.

Of the 45 species/host pairs examined, 15 (33.3%) harbored multiple strains. Of these 15, 13 were colonized by two strains, while in two separate hosts (*am* and *an*), the species *B. vulgatus* was composed of three strains. The number of fixed differences between strains varied between 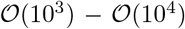, though due to our conservative filters for both calling SNVs from reads and assigning SNVs to strains, these likely represent underestimates of the true divergence between these strains. For further discussion of inter-strain genetic divergence, see S1 Text section 3.1.

As may be expected given our 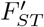 results, most strains, by visual inspection, exhibit frequency dynamics that are heuristically consistent with stationarity (see Supplementary Text S2—S5 for strain trajectories of all species). The relative frequencies of most strains appear to fluctuate around a constant value throughout the sampling period. In a typical example of this kind of behavior, the dominant strain of *B. vulgatus* in host *am* fluctuates around 60% frequency for more than 500 days (Figure 1C). However, in a minority of cases, strain frequencies shifted dramatically throughout the sampling period. The most striking example of this is *F. prausnitzii* in *ao*, which we have already seen underwent fluctuations in 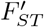 (Figure 1B) of the magnitude of inter-host differences in genetic composition. In this species, an initially rare strain almost fully supplants the initially dominant strain within the span of 60 days, before a partial reversion later in the timecourse (Figure 1D). In another case, *P. distasonis* in host *ao*, a single strain which initially falls below the detection threshold rises to detectable abundance midway through the sampling timecourse (S3 Text). Similarly, the minor strain of *B. vulgatus* in *am* is initially very rare before rapidly increasing in frequency around day 300, and subsequently reverting to an intermediate steady state below its maximum frequency. This subtle shift in strain frequencies is not, interestingly, detected by our ADF test of 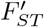 as a departure from stationarity—likely due to the fact that the absolute magnitude of the shift in allele frequencies between the beginning and end of the timecourse is small (less than 5%).

We further note the striking visual correspondence between the 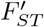 and strain frequency trajectories, as is evident in Figure 1 for both *B. vulgatus* and *F. prausnitzii*. Because fluctuations in the relative frequencies of co-colonizing strains with respect to one another determine allele frequencies at a very large fraction of all polymorphic sites, genome-wide average diversity statistics like 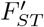 will reflect strain dynamics. This correspondence is therefore supporting evidence that the overwhelming majority of genetic variation in these species is due to fixed differences between strains, rather than among lineages belonging to strains. However, as is evident from the example of the minor strain of *B. vulgatus* in *am*, subtle but important strain dynamics can also be obscured when only considering genome-wide average patterns of diversity.

### Stochastic logistic model

In the preceding sections, we have seen that the genetic composition of most species examined exhibit stationary dynamics over the timescale of observation. We hypothesized that this behavior might result from the underlying strains fluctuating around fixed absolute carrying capacities.

To test this hypothesis, we assessed the fit and predictive capacity of the stochastic logistic model (SLM)—a model of a population experiencing stochastic fluctuations around a fixed abundance. Recent work in microbial ecology has demonstrated the power of minimal models like the SLM, requiring the fit of no free parameters, to reproduce qualitative and quantitative features of natural microbial community dynamics [1, 59, 58, 11, 80]. Here, we tested the capacity of the SLM to forecast future strain behavior when trained on an initial subset of timepoints. By training only on a subset of initial points, we can assess whether strain dynamics are consistent through time.

Under the assumptions of the SLM, each population *i* has a long-term carrying capacity *K_i_*, and temporal fluctuations in abundance around this value are driven by environmental noise with am-plitude *σ_i_*. The dynamics of a population governed by an SLM can be expressed with the following stochastic differential equation:

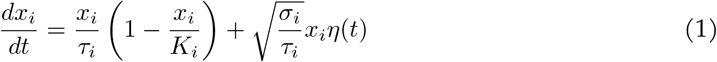

where 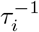 is the growth rate, and *η*(*t*) is a Brownian noise term.

Populations following an SLM may experience large fluctuations in abundance over short timescales, and may even be temporarily found far from their long-term average value, but these deviations will be transient. Over long timescales, the observed distribution of abundances will converge to a stationary Gamma distribution [1]:

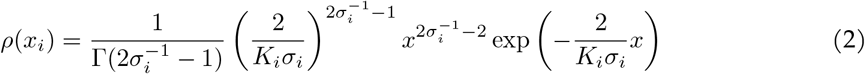

To determine whether strain trajectories could be described by an SLM, we first obtained time series of strain abundances by multiplying the relative frequencies of the strains inferred in the previous section by the relative abundance of the species to which they belong.

Next, we estimated *K_i_* and *σ_i_* from the first third of timepoints for each strain. *K_i_* and *σ_i_* are not free parameters, but rather are functions of the mean and variance of observed abundances (for details, see S1 Text section 4). To assess quantitatively whether the time series of strains in our cohort could be adequately described by an SLM, we developed and implemented a goodness-of-fit test. The test determines whether the transitions between subsequent timepoints are consistent with an SLM (for further details, see S1 Text section 5). Qualitatively, if a strain follows an SLM, its average and variance in abundance in the latter two-thirds of the time series should match that of the former third, and the strain should have a tendency to revert to its carrying capacity *K_i_*.

Returning to our case studies, we see that the dominant strain of *B. vulgatus* is well described by the SLM (Figure 1E). The true abundance trajectory (in blue) explores largely the same space as the SLM simulations (in red). Moreover, the empirical distribution of abundances across the entire timecourse appears to approach the stationary Gamma distribution (Equation 2) predicted by the SLM (Figure 1E, at right). By contrast, the dynamics of the invasive strain of *F. prausnitzii* deviate strongly from the SLM (Figure 1F).

Overall, 79% (49/62) of strains passed our goodness of fit test, with 83% passing in *am*, 86% passing in *ao*, 73% passing in *an*, and 69% passing *ae* (Figure 2). While the dominant strain of *B. vulgatus* passes, both strains of *F. prausnitzii* in *ao*, as well as the minor strain of *am*, fail the SLM. Thus, the SLM recapitulates the qualitative behavior of a large majority of strains even when trained on only a subset of initial points, while also having power to discriminate instances in which the dynamics of strains are evidently quite non-stationary.

**Figure 2:**
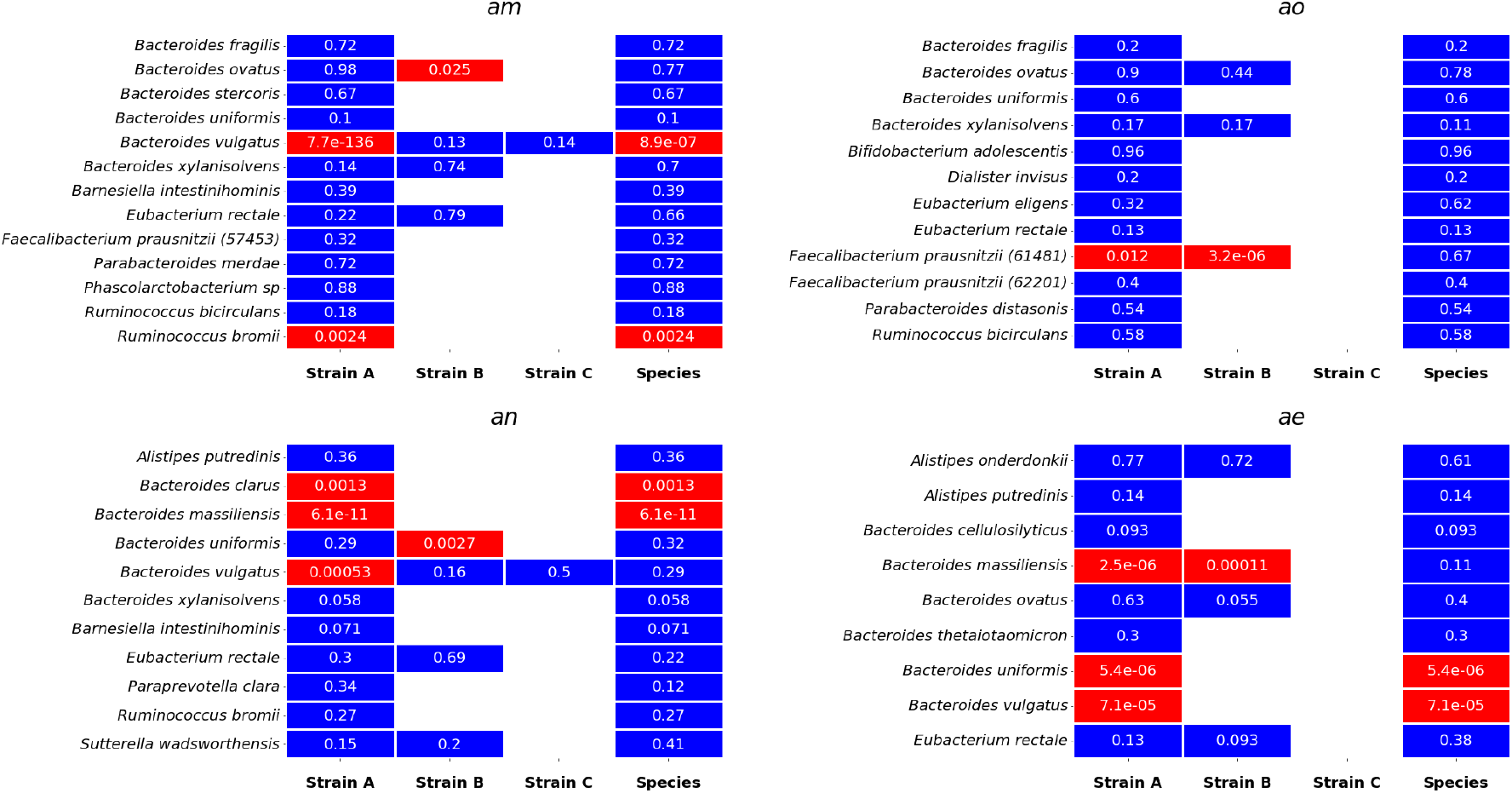
Results of the SLM goodness of fit test, by host. 79% (49/62) of strains, and 86.6% (39/45) of species, exhibit stochastic logistic dynamics across the sampling interval. The percentage of strains passing the test varied between hosts, with 83% passing in *am*, 87% passing in *ao*, 75% passing in *an*, and 69% passing in *ae*. The p-value associated with each strain or species is shown in white within each cell.

Moreover, the likelihood of exhibiting SLM dynamics is independent of the presence of other strains. Of the 32 strains for which another con-specific strain was present, 24 (75%) passed the SLM test, while among the 30 singly colonizing strains, 25 (83%) passed test. Thus, while singly colonizing strains tend to be moderately more likely to exhibit stable dynamics, the difference in pass rate is not statistically significant (χ^2^ = 0.24, p-value = 0.62), indicating that the presence of con-specific strains is not *prima facie* destabilizing for a focal strain.

Next, we conducted an identical test of the SLM at the species level. Overall, 84% (38/45) of species exhibited dynamics consistent with a SLM. In the case of *F. prausnitizii*, for instance, the abundance of the species overall fluctuated stably, obeying the SLM. Thus, despite a partial replacement event, the total abundance of the species remained roughly constant. One interpretation of this observation is that these strains strongly compete with one another for the same species’ niche. Interestingly, *F. prausnitizii* is known to experience higher rates of replacement over multi-year timescales that other gut commensals, and these replacements are associated with alterations in levels of plasma metabolites which affect host immunity [6]. However, we emphasize that the replacement was only partial, and the “displaced” strain recovered temporarily to intermediate abundance. This example highlights the complexity of strain dynamics, as well as their potential relevance for host phenotypes.

### Macroecology of strains

Much as the dynamics of individual strains can largely be recapitulated with a single relatively simple model, we can also attempt to parsimoniously characterize patterns of variability across strains collectively. Such low-dimensional representations of complex community dynamics are the natural purview of macroecology, which attempts to characterize variation within and among communities by observing the statistical patterns of abundance, distribution and diversity across their constituent members. In many kinds of microbial ecosystems, including the human gut, patterns of species abundance and distribution have been shown to broadly follow a number of macroecological laws, including Taylor’s Law and a Gamma distribution of abundance fluctuations [1, 60, 11]. We show here that these macroecological relationships also characterize patterns of variation in the abundance of strains across our cohort.

The first pattern examined is a power law scaling between the mean and variance of abundance, known in ecology as Taylor’s Law, which can be stated as:

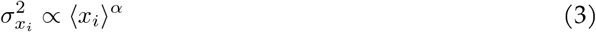

where 〈*x_i_*〉 and 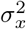 are the mean and variance of *x_i_*, respectively, and *α* is the scaling exponent of the power law.

Many mechanisms can give rise to Taylor’s Law. For instance, when the only source of variability between communities (or, in our case, longitudinal samples) is due to sampling noise, the Taylor’s Law exponent α will equal one. By contrast, in communities where the scale of fluctuations is independent of abundance—that is, where all populations have identical per-capita fluctuations—*α* will equal two [60, 1] (for further details, see S1 Text section 6). We observed a Taylor’s Law scaling with an exponent of *α* = 1.8 among all strains (Figure 3A), mirroring previous findings at the species level [60]. This Taylor’s Law exponent indicates that higher abundance strains are proportionally less variable than lower abundance strains (*α* < 2), and that variation in strain abundance is not driven solely by sampling noise (*α* > 1), but rather reflects true, underlying biological variability. However, the existence of a power-law scaling between mean and variance cannot, by itself, conclusively prove that any specific ecological model governs community dynamics. Indeed, the fit of the SLM does not depend on the existence of a Taylor’s Law scaling, or vice versa, as the SLM can hold with arbitrary mean and variance.

**Figure 3:**
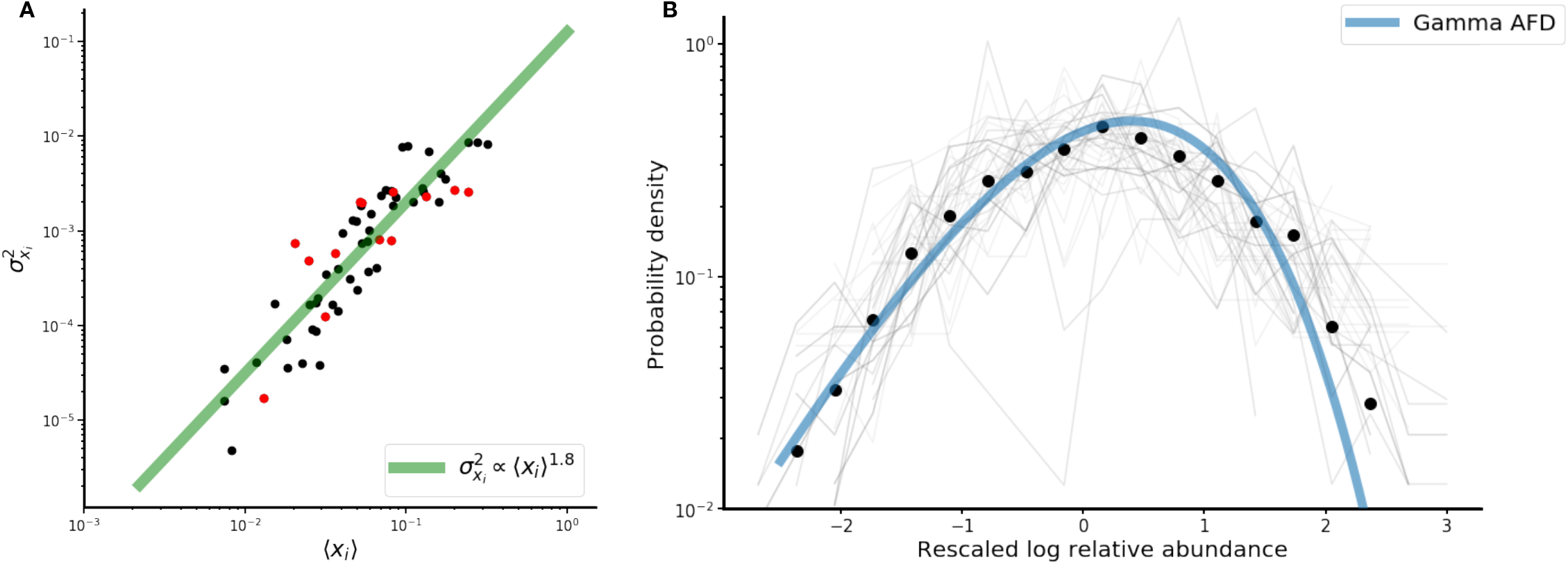
**A**) Scaling of variance in strain abundance with mean abundance obeys Taylor’s Law with exponent of 1.8. Black dots are strains passing the SLM, while red dots are strains failing the SLM. **B**) Strain abundances approximately follow a Gamma distribution, which is the stationary distribution of the SLM. Black circles are the average probability densities of the rescaled abundance across all strains, and the blue line is the Gamma fit of the bin means of the rescaled distributions. Light gray lines are the individual rescaled abundance distributions for each strain individually (62, in total).

The next pattern considered is the abundance fluctuation distribution (AFD), the overall distribution of abundances of a population through time. It is known that in a variety of microbial ecosystems, the AFD of many species tends to approach a Gamma distribution [1]. As discussed above, a population governed by stochastic logistic dynamics will tends towards a Gamma distribution of abundances over long timescales (see, for instance, the histogram of abundances for the dominant strain of *B. vulgatus* in Figure 1E, at right). Given the generally excellent fit of the SLM to the population time series, the abundances of strains might generically be expected to each individually follow a Gamma distribution. In Figure 3B, we see that the distributions of strain abundances are indeed, on average, well-described by a Gamma distribution (black dots, blue line), though some individual strains (grey lines) deviate somewhat from the Gamma stationary distribution. Recalling that the SLM of a given strain is uniquely determined by its mean and variance, it is apparent that the collapse of the AFDs to a single Gamma is in fact a consequence of the strong constraint Taylor’s Law places on these quantities across strains.

## Discussion

In this study, we sought to characterize the within-species population dynamics of the human gut microbiomes of four healthy hosts. Previous efforts have shown that within hosts changes in the genetic composition of gut microbial populations through time tend to be small compared with inter-host differences [38, 40]. We build on this result by demonstrating that at daily temporal resolution, intra-specific diversity tends to fluctuate around a long-term average value within the hosts examined over periods of years. We show, crucially, that the abundance fluctuations of a large majority of strains we detect can be predicted by the stochastic logistic model (SLM) of growth, a model that also recapitulates fluctuations at the species level [1, 11, 80]. Lastly, we find that empirical patterns of strain abundance variation in these hosts follow macroecological laws which have also previously been demonstrated to hold at the species level, including Taylor’s Law and a Gamma abundance fluctuation distribution [1, 11, 60]. Together, our results indicate that many of the broad properties of the gut observed at higher taxonomic levels of organization—such as its ecological and functional stability—may in fact emerge at the level of strains.

While the SLM was able to sufficiently describe strain dynamics for a majority of strains across species, its success was not universal, and deviations from this typical pattern were also informative. In one host, for instance, two strains of *F. prausnitzii* appear to undergo a rapid strain replacement, and fail the test. Whether this replacement was due to shifting environmental conditions, or direct inter-strain competition, is unclear. Regardless, our work indicates any successful description of gut microbial dynamics must incorporate the possibilities of both coexistence and rapid replacement. Over very long timescales, in fact, strain replacement may dominate the stable dynamics we observe here. Prior work [15, 26] suggests that over the course of decades, a large fraction of strains are ultimately replaced. One hypothesis is that this timescale reflects a waiting time for large environmental perturbations, such as antibiotics [36, 48] or bowel cleanses [64], but this is just one of many hypotheses. Indeed, this hypothesis is partially challenged by [36], where the strain content of an adult gut was perturbed during a course of antibiotics, but ultimately largely recovered to its pre-treatment state. This antibiotics study is a powerful demonstration of the stability of strains, even in the face of large perturbations. Investigating the possible explanations for the discrepancy between years-long and decades-long population dynamics at the strain level is an important problem that can be addressed with more extended timescales of observation.

While in this work we characterize the population dynamics of strains as ecological units, strain are by no means internally genetically homogeneous. The deep divergences we detect between conspecific strains 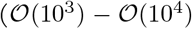 SNVs) are in fact genetic backgrounds, representing timescales of divergence likely far preceding colonization of the host. However, individual lineages bearing these backgrounds can differ from one another both at sites in the core genome and in gene content, and the relative frequencies of these different lineages with respect to one another can change due to evolution. Previous studies have shown that, at the level of lineages belonging to a strain, populations of gut microbes can experience both rapid selective sweeps [15, 34] and diversification into stably coexisting lineages [46].

How evolution impacts the ecological dynamics of strains and how in turn these ecological dynamics constrain and channel evolution, is an active area of research [30]. In the context of the SLM, these eco-evolutionary feedbacks can viewed as tuning a strain’s carrying capacity *K_i_*, growth rate 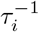, and sensitivity of the growth rate to environmental perturbation *σ_i_*. Naively, it is expected that evolution would tend to increase carrying capacity while minimizing the sensitivity of growth to abiotic fluctuations, but evolutionary modifications driving changes in one quantity may affect the other. The observed power-law scaling between the mean and variance in abundance (Taylor’s Law) is, in essence, a constraint on *K_i_* given *σ_i_*, and vice versa. The SLM thus not only describes ecological dynamics, but also, in conjunction with the empirical observation of macroecological laws, provides a useful framework for investigating the ecological effects of adaptation.

The SLM is ultimately a phenomenological model, not a mechanistic one, and its success at the strain level does not explain why strains coexist. How and why closely related strains coexist in the human gut is one of the central biological questions raised by our results. Spatial segregation between strains, perhaps occupying different colonic crypts, or partitioning luminal and mucosal niches, could contribute to the observed pattern of strain coexistence [33, 42, 82], much as it does among *Cutibacterium acnes* strains inhabiting different pores on the facial microbiome [57]. However, spatial structure is far from the only mechanism that can foster coexistence between strains. For instance, genetic differences at polysaccharide utilization loci content may contribute to intrahost metabolic niche differentiation, potentially favoring coexistence of closely related strains [81]. Moreover, stably coexisting strains have been reported under laboratory conditions, as well [5, 13]. In these experiments, strains may coexist by finely partitioning some aspect of the abiotic environment or by engaging in ecological interactions (e.g. cross-feeding), or by some combination of both. Indeed, recent theoretical work suggests that even subtle differences in resource uptake rates in high and low nutrient conditions may, in the presence of a temporally variable environment, lead to the coexistence of small numbers of closely related strains [76]. Investigating which of these mechanisms promotes strain coexistence in the human gut microbiome, and identifying the relevant genomic architectures, is an important avenue for future research.

We note that the four hosts examined here are not a representative sample of the full diversity of human lifestyles. For instance, all hosts were between the ages of 21 and 37, and residing in the United States at the time of sampling. The proportion of strains exhibiting stable or unstable dynamics may vary in different cohorts, but the tests we developed here will nonetheless be useful in identifying such differences. A future avenue of work will be to assess the generality of these findings in different cohorts—using new time series which are at least as long and densely temporally sampled as the BIO-ML data analyzed here, and of similar quality—including diseased or perturbed cohorts which may exhibit quite different dynamics.

Finally, our work highlights the importance of strains in understanding community structure and dynamics in the human gut microbiome [13, 76]. The ambiguity surrounding the bacterial species concept is well known [50] and reasonable alternatives have been proposed [32], but operationally species are, nonetheless, the predominant focus of attention in gut microbiome ecology. This focus is reasonable, as within-host strain structure is a comparatively recent discovery [69, 15, 31] and 16S rRNA sequencing provides an inexpensive, high-throughput means to examine community dynamics. However, it is reasonable to propose that for the human gut, and perhaps other microbial ecosystems, many higher-level macroecological patterns of abundance and diversity may originate at the level of strains.

## Materials and Methods

### Longitudinal data

To investigate the temporal dynamics of the human gut microbiome, we analyzed densely sampled shotgun metagenomic time series data from four hosts from the BIO-ML project [34]. By analyzing shotgun metagenomic sequences, we capture longitudinal patterns of intra-specific genetic variation for many bacterial species in these communities simultaneously. A total of 402 samples were drawn from these four individuals (aliases *am*, *ao*, *an*, and *ae*), with 206 samples coming from *am*, 74 from *ao*, 63 from *an*, and 59 from *ae* (see Table S1 and Figure S2 in S1 Text for further details on sampling). All four hosts were healthy adults between the ages of 21 and 37, of whom three were male and one female, and all were residing in the USA at the time of sampling. Crucially, for our purposes, these hosts were sampled at very fine temporal resolution, with a median interval between successive samples of either one or two days in each host, over a period of five (*ao*) to 18 (*am*) months.

### Aligning reads

To call single nucleotide variants (SNVs) and gene content, we aligned shotgun metagenomic reads to a panel of species which were prevalent and abundant within each host using MIDAS [18] (see S1 Text section 2 for further details on the bioinformatic pipeline employed). In total, we detected 45 species across the four hosts which met our coverage and prevalence criteria.

### Stationarity of intra-specific genetic variation

To calculate *F_ST_*, a measure of sub-population differentiation, we used the estimator:

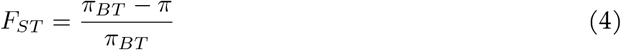

where *π* is the nucleotide diversity within a population and *π_BT_* is the level of diversity between populations.

The nucleotide diversity *π* is a classical population genetic measure of polymorphism, representing the average number of pairwise SNV differences between randomly chosen members of a population. To determine *π* for a given species within a sample, we used the estimator:

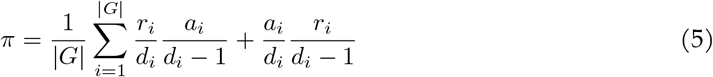

where *r_i_* is the count of the reference allele at site *i, a_i_* is the count of the alternate allele, *d_i_* is the depth of coverage, and |*G*| is the total number of sites in the genome. This quantity was calculated after first excluding sites with low read depth (< 5x), as reliable estimates of true allele frequency cannot be made for such sites. Our *π* calculations follow the same methodology as [38]—an early, foundational work characterizing patterns of genetic diversity in gut microbial populations within and across hosts—and are meant to be directly comparable.

Similarly, *π_BT_*, the diversity between timepoints, was calculated:

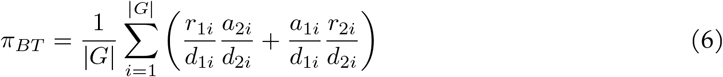

where *r_ji_, a_ji_*, and *d_ji_* are the reference allele count, alternate allele count, and depth of coverage of site *i* in sample *j*, and |*G*| is the total number of sites in the genome.

We calculated *π* for each species in each sample in each host, and *π_BT_* for each sample relative to the initial timepoint.

To calculate *F_ST_* between the initial timepoint (sample *i*) and sample *j*, we used the formula:

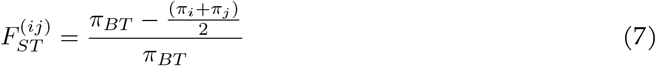

Lastly, we obtained our normalized statistic 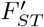 by dividing 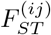 by the species mean inter-host *F_ST_*, estimated from a panel of 250 North American subjects sequenced by the Human Microbiome Project [28, 29]. To determine this species mean *F_ST_*, we first calculated the pairwise *F_ST_* for each pair of samples in which a species appeared, and then took the mean of these values.

To implement the Augmented Dickey-Fuller test on each time series of 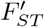 values, we used the `adfuller` function from the python `statsmodels` library [83].

### Strain inference

To phase strains, we use a modified version of the allele frequency trajectory clustering algorithm developed by Roodgar *et al*. [36]. While the approach of Roodgar *et al*. was appropriate for their purposes—namely, detecting selective sweeps of linked variants that deviated substantially from the overall background—our clustering scheme is designed to detect only large clusters of SNVs (minimally, greater than 1000 SNVs within a cluster) whose linkage patterns are consistent with perfect linkage on a single haplotype background. Our choice of 1000 SNVs as a cutoff was informed by previous work estimating the typical scale of genetic divergence between strains found in different hosts (See S1 Text section 3.1 for further information). While lineages can and do diverge as a result of diversifying evolution within hosts [34, 46], by imposing the minimum cluster size of 1000 core genome SNVs, we expect largely to exclude cases of within-host diversification. When no cluster of greater than 1000 SNVs was detected, only a single strain was inferred to be present.

### Macroecology

To fit Taylor’s Law (Figure 3A), we used the polyfit function from the python `numpy` library to fit a power law regression between the mean and variance of each strain’s abundance distribution. To obtain the log rescaled Gamma AFD (Figure 3B), each strain’s abundance distribution was first log rescaled, and then normalized to have 0 mean and unit variance. We then binned each strain’s rescaled abundance distribution into 20 evenly spaced bins, and fit the Gamma AFD to the bin-wise mean across strains. To perform the Gamma AFD fit, we adapted code from [1].

## Code availability

All necessary metadata, as well as the source code for the MIDAS metagenomic pipeline, downstream analyses, and figures, are available on GitHub: https://github.com/richardwolff/strain_stability.

## Acknowledgments

We thank Colin Kremer for his crucial comments on this manuscript, Van Savage for early discussions on this work, Jacopo Grilli for code and important discussions, as well as members of the Garud lab for continued feedback. Lastly, we thank two anonymous reviewers for their constructive feedback. This work was supported by the NSF Postdoctoral Research Fellowships in Biology Program under Grant No. 2010885 (W.R.S.). as well as the the Paul Allen Foundation, the UCLA Hellman Fellowship, and the Research Corporation for Science Advancement (N.R.G)

