## Supplementary material for "Ecological Stability Emerges at the Level of Strains in the Human Gut Microbiome": S1 Text

### Supplementary materials

#### Supplementary text

#### Supplementary Figures and Tables

| **Host alias** | **Samples** | **Unique sample days** | **Sampling period (days)** | **Mean samples per day** | **Median days between samples** | **Max days between samples** |
|---|---|---|---|---|---|---|
| *am* | 206 | 193 | 539 | .35 | 1 | 51 |
| *ao* | 74 | 71 | 144 | .49 | 1 | 14 |
| *an* | 63 | 61 | 375 | .16 | 2 | 58 |
| *ae* | 59 | 58 | 375 | .29 | 2 | 41 |

**Table S1:** Host sampling metadata for the four hosts considered in the study. When sample metadata indicated that multiple samples had been collected on the same day, or were technical replicates of one another, samples were merged by concatenating fastq files. Hence, the total number of samples reported (**Samples**) differed from the number of samples analyzed (**Unique sample days**). While gaps between successive successive samples were as long as 58 days (*an*), the median time between samples was either one or two days.

| Species name | Genome length (bp) |
|---|---|
| *Alistipes onderdonkii* | 3,864,643 |
| *Alistipes putredinis* | 2,550,678 |
| *Bacteroides cellulosilyticus* | 6,870,144 |
| *Bacteroides clarus* | 3,746,690 |
| *Bacteroides fragilis* | 5,723,061 |
| *Bacteroides intestinalis* | 6,052,596 |
| *Bacteroides massiliensis* | 4,443,004 |
| *Bacteroides ovatus* | 7,880,760 |
| *Bacteroides stercoris* | 4,009,829 |
| *Bacteroides thetaiotaomicron* | 6,855,195 |
| *Bacteroides uniformis* | 4,719,097 |
| *Bacteroides vulgatus* | 5,163,189 |
| *Bacteroides xylanisolvens* | 5,986,762 |
| *Barnesiella intestinihominis* | 3,433,706 |
| *Bifidobacterium adolescentis* | 2,304,613 |
| *Dialister invisus* | 1,895,960 |
| *Eubacterium eligens* | 2,831,389 |
| *Eubacterium rectale* | 3,344,951 |
| *Faecalibacterium prausnitzii* (57453) | 3,127,383 |
| *Faecalibacterium prausnitzii* (61481) | 3,090,349 |
| *Faecalibacterium prausnitzii* (62201) | 3,321,367 |
| *Parabacteroides distasonis* | 4,887,173 |
| *Parabacteroides merdae* | 4,434,377 |
| *Paraprevotella clara* | 4,187,245 |
| *Phascolarctobacterium sp* | 2,369,100 |
| *Ruminococcus bicirculans* | 2,968,500 |
| *Ruminococcus bromii* | 2,249,085 |
| *Sutterella wadsworthensis* | 2,949,098 |

**Table S2:** Reference genome lengths used by MIDAS in aligning shotgun reads for the purpose of calling SNVs.

| *am* | |
|---|---|
| **Species name** | **Number of strains** |
| *Bacteroides fragilis* | 1 |
| *Bacteroides ovatus* | 2 |
| *Bacteroides stercoris* | 1 |
| *Bacteroides uniformis* | 1 |
| *Bacteroides vulgatus* | 3 |
| *Bacteroides xylanisolvens* | 2 |
| *Barnesiella intestinihominis* | 1 |
| *Eubacterium rectale* | 2 |
| *Faecalibacterium prausnitzii* (57453) | 1 |
| *Phascolarctobacterium sp.* | 1 |
| *Ruminoccocus bicirculans* | 1 |
| *Ruminoccocus bromii* | 1 |

**Table S3:** Number of strains inferred for each species analyzed in host *am*.

| *ao* | |
|---|---|
| **Species name** | **Number of strains** |
| *Bacteroides fragilis* | 1 |
| *Bacteroides ovatus* | 2 |
| *Bacteroides uniformis* | 1 |
| *Bacteroides xylanisolvens* | 2 |
| *Bifidobacterium adolescentis* | 1 |
| *Dialister invisus* | 1 |
| *Eubacterium eligens* | 1 |
| *Eubacterium rectale* | 1 |
| *Faecalibacterium prausnitzii* (61481) | 2 |
| *Faecalibacterium prausnitzii* (62201) | 1 |
| *Parabacteroides distasonis* | 1 |
| *Ruminoccocus bicirculans* | 1 |

**Table S4:** Number of strains inferred for each species analyzed in host *ao*.

| *an* | |
|---|---|
| **Species name** | **Number of strains** |
| *Alistipes putredinis* | 1 |
| *Bacteroides clarus* | 1 |
| *Bacteroides massiliensis* | 1 |
| *Bacteroides uniformis* | 2 |
| *Bacteroides vulgatus* | 3 |
| *Bacteroides xylanisolvens* | 1 |
| *Barnesiella intestinihominis* | 1 |
| *Eubacterium rectale* | 2 |
| *Paraprevotella clara* | 1 |
| *Ruminoccocus bromii* | 1 |
| *Sutterella wadsworthensis* | 2 |

**Table S5:** Number of strains inferred for each species analyzed in host *an*.

| *ae* | |
|---|---|
| **Species name** | **Number of strains** |
| *Alistipes onderdonkii* | 2 |
| *Alistipes putredinis* | 1 |
| *Bacteroides cellulosilyticus* | 1 |
| *Bacteroides massiliensis* | 2 |
| *Bacteroides ovatus* | 2 |
| *Bacteroides thetaiotaomicron* | 1 |
| *Bacteroides uniformis* | 1 |
| *Bacteroides vulgatus* | 1 |
| *Eubacterium rectale* | 2 |

**Table S6:** Number of strains inferred for each species analyzed in host *ae*.

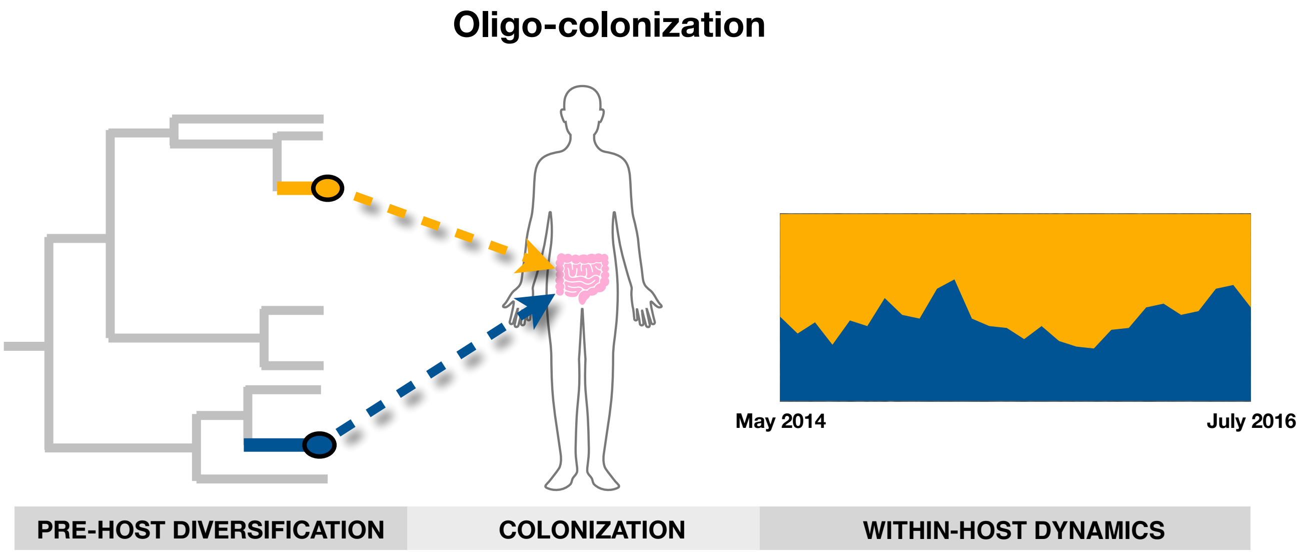

**Figure S1:** A range of human gut microbiome studies have shown that only a small handful of strains belonging to a species are present within a host at any one time, a phenomenon known as oligo-colonization [5, 8, 9, 10, 11, 14]. Strains diversify before colonizing the host, typically accumulating $\mathcal{O}(10^3) - \mathcal{O}(10^4)$ mutations in their shared core genome. In this study, we explore the within-host dynamics of oligo-colonization both in cases where a single strain belonging to a species is present and when multiple strains are present. We find that a large majority of strains stably colonize hosts for periods of months to years, while a smaller percentage show highly non-stationary dynamics. For further discussion of oligo-colonization, Supplementary section 3.1.

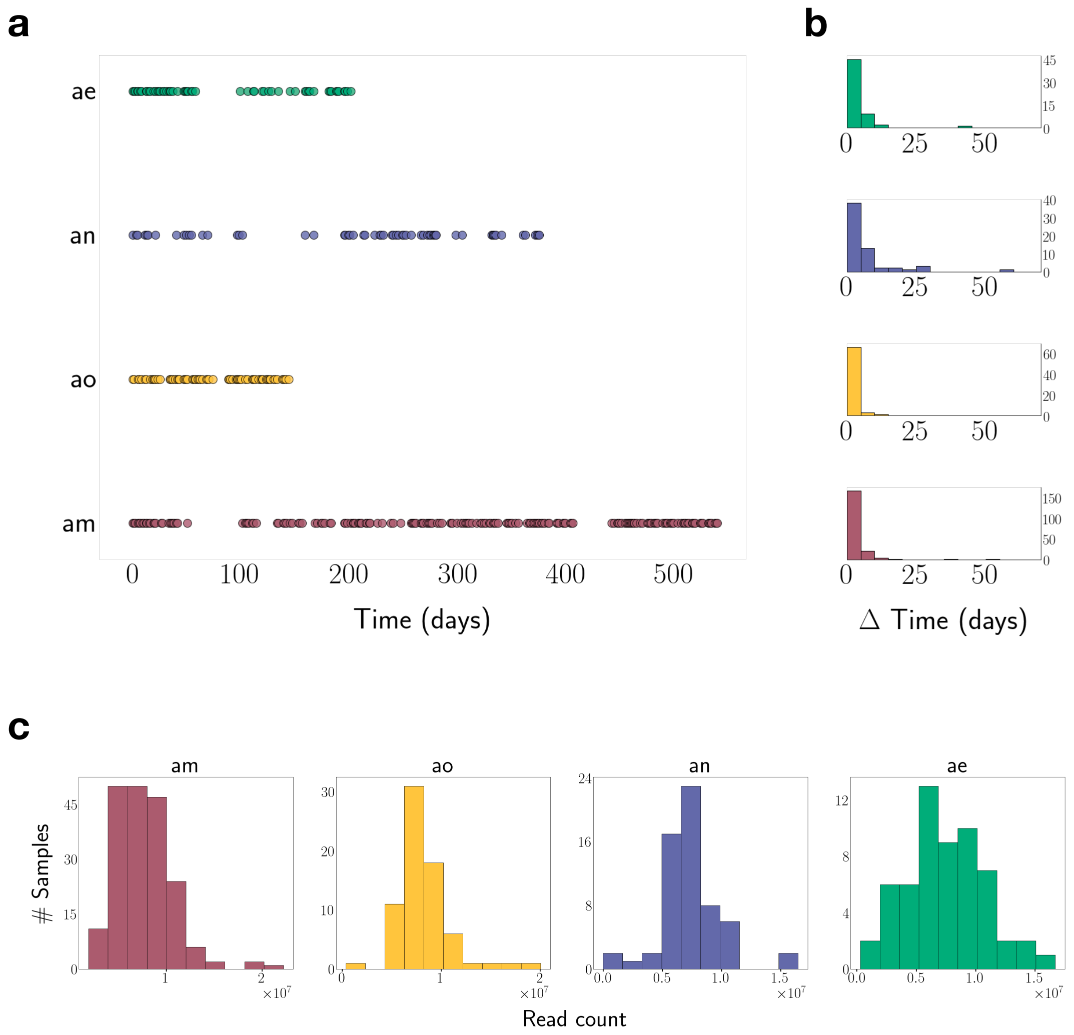

**Figure S2: a.** Four BIO-ML stool donors (aliases: *am*, *ao*, *an*, and *ao*) were longitudinally sampled more than 50 times. Circles indicate that shotgun metagenomic sequencing data was collected at that timepoint. **b.** For each host, we plot the distribution of the number of days between consecutive samples. While gaps between successive successive samples were as long as 58 days, the median time between samples was either one or two days for each host (Supplementary Table 1). **b.** Distribution of the number of shotgun reads per sample, by host. The mean number of reads per sample in each host were am: 7.5m, ao: 7.6m, an: 7.3m, and ae: 8.3m.

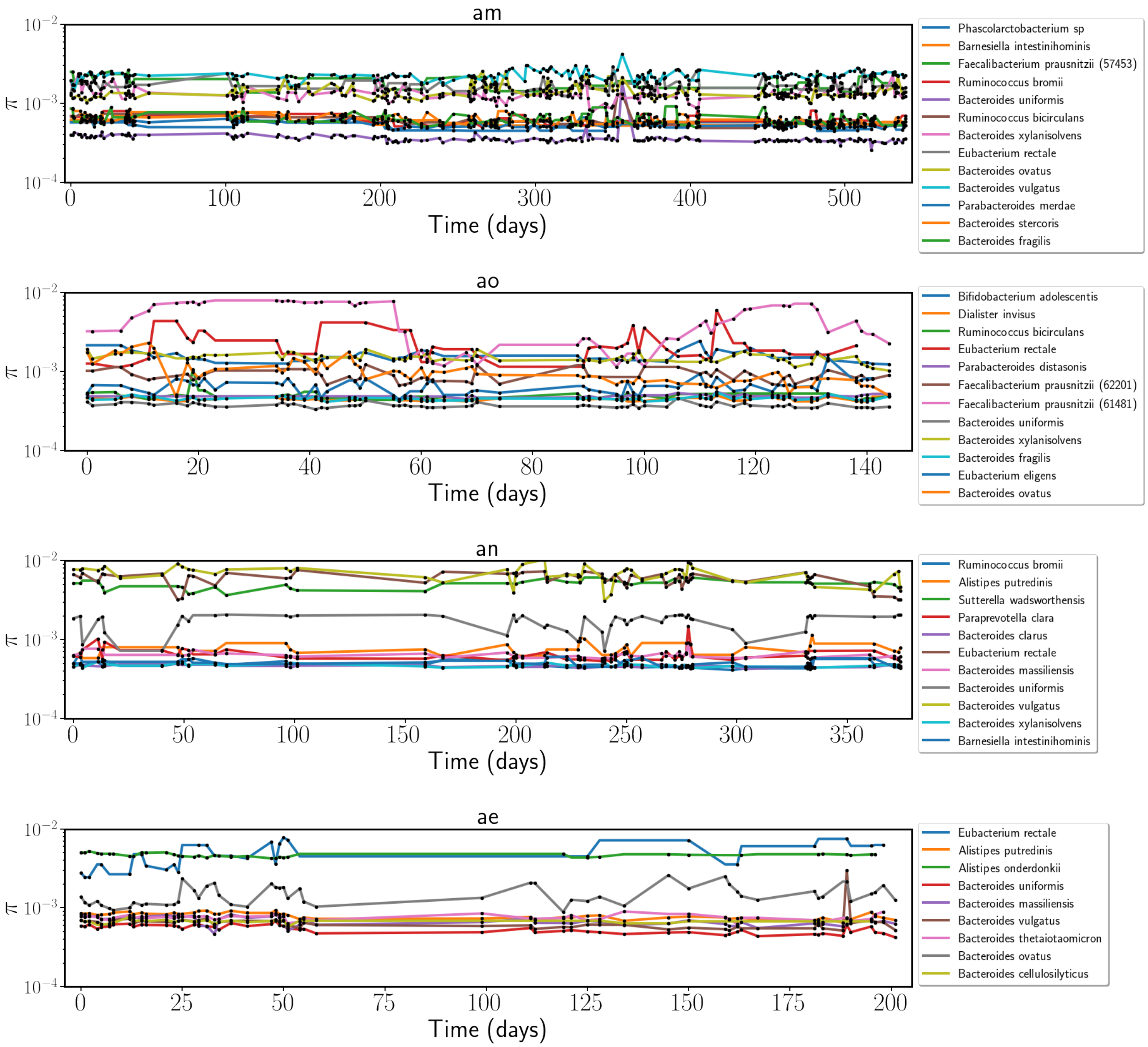

**Figure S3:** Nucleotide diversity $\pi$ for each species/host combination.

.

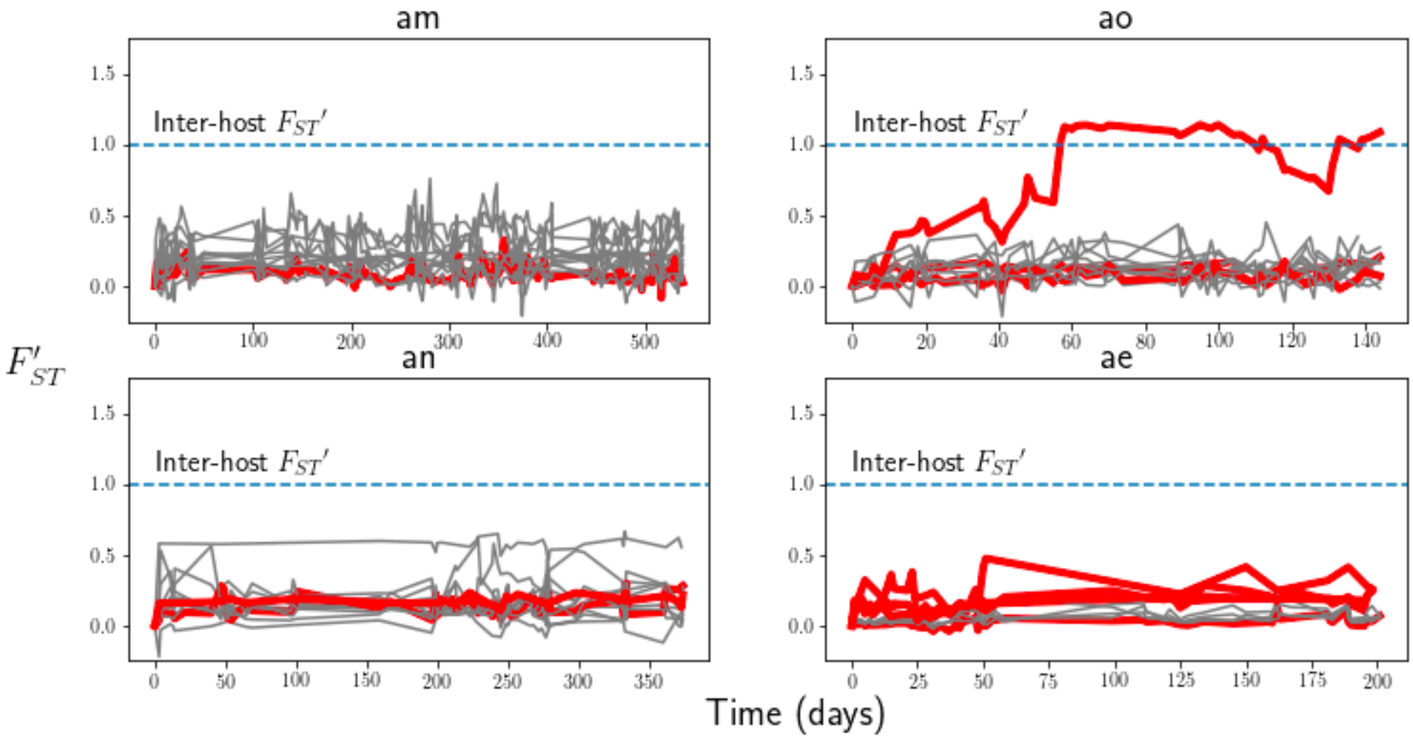

**Figure S4:** $F_{ST}'$ values plotted for each species in each host, with species failing the ADF test of stationarity highlighted in red.

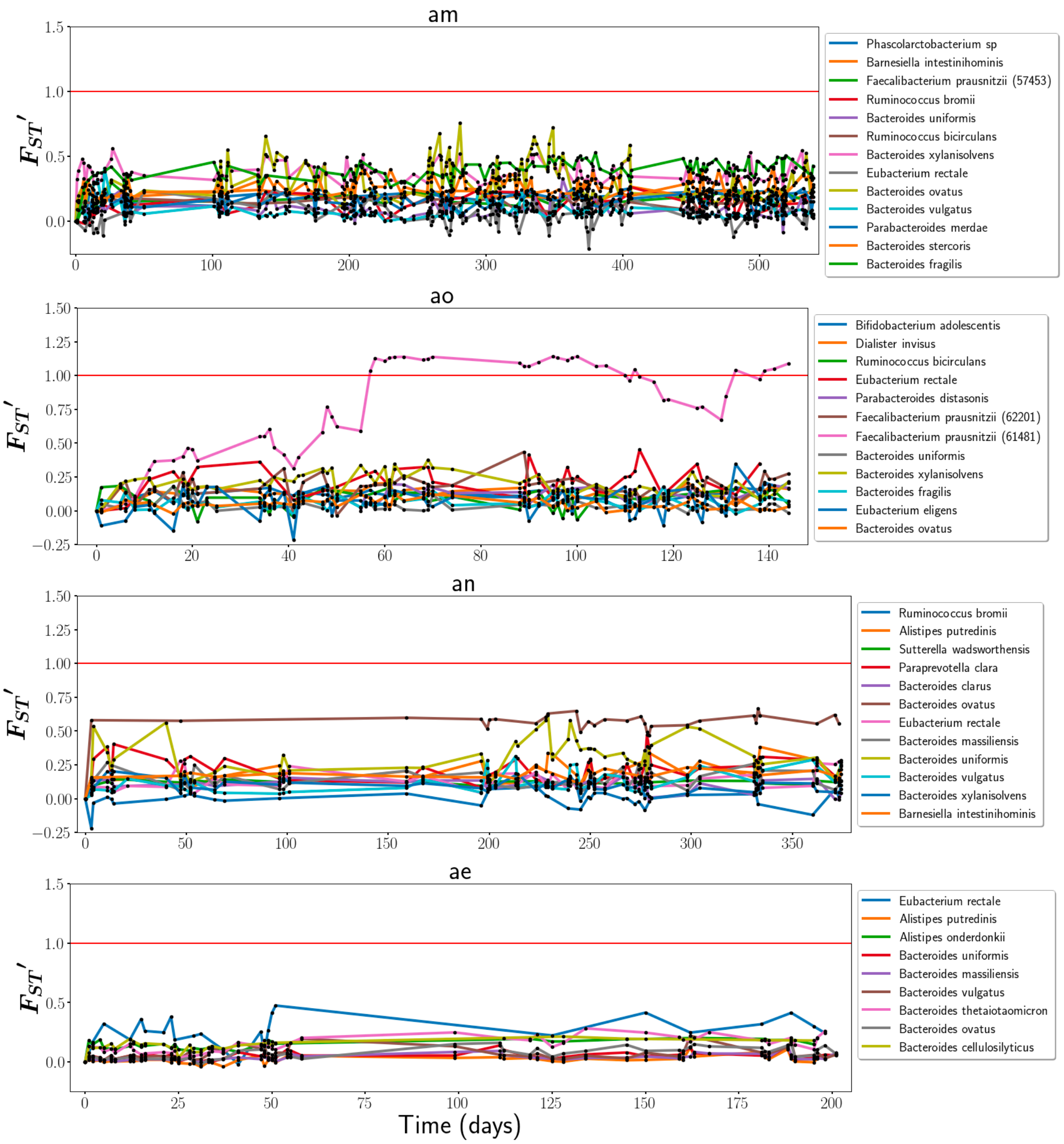

**Figure S5:** $F_{ST}'$ values plotted for each species in each host. For details on how $F_{ST}'$ was calculated, see Materials and Methods of Main Text.

### Supplementary Text

#### 1 Data

Publicly available shotgun metagenomes from the Broad Institute-OpenBiome Microbiome Library (BIO-ML) were downloaded from the Sequence Read Archive (BioProject PRJNA544527). A total of 402 samples drawn from four individuals (aliases *am*, *ao*, *an*, and *ae*) were downloaded, with 206 samples coming from *am*, 74 from *ao*, 63 from *an*, and 59 from *ae* (Table S1). All four hosts were healthy adults between the ages of 21 and 37, of whom three were male and one female. The full metadata concerning these hosts reported by BIO-ML is available as Supplementary File S1.

The four hosts analyzed in this paper were sampled near daily for between 144 days (*ao*) and 539 days (*am*). While gaps between successive successive samples were as long as 58 days (*an*), the median time between samples was either one or two days (Table S1).

When sample metadata indicated that multiple samples had been collected on the same day, or were technical replicates of one another, samples were merged by concatenating fastq files before running subsequent metagenomic processing scripts (Supplementary section 2). Thus, all samples taken from a single host used in this study had a unique date, and were temporally ordered.

Additionally, to calculate between-host $F_{ST}$, we used data from a previously collated panel [8] of healthy human subjects of 250 North American subjects sequenced by the Human Microbiome Project [19, 20]. Using this much broader database of samples, we estimated levels of inter-host genetic diversity.

#### 2 Metagenomic pipeline

We used the MIDAS software package to align metagenomic reads to reference genomes, estimate species abundances, and call copy number variation (CNV) and single nucleotide variant (SNV) content [3]. MIDAS's reference database is made up of 31,007 high quality bacterial genomes, clustered into 5,952 species groups based on a 96.5% sequence identity across 30 universal marker genes (MIDAS reference database version 1.2). This marker gene average nucleotide identity (ANI) threshold was chosen by the original authors of MIDAS to ensure a 95% genome-wide ANI species definition. Throughout this manuscript, strains belonging to a species are much more closely related than this 95% threshold—typically, between 99% and 99.9% ANI.

All three basic modules in the MIDAS pipeline (estimating species abundance, calling CNV content, and calling SNV content) consist of two steps. First, each sample is processed individually to ascertain the relative abundance of all species, as well as the CNV and SNV content for those species passing user-specified thresholds in that sample (for details on threshold choices, see Supplementary section 2.2). And second, individual samples belonging to the same host are integrated to identify patterns of longitudinal variation within hosts.

##### *2.1 Species abundance*

MIDAS first estimates species abundances by aligning reads to 15 universal, single-copy marker gene sequences, which are known for each species in the MIDAS database. The average read coverage across these single-copy marker genes is then taken to be the abundance of the species in the sample. In our subsequent analyses, we excluded species which were considered present in fewer than 25 samples (see Supplementary section 5).

##### 2.2 *Building reference panels*

When calling CNV and SNV content, MIDAS relies on a sample-specific reference panel to ensure that reads are only mapped to species which are truly present. We considered a species present if it had an average marker gene coverage $\geq 3$ in that sample. While many more species are present at lower abundances, and are detected as such by MIDAS, SNV and CNV content cannot be confidently determined in these cases due to their relatively poor coverage [8].

To conservatively guard against spurious detection of intra-species genetic diversity, a "blacklist" of genes defined in [8] which are known to be commonly shared between species were excluded from further analysis were excluded from all downstream analyses.

Lastly, to ameliorate read stealing and donating, we included all species which were present at any timepoint within a host in the reference panel for all samples in that host. By doing so, we keep the set of pan-genomes to which reads are aligned consistent across samples. By taking this precaution, we ensure that the species a specific read is mapped to will remain the same regardless of the sample in which this read originates.

##### 2.3 *CNV content*

CNV content is determined by aligning reads to pan-genomes present in the MIDAS database for each species present in our data. MIDAS defines a pan-genome as the set of unique genes identified across all genomes available in the MIDAS reference database belonging to a species.

Reads were aligned to the host-specific reference panel using default MIDAS settings (local alignment, MAPID $\geq$ 94.0%, READQ $\geq$ 20, and ALN_COV $\geq$ 0.75), and the average coverage was estimated by dividing the total number of reads mapping to the gene cluster by its total target size.

MIDAS determines the copy number $c$ of a gene by dividing the average coverage of the gene by the average coverage of the species at its universal, single-copy markers (i.e. the species abundance). Using these copy number estimates, we then defined the core genome for each species in each host as the set of genes with copy number $c \geq 0.3$ in $90\%$ of samples belonging to that host. Because we defined the core genome in a host-specific manner, this core represents the set of genes which are shared between all strains *within a host*. Species detected in multiple hosts, therefore, may have somewhat different gene content in the "core" genome.

##### 2.4 *SNV content*

MIDAS calls SNVs with reference to a single representative genome per species group (specified in the MIDAS database, and chosen so as to have minimal marker gene distance to all other isolate genomes associated with the same species). The total length, in base pairs, of the representative genomes used by MIDAS for SNV calling purposes for species analyzed in this work is shown in (**Table S2**).

To ascertain SNVs, we aligned reads to the species included for a given sample's reference panel using default MIDAS mapping parameters: global alignment, MAPID $\geq$ 94.0%, READQ $\geq$ 20, ALN_COV $\geq$ 0.75, and MAPQ $\geq$ 20. After alignment, we further excluded from analysis species that did not have a mean genome-wide coverage $\geq 5$ and non-zero coverage at $\geq 40\%$ of reference sites within a sample.

### 3 Detecing strains

#### *3.1 Oligo-colonization*

A wide range of human microbiome studies have established that gut microbial species harbor genetic diversity within individual hosts [6, 2, 20]. Species are in fact frequently composed of multiple strains, and which are much more distantly genetically related than lineages which diverged subsequent to colonization [10, 14, 23]. When levels of recombination between such strains are sufficiently low, the clonal descendants of the initial colonizers may persist within hosts as genetically distinguishable subpopulations [24]. Interestingly, a number of studies have demonstrated that this kind of multiple colonization is subject to some degree of ecological constraint: only a small handful of strains (typically between one and four) are ever observed within a host at any one time, a phenomenon dubbed "oligo-colonization" (see Figure S1) [5, 8, 11, 9]. The general ecological and evolutionary mechanisms enabling a small number of strains to colonize a host and rise to high frequency, but preventing a large number of exogenous strains from doing the same, are not known as yet. Indeed, the factors governing strain coexistence may vary greatly between species. For instance, only a single strain of the common gut commensal *Bacteroides fragilis* is ever found within a host, potentially due to unusually potent inter-strain competition in this species mediated by the Type-VI secretion system [9]. However, other species, like *Bacteroides vulgatus*, frequently harbor several strains [5, 23]. Whether such fine-scale diversity is maintained due to the partitioning of metabolic niches, by spatial segregation, or due to chaotic or neutral dynamics is an active area of research [17, 18]. In this work, we examine the dynamics of oligo-colonization through time.

Genetically diverged strains of a species residing in different hosts will typically differ from one another at $\mathcal{O}(10^3) - \mathcal{O}(10^4)$ sites in their shared core genome [2, 4, 8, 23]. Likewise, when multiple strains of the same species inhabit the same host, they also tend to differ from one another at a similar number of sites—put differently, each strain will share a more recent common ancestor with a strain found in a different host than with its other co-colonizing strains [23]. On the other hand, even under conservative estimates of the per basepair mutation rate and host lifetime, lineages sharing a common ancestor within the host are expected to harbor orders of magnitude fewer segregating mutations [8]. We will leverage this separation of scales in the number of sites expected to segregate between lineages sharing a common ancestor within the host, and those belonging to separately colonizing strains, to infer the presence of strains.

Resolving the number, relative abundances, and genotypes of co-colonizing strains from a single shotgun metagenomic sample is a difficult inverse problem. Because short read shotgun sequencing destroys evidence of physical linkage information over , methods of resolving strains within a sample rely on statistically deconvolving the distribution of allele frequencies to find areas of high density [15, 16]. However, due to finite and variable metagenomic read coverage across the genome, even perfectly linked alleles residing on a single strain's genetic background may have quite different observed allele frequencies due solely to sampling noise. However, several recent studies [5, 13, 14, 24, 26] have shown that strain inference can be significantly improved when many samples have been collected longitudinally from a host, as alleles linked on a single strain's genetic background will have clustered allele frequency trajectories. Strain inference algorithms designed specifically for such longitudinal data can leverage these temporal correlations between trajectories as a potent additional source of information when inferring true patterns of linkage within species.

##### 3.2 *Strain inference*

In our study, we modified the allele frequency trajectory clustering algorithm developed by Roodgar *et al.* [5] to specifically detect large clusters of SNVs ( $>1000$ SNVs) in a species' core genome which strongly support the presence of multiple strains. While the approach of Roodgar *et al.* was appropriate for their purposes—namely, detecting selective sweeps of linked variants that deviated substantially from the overall background—the original clustering UPGMA-based clustering procedure was not sensitive enough to reliably detect low frequency strains in our dataset and thus required minor modifications. Our choice of 1000 SNVs as a cutoff was informed by the previously mentioned (Supplementary section 3.1) lower bound on the typical number of SNVs segregating between strains found in different hosts ($\mathcal{O}(10^3)$ sites). While lineages can and do diverge due to evolution within hosts [6, 7], by imposing the minimum cluster size of 1000 core genome SNVs, we expect largely to exclude cases of within-host diversification.

To cluster SNV trajectories into strains, we first calculated the distance metric defined in Roodgar *et al.* (which we review here) between every pair of polymorphic SNVs with sufficient coverage. To be considered, we required a site to have depth of coverage $D \geq 10$ and some level of polymorphism ($0 < f < 1$, where $f$ is the frequency of the reference allele) in at least 25% of timepoints. By imposing these filters, we excluded sites that were not truly polymorphic, and/or for which true allele frequencies could not be accurately estimated.

This metric weights differences in allele frequency trajectories between sites by their joint sequencing depth, such that small differences in frequency at high coverage sites contribute proportionally more than equally small differences at low coverage sites. In practice, this ameliorate the of sampling noise. For a pair of SNVs $i$ and $j$, the distance between their observed allele frequency trajectories $\hat{f}_i$ and $\hat{f}_j$ is:

$$d(\hat{f}_i, \hat{f}_j) = \frac{1}{T} \sum_{t=1}^{T} \frac{2(D_{it} + D_{jt})(\hat{f}_{it} - \hat{f}_{jt})^2}{(\hat{f}_{it} + \hat{f}_{jt})(1 - \hat{f}_{it} + 1 - \hat{f}_{jt})} \tag{1}$$

where $T$ is the number of sampling timepoints, $\hat{f}_{it}$ and $\hat{f}_{jt}$ are the allele frequencies of $i$ and $j$ at a timepoint $t$, and $D_{it}$ and $D_{jt}$ are the depths of sequencing coverage at timepoint $t$.

At any biallelic site, there are two possible choices of allelic reference state. While the choice is *a priori* arbitrary, it will impact our ability to infer strain structure, as the metric (Equation 1) relies on both sites having the same relative polarization. To account for differences in relative polarization, we define a pair of metrics:

$$\begin{aligned} d_{ij}^{+} &= d(\hat{f}_i, \hat{f}_j) \\ d_{ij}^{-} &= d(\hat{f}_i, 1 - \hat{f}_j) \end{aligned} \tag{2}$$

and take the distance between $i$ and $j$ to be the minimum of the two polarizations:

$$d_{ij} = \min(d_{ij}^{+}, d_{ij}^{-}). \tag{3}$$

Still following [5], we calculated $d_{ij}$ between every pair of SNVs, forming a distance matrix. However, having calculated this SNV distance matrix, we took a different approach to clustering the SNVs into strains. Specifically, while [5] applied UPGMA clustering to the matrix of SNV distances, we implemented a greedy, network-based clustering algorithm. Our algorithm allows us to extract only large clusters of SNVs which are likely in perfect linkage with one another.

We considered each SNV a node in a network, and formed an edge between any pair of nodes $(i, j)$ if $d(\hat{f}_i, \hat{f}_j) < d^*$, where $d^*$ is a critical distance indicating likely true linkage. We used the empirically derived threshold maximum distance criterion of $d^* = 3.5$, established in [5], to identify pairs of SNVs that were likely linked to one another on a haplotype background. In essence, this network connects SNVs whose allele frequency trajectories indicate that they likely rely on the same haplotype background.

Next, we calculated the degree $|D_i|$ (i.e. number of edges) for all nodes in the network. We identified the node $k$ with the highest degree $|D_k|$ in the network, and if $|D_k| > 1000$, we assigned $k$ and all nodes connected to it to be a strain. To ensure that no outlier SNVs not truly linked to this cluster were assigned to it, we removed all nodes from the cluster which were connected to less than $25\%$ of other nodes. Lastly, we removed all nodes assigned to this cluster from the network. We then repeated this procedure until no clusters of greater than 1000 nodes remained. If there were no clusters of more than 1000 SNVs initially, we inferred that only a single strain of the species was present.

When two strains are present, all SNVs which segregating between the strains will ultimately be polarized in the same way, and so only a single cluster of SNVs will be detected (for ease of visualization, we have shown both the detected cluster and in the Figure 1 of the Main text, as well as in the Supplemental strain figure files). However, when three strains are present, each pair of strains will have some number of sites segregating between them. Consequently, our strain deconvolution algorithm will detect three clusters of SNVs when three strains are present. In the two cases in which we detected three strains, we determined the correct relative polarizations of these clusters with respect to one another by summing all different possible polarizations, and looked for the choice of polarization that yielded a sum which was close to one at all timepoints, as the sum of the relative strain frequencies should be precisely one at each timepoint. In both cases, there was a single choice of polarization that resulted in the sum of strain trajectories remaining near one at all timepoints (green trajectories), and these polarizations were therefore chosen when determining the true frequencies of the strains (Figure S6).

Recently, Zheng *et al* [25] employed a high-throughput single-cell barcoding scheme to analyze strain dynamics in samples collected from host *am* of the BIO-ML project. As in our work, Zheng *et al* uncovered three dominant strains of *B. vulgatus* (a fourth strain was recovered from their co-assembly, but was detected only at a single timepoint, had very low genome completeness (6%), and was of “low quality"). Though collected at a far coarser sampling interval (seven timepoints, versus the 193 timepoints available using shotgun metagenomic samples), there is a striking visual concordance in the dynamics of the three dominant strains of *B. vulgatus* between our inference and that of Zheng *et al*. Interestingly, in the original BIO-ML publication [6], isolates of *B. vulgatus* collected from *am* indicated the presence of only two strains. That our inference, which relies on relatively easily obtainable shotgun metagenomic sequencing data processed through publicly available software in a matter of days, reveals similar strain dynamics at far greater temporal resolution as the state-of-the-art sequencing approach of Zheng *et al* in this case underscores the power of densely temporally sampled, high coverage shotgun sequencing data to recover patterns of intraspecific genetic variation in the microbiome.

#### 4 SLM

To simulate the SLM, we used the Euler-Maruyama method:

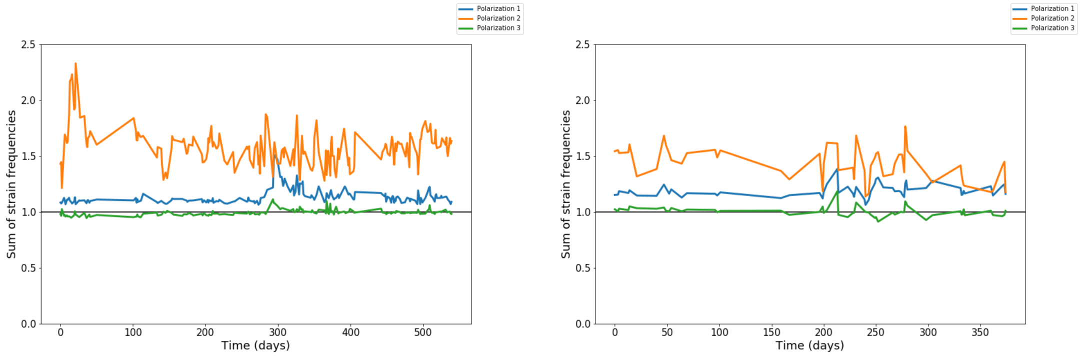

**Figure S6:** In two samples, *am* (left) and *an* (right), three strains of *B. vulgatus* were detected. In order to assess the correct relative polarizations of the three strains, we summed all different possible polarizations of the detected clusters. In both cases, the green polarization shown in the figure above was approximately equal to one at all timepoints, and so this polarization was chosen when determining the true frequencies of the strains.

$$X(t+\delta t) = X(t) + \frac{x(t)}{\tau_i}\left(1 - \frac{x(t)}{K}\right)\delta t + \sqrt{\frac{\sigma}{\tau}}x(t)Z_t\sqrt{\delta t} \tag{4}$$

where $Z_t$ is a standard normal random variable. In simulations, we set $\delta t = \frac{1}{1000}$.

The SLM associated with population $i$ depends on three parameters: $K_i$, $\sigma_i$, and $\tau_i$. $K_i$ and $\sigma_i$ are not fit, but rather are determined directly from the mean and variance of the actual time series using the formulae:

$$\sigma_i = \frac{2}{\frac{\langle x_i \rangle^2}{\sigma^2_{x_i}} - 1}, \quad K_i = \frac{\langle x_i \rangle}{1 - \frac{\sigma_i}{2}} \tag{5}$$

where $\langle x_i \rangle$ is the mean abundance of the population and $\sigma^2_{x_i}$ is its variance. The parameter $\tau_i$ was held constant ($\tau_i = 1$) for all strains to avoid overfitting.

For each time series of strain abundances, we fit the $K_i$ and $\sigma_i$ parameters using the first third of timepoints.

#### 5 Goodness of fit test

To assess the predictive capacity of the SLM, we devised a goodness of fit test. The test seeks to determine not only whether the true dynamics of a strain's abundance match the behavior expected under the SLM, but also whether these dynamics can be predicted from a subset of initial points. Heuristically, the test determines whether the full sequence of transitions between timepoints statistically resembles the sequence of transitions expected were the data truly generated by the SLM.

First, we calculated the total abundance of the strain in question, observed at timepoints $t_0, t_1, t_2, ..., t_T$, to be $x(t_0), x(t_1), x(t_2), ..., x(t_T)$. The total abundance of a strain is it relative abundance within the community at large—that is, its relative frequency within the species (determined using the strain inference algorithm described in the preceding section), multiplied by the relative frequency of the species within the community.

For each strain, the SLM is "trained" on the first 33% of data points (i.e. $x(t_0), x(t_1), ..., x(t_{T/3})$), with the model parameters $K_i$ and $\sigma_i$ determined as described in the main text directly from the mean and variance in abundance across these timepoints.

The fundamental approach employed to test the fit of the SLM is to estimate an empirical distribution under the model of possible abundances at $t_{i+1}$, given the known initial abundance $x(t_i)$, and then observing where the true $x(t_{i+1})$ lies. To do so, we simulated the model 1000 times between each pair of subsequent timepoints $t_i$ and $t_{i+1}$ $(i \in [(T/3+1), (T/3+2), ..., T])$, starting with initial abundance $x(t_i)$. Next, to asses where the true $x(t_{i+1})$ lay relative to our simulations, we subdivided this distribution into $M$ equally likely quantiles (that is, $M$ intervals of the real line each having equal probability under the empirical distribution of simulations), determined which quantile $x(t_{i+1})$ fell in, and denoted this value $q_i$.

Under the null hypothesis, $x(t_{i+1})$ should have equal probability of lying in each quantile, precisely because the quantiles are non-overlapping areas of equal probability under the SLM. Therefore, $\{q_i\}$, the set of quantiles determined in this manner between all pairs of consecutive timepoints, is expected to follow a uniform distribution on $[1, 2, ..., M]$ if the true trajectory was generated by the SLM. To assess whether this was in fact the case, we performed a $\chi^2$ test with $M$ degrees of freedom. We rejected the null at a significance level of 5%. We do not perform multiple hypothesis testing here, as we only make a single comparison (pass/fail of SLM).

In determining $M$, we followed the convention for $\chi^2$ tests that the expected number of observations in each bin (here, $[1, M]$) should exceed 5. As there are a total of $T$ data points, $M$ was chosen so that $\frac{T}{M} > 5$. Additionally, to ensure that our $\chi^2$ test had sufficient power, we only considered trajectories which had $T > 25$ timepoints [1].

#### 6 Taylor's Law

Taylor's Law, a power law scaling between the mean and variance of the observed abundances of populations within a community across either space or time, is an extremely general phenomena observed in a wide variety of ecosystems [28, 27, 30, 31, 32]. The specific scaling exponent observed is affected both by true biological variability in the underlying populations' abundances and by noise introduced during the sampling process. While many different ecological models can produce Taylor's Law with a wide variety of scaling exponents [29, 33], sampling noise, by itself, is expected to induce a Taylor's Law $\alpha$ exponent of one. As an example, consider a community where population abundances are governed by Poisson fluctuations introduced solely by sampling noise:

$$P(x_i = k) = \frac{\lambda^k e^{-\lambda}}{k!} \tag{6}$$

where $\lambda$ is a composite rate parameter, the product of the total strength of observation (e.g. total number of individuals censused in a community) and the focal population's relative abundance. Because the mean of the Poisson distribution equals its variance (both are equal to $\lambda$, in this case), we should observe a Taylor's Law with $\alpha = 1$ (i.e. $\sigma_{x_i}^2 \propto \langle x_i \rangle$, trivially).

Real metagenomic data, however, is compositional—when comparing samples, we do not assess abundance directly, but rather observe the relative abundances of populations. If we assume that there are a finite number reads, we can model the situation in which the variation in observed

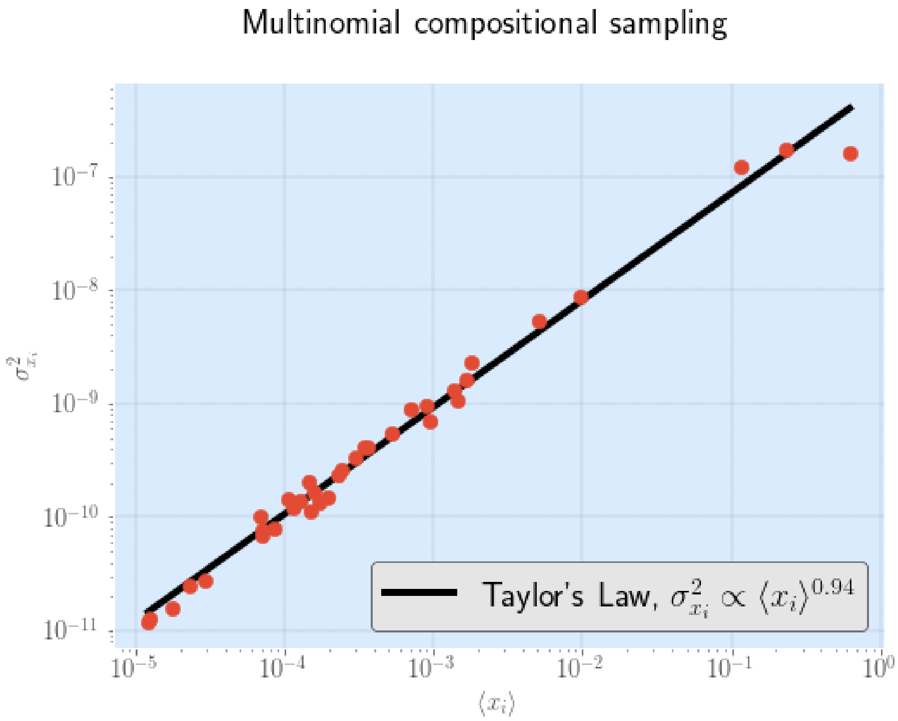

**Figure S7:** $\alpha$ is approximately equal to 1 under multinomial sampling

frequencies is determined solely by sampling noise by, first, repeatedly sampling total abundances from a multinomial distribution:

$$P(x_1, x_2, ..., x_N; N, f_1, f_2, ..., f_N) = \frac{S!}{x_1! \, x_2! \, ... x_N!} f_1^{x_1} f_2^{x_2} ... f_N^{x_N} \tag{7}$$

where $S$ is the total number of reads, $N$ total number of populations present, and $\bar{\mathbf{f}}$ is vector of underlying true population abundances (here, drawn from a lognormal distribution), and $T$ is the number of timepoints. Following this sampling step, we can then characterize fluctuations in relative abundance by dividing all observed abundances by the total read depth, $S$. Doing so, we observe a Taylor's Law scaling with $\alpha \approx 1$, as expected (Figure S7).

Moreover, even if the total read count $S$ is allowed to vary between samples, $\alpha$ still remains approximately equal to 1. Specifically, if $S$ is itself drawn from a Poisson distribution—as would be expected if sampling intensity was roughly equal between samples—and observed population frequencies are affected only by sampling, we continue to observe an $\alpha \approx 1$ (Figure S8).

Though the relative abundances of all strains we observe in this study are certainly affected by sampling noise, and (as we have seen) a Taylor's Law scaling can emerge due solely to compositional sampling, compositionality by itself cannot explain the scaling observed in the data. Specifically, the specific exponent we observe here ($\alpha = 1.8$, Main text Figure 3) is highly unlikely to be primarily driven by sampling effects, whether compositional or not, but rather reflects true underlying biological variability in the relative abundance of the populations.

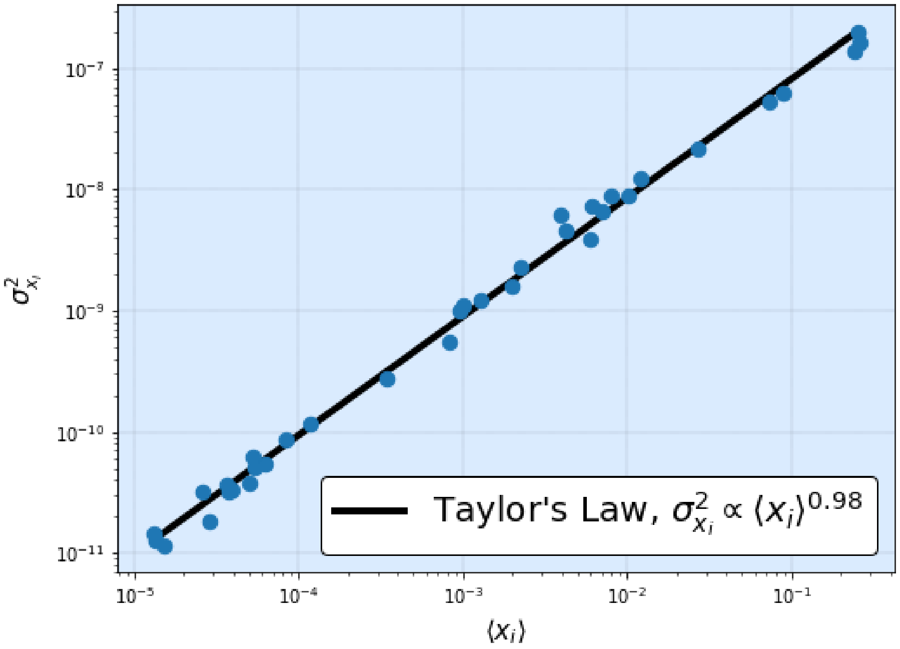

**Figure S8:** $\alpha$ is approximately equal to 1, even when coverage varies between samples.
