## Supplementary material for "Ecological Stability Emerges at the Level of Strains in the Human Gut Microbiome": S2 Text

$F_{ST}$', strain frequency, and strain abundance dynamics plots for all species analyzed, for host *am.* These plots are analogous to Main Text Figure 1. When only a single strain was detected, only $F_{ST}$' and strain abundance dynamics plots, but no strain frequency plot, is included.

*Table of Contents*


| | |
|---|---|
| *Bacteroides fragilis A* | *1* |
| *Bacteroides ovatus A* | *2* |
| *Bacteroides ovatus B* | *3* |
| *Bacteroides stercoris A* | *4* |
| *Bacteroides uniformis A* | *5* |
| *Bacteroides vulgatus A* | *6* |
| *Bacteroides vulgatus B* | *7* |
| *Bacteroides vulgatus C* | *8* |
| *Bacteroides xylanisolvens A* | *9* |
| *Bacteroides xylanisolvens B* | *10* |
| *Barnesiella intenstinihominis A* | *11* |
| *Eubacterium rectale A* | *12* |
| *Eubacterium rectale B* | *13* |
| *Faecalibacterium prausnitzii (57453) A* | *14* |
| *Phascolarctobacterium sp. A* | *15* |
| *Parabacteroides merdae A* | *16* |
| *Ruminoccocus bicirculans A* | *17* |
| *Ruminoccocus bromii A* | *18* |

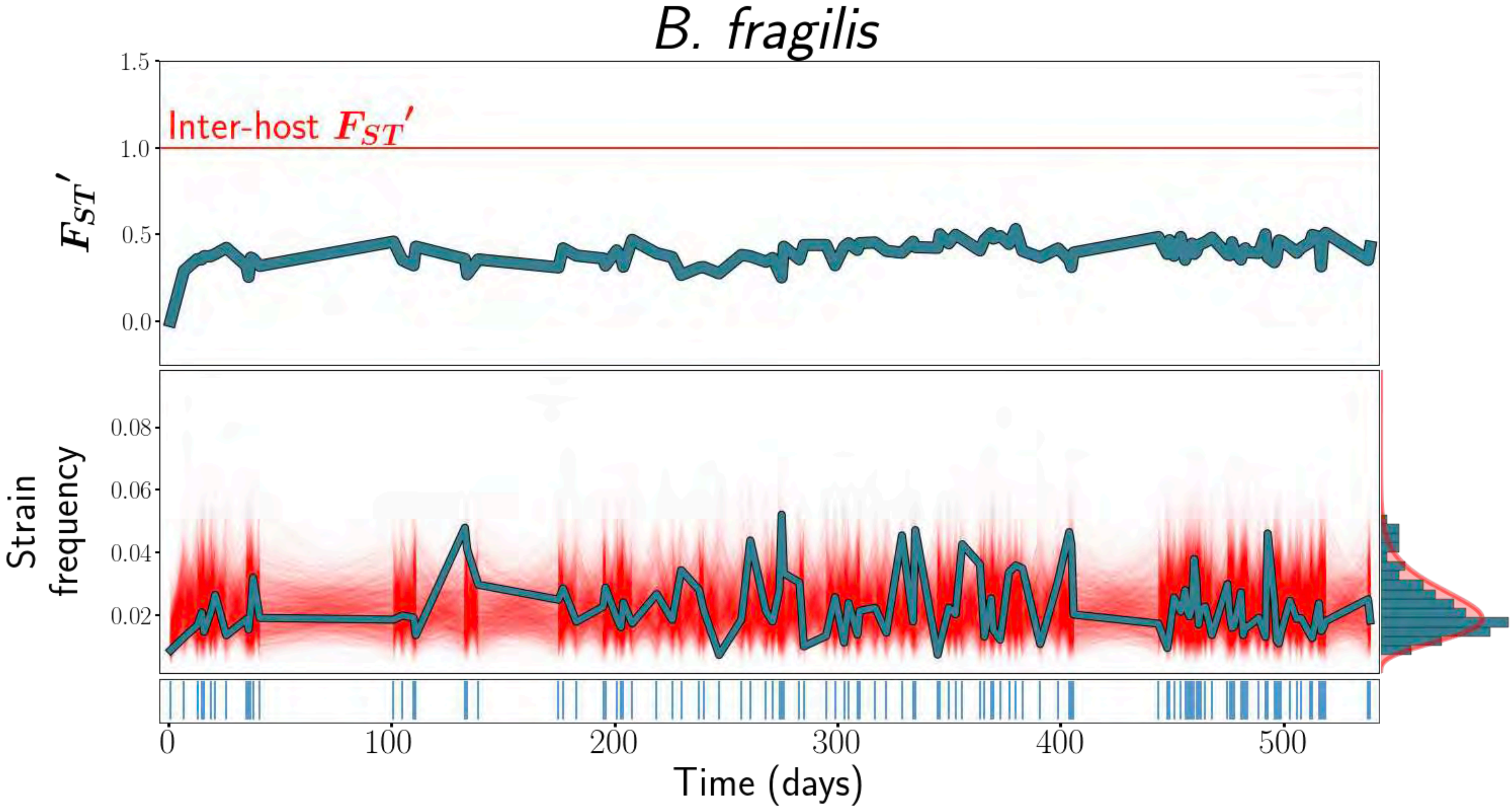
B. fragilis
Inter-host $F_{ST}'$
$F_{ST}'$
1.5
1.0
0.5
0.0
Strain frequency
0.08
0.06
0.04
0.02
0
100
200
300
400
500
Time (days)

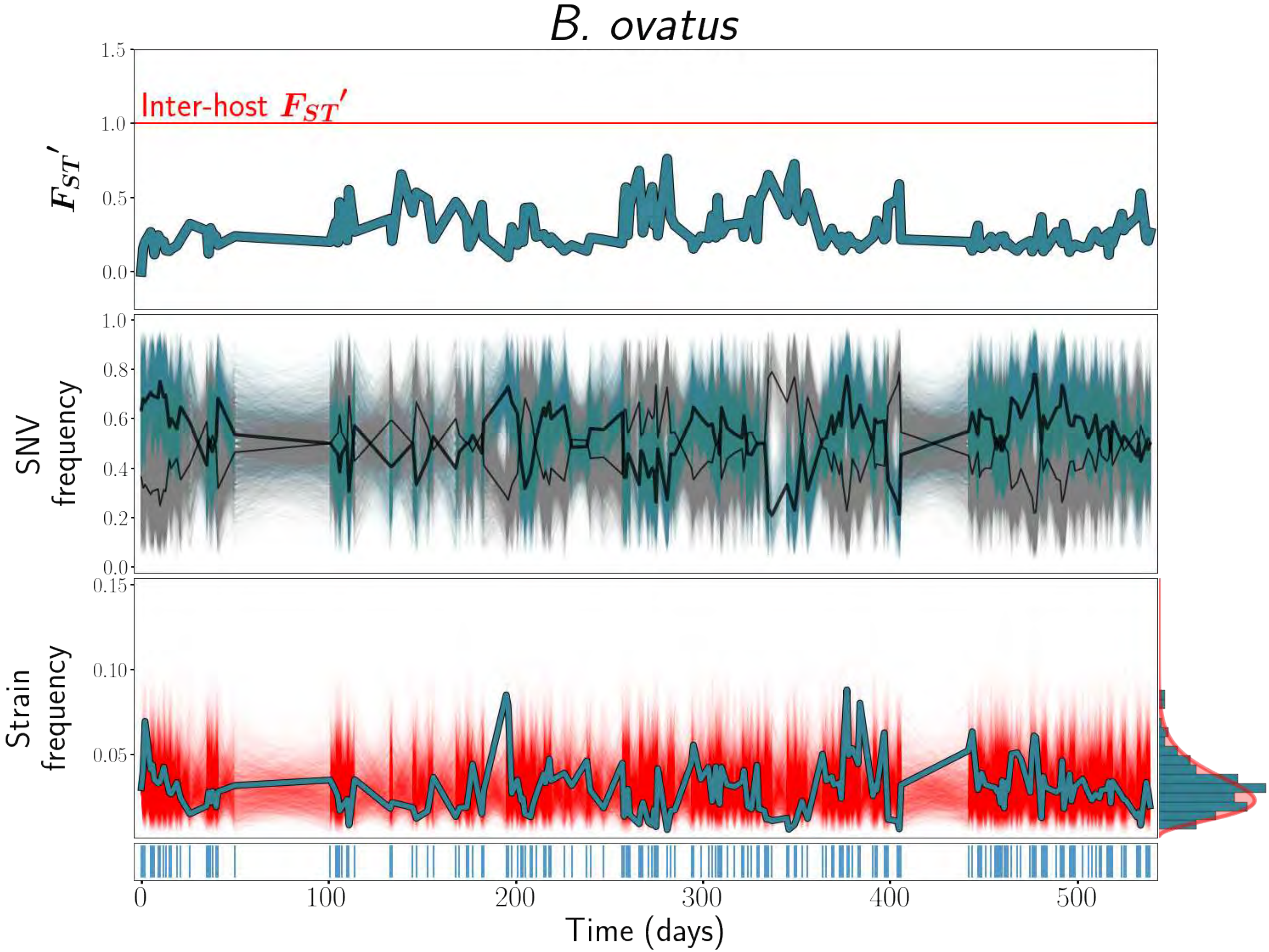

B. ovatus
Inter-host $F_{ST}'$
$F_{ST}'$
1.5
1.0
0.5
0.0
SNV frequency
1.0
0.8
0.6
0.4
0.2
0.0
Strain frequency
0.15
0.10
0.05
0
100
200
300
400
500
Time (days)

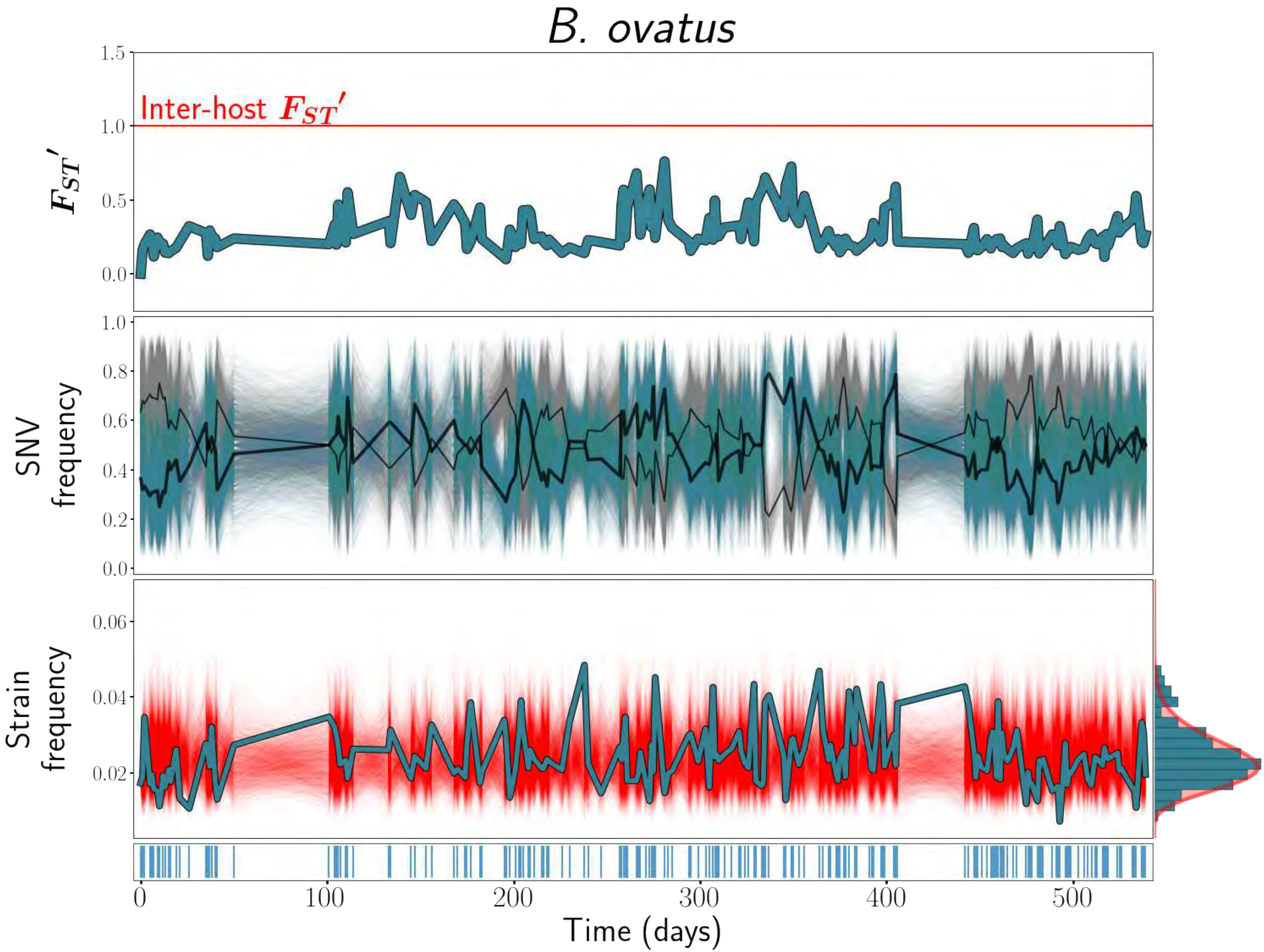
B. ovatus
Inter-host $F_{ST}'$
$F_{ST}'$
1.5
1.0
0.5
0.0
SNV frequency
1.0
0.8
0.6
0.4
0.2
0.0
Strain frequency
0.06
0.04
0.02
0
100
200
300
400
500
Time (days)

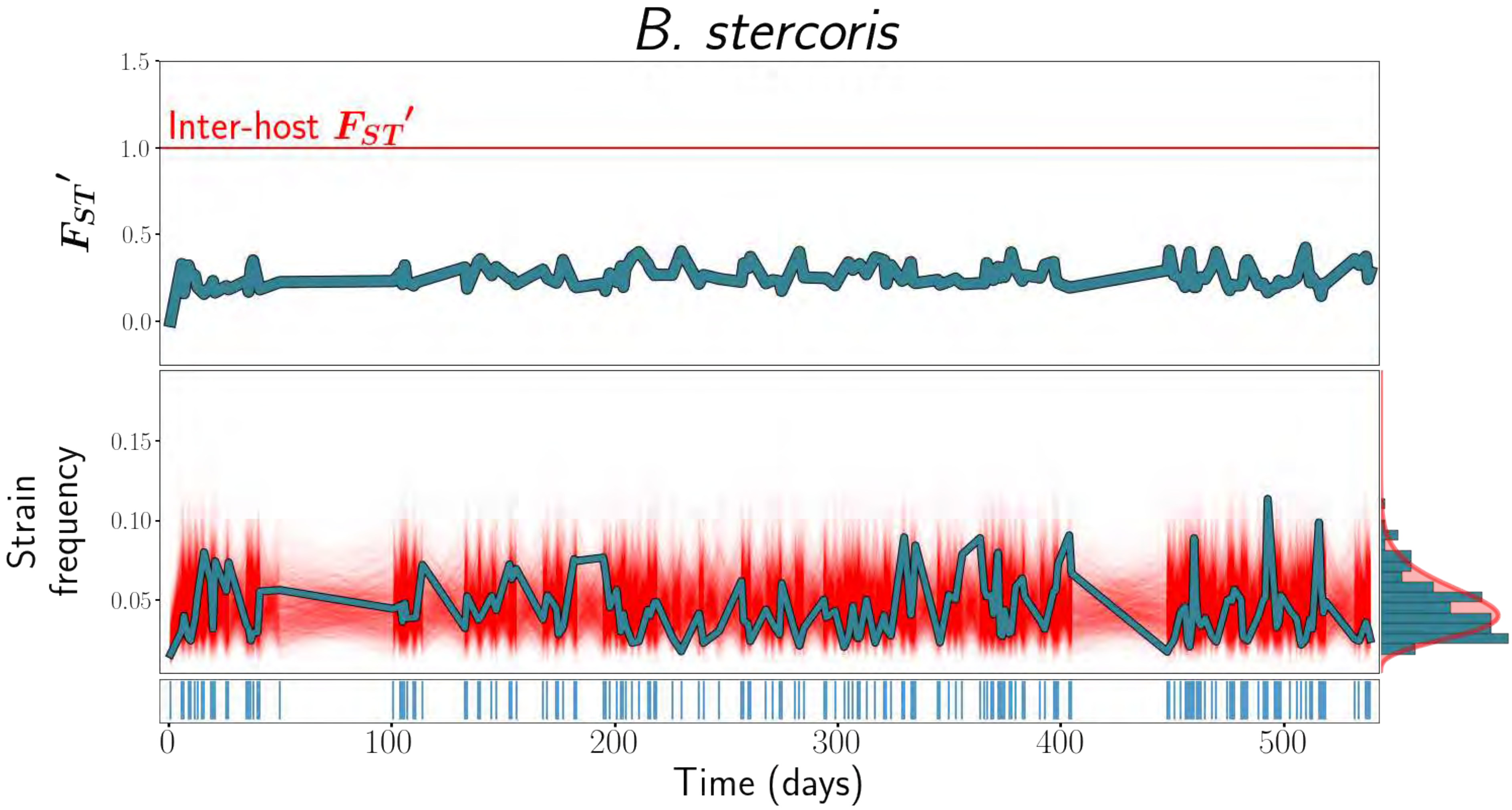

B. stercoris
Inter-host $F_{ST}'$
$F_{ST}'$
Strain frequency
Time (days)
1.5
1.0
0.5
0.0
0.15
0.10
0.05
0
100
200
300
400
500

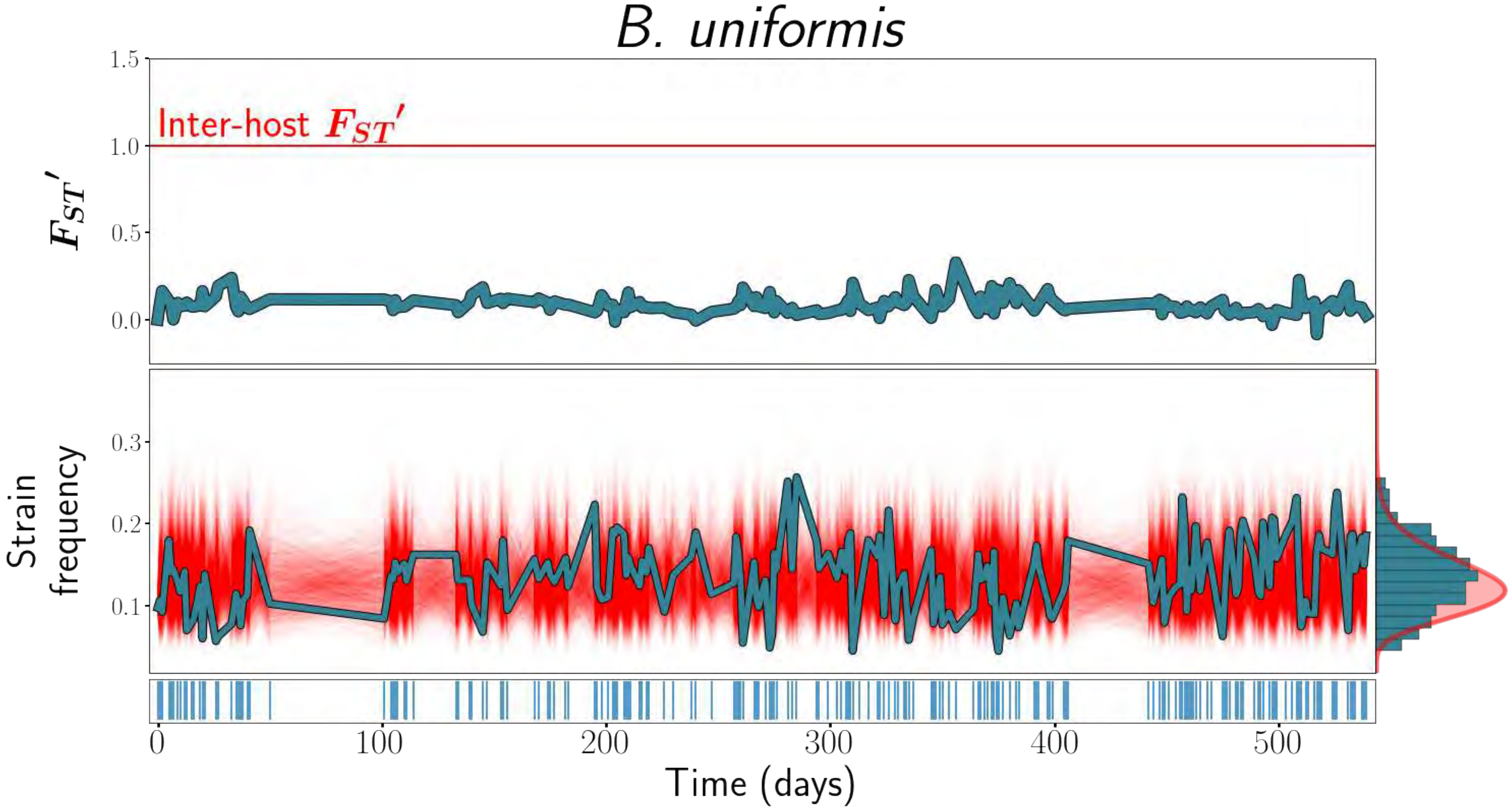

B. uniformis
1.5
Inter-host $F_{ST}'$
1.0
$F_{ST}'$
0.5
0.0
0.3
Strain frequency
0.2
0.1
0
100
200
300
400
500
Time (days)

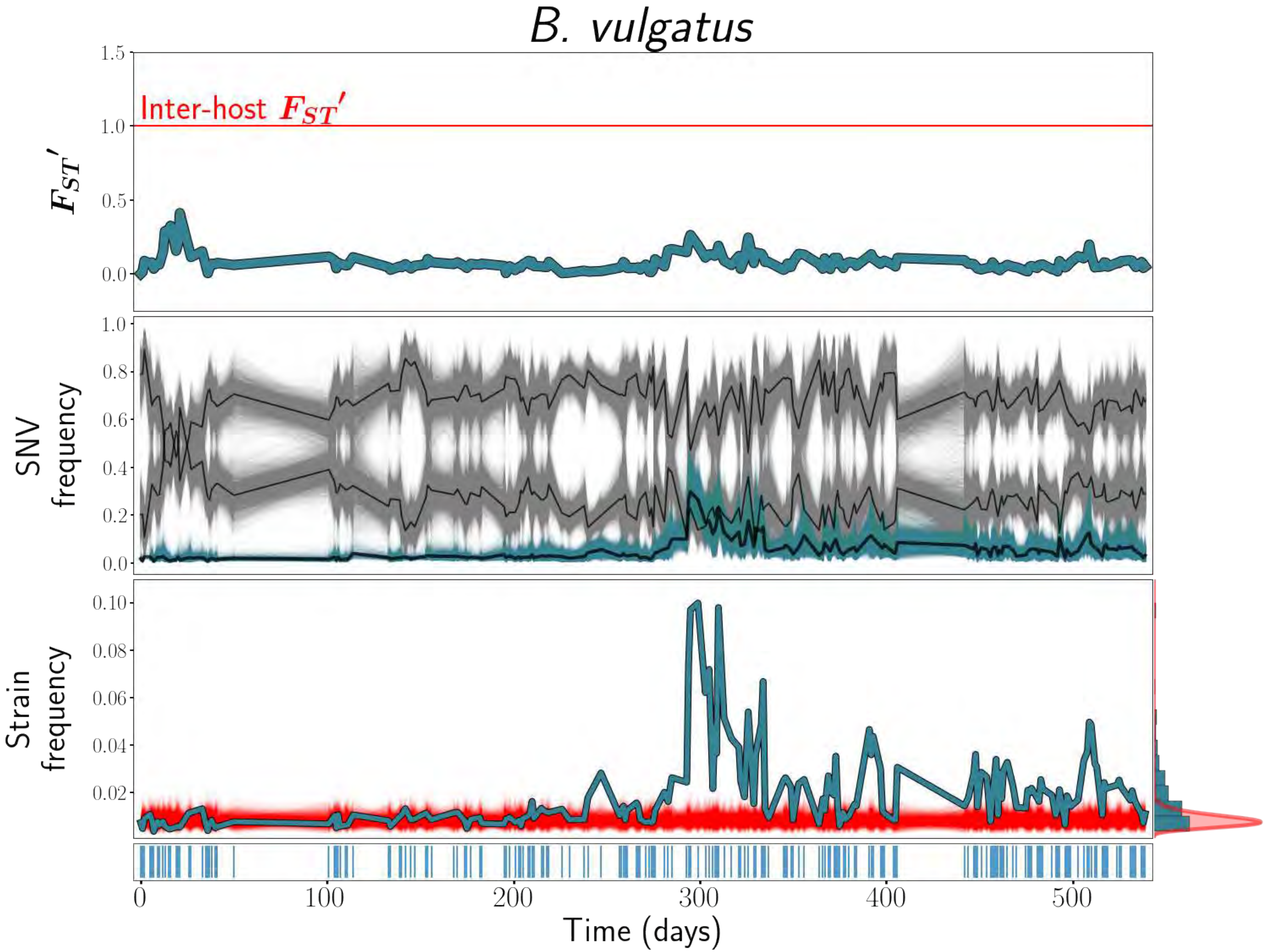
B. vulgatus
1.5
Inter-host $F_{ST}'$
1.0
$F_{ST}'$
0.5
0.0
1.0
0.8
0.6
SNV frequency
0.4
0.2
0.0
0.10
0.08
Strain frequency
0.06
0.04
0.02
0
100
200
300
400
500
Time (days)

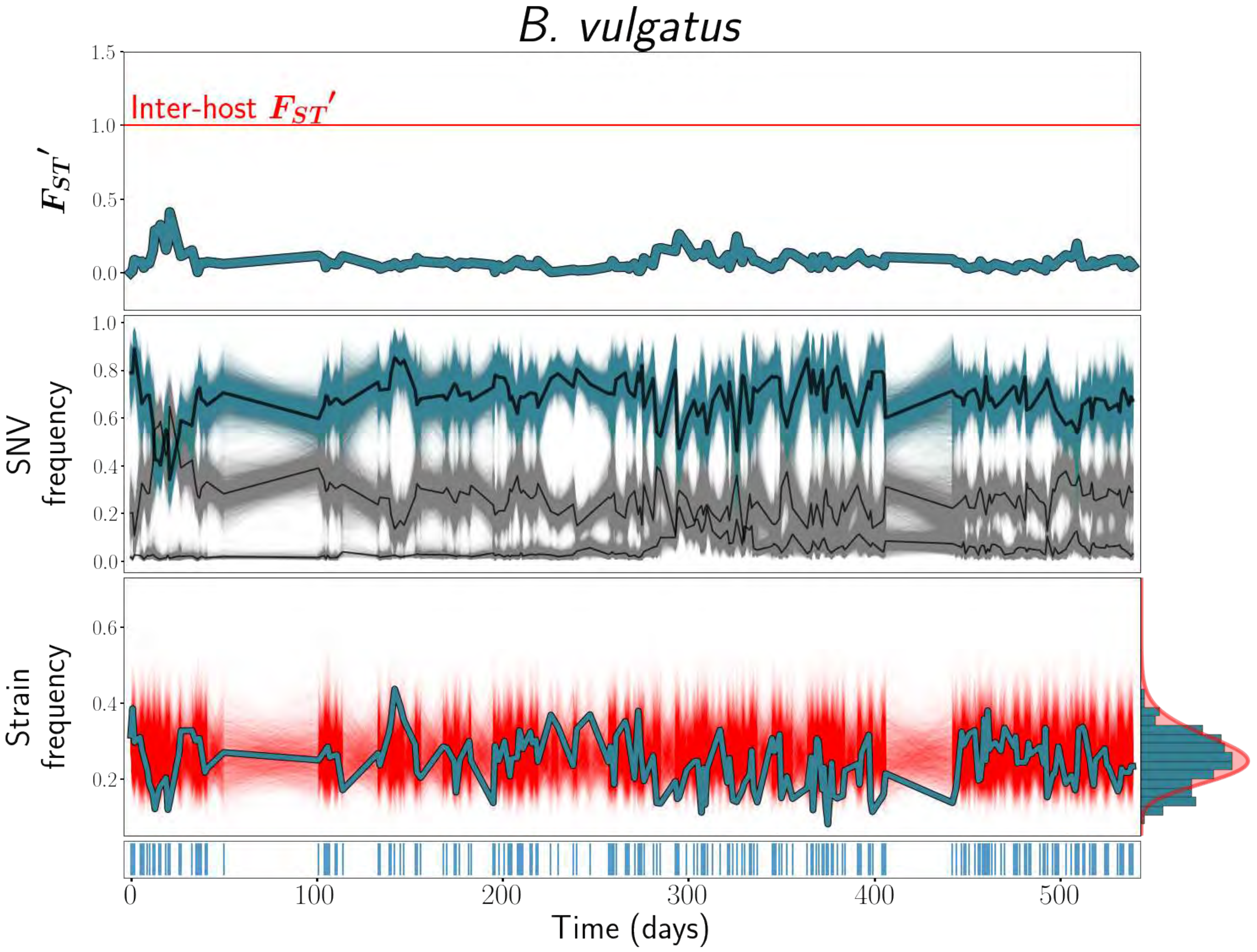
B. vulgatus
Inter-host $F_{ST}'$
$F_{ST}'$
SNV frequency
Strain frequency
Time (days)
1.5
1.0
0.5
0.0
0.8
0.6
0.4
0.2
0
100
200
300
400
500

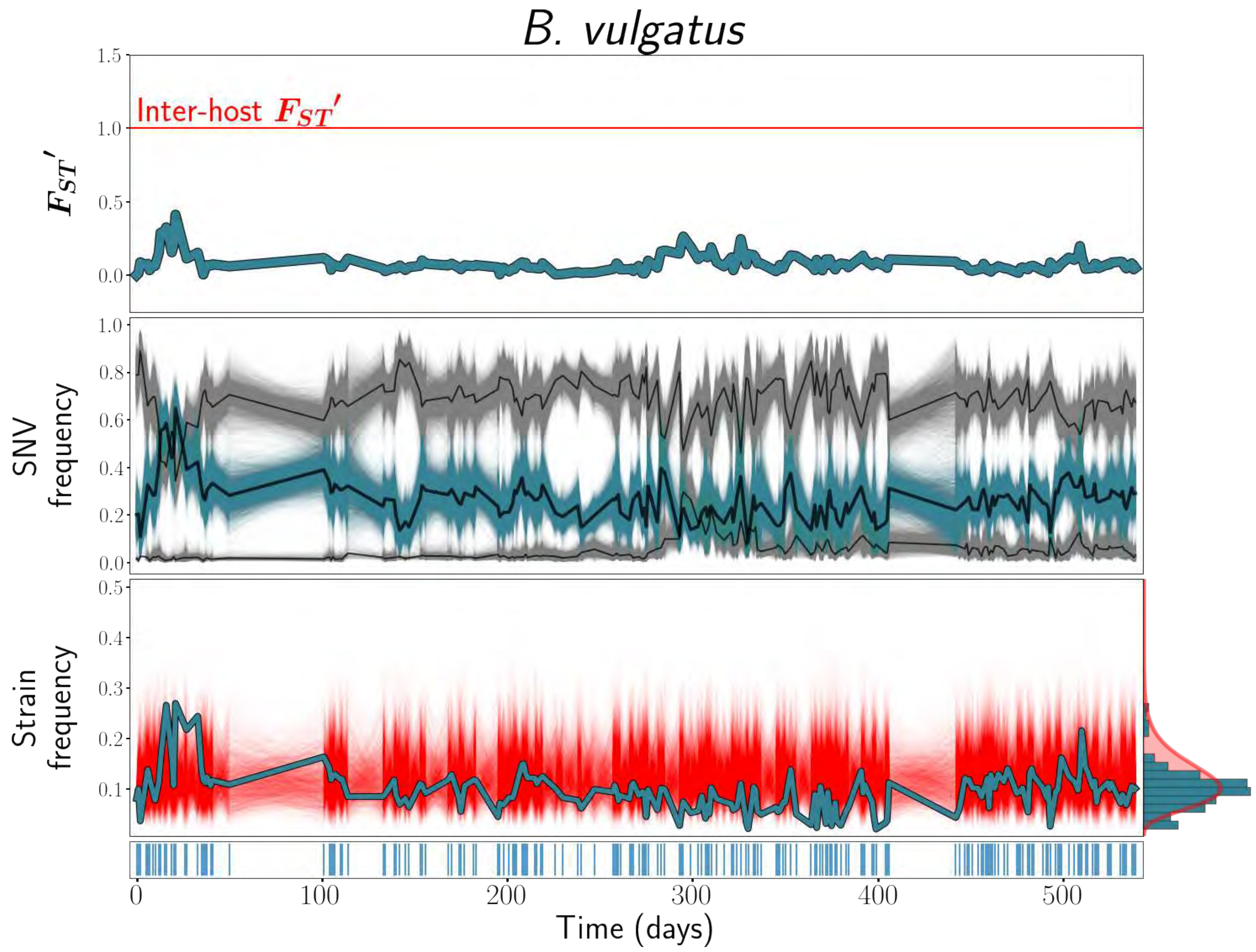
B. vulgatus
Inter-host $F_{ST}'$
$F_{ST}'$
SNV frequency
Strain frequency
Time (days)
1.5
1.0
0.5
0.0
0.8
0.6
0.4
0.2
0.3
0.1
0
100
200
300
400
500

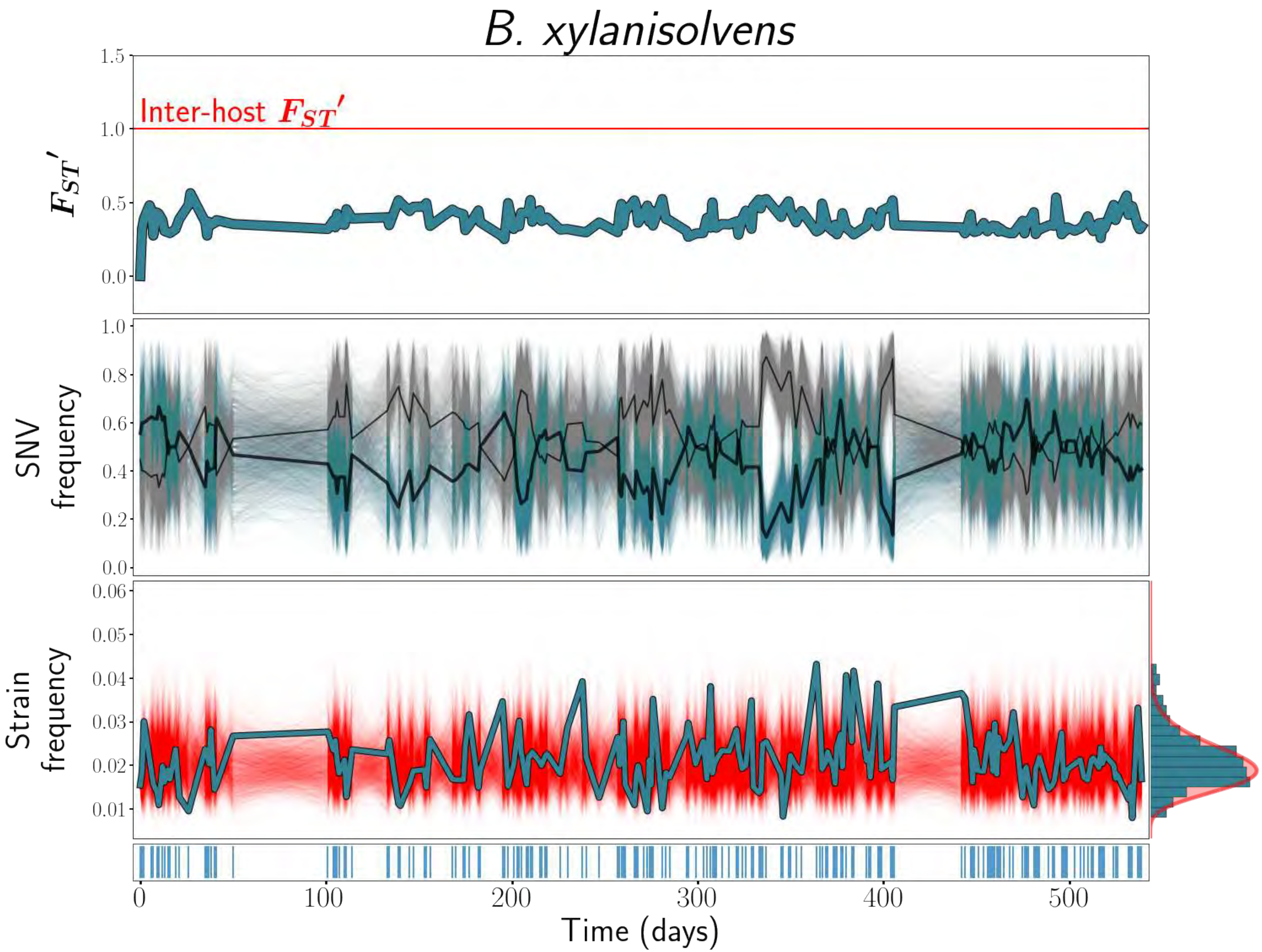
B. xylanisolvens
Inter-host $F_{ST}'$
$F_{ST}'$
1.5
1.0
0.5
0.0
SNV frequency
1.0
0.8
0.6
0.4
0.2
0.0
Strain frequency
0.06
0.05
0.04
0.03
0.02
0.01
0
100
200
300
400
500
Time (days)

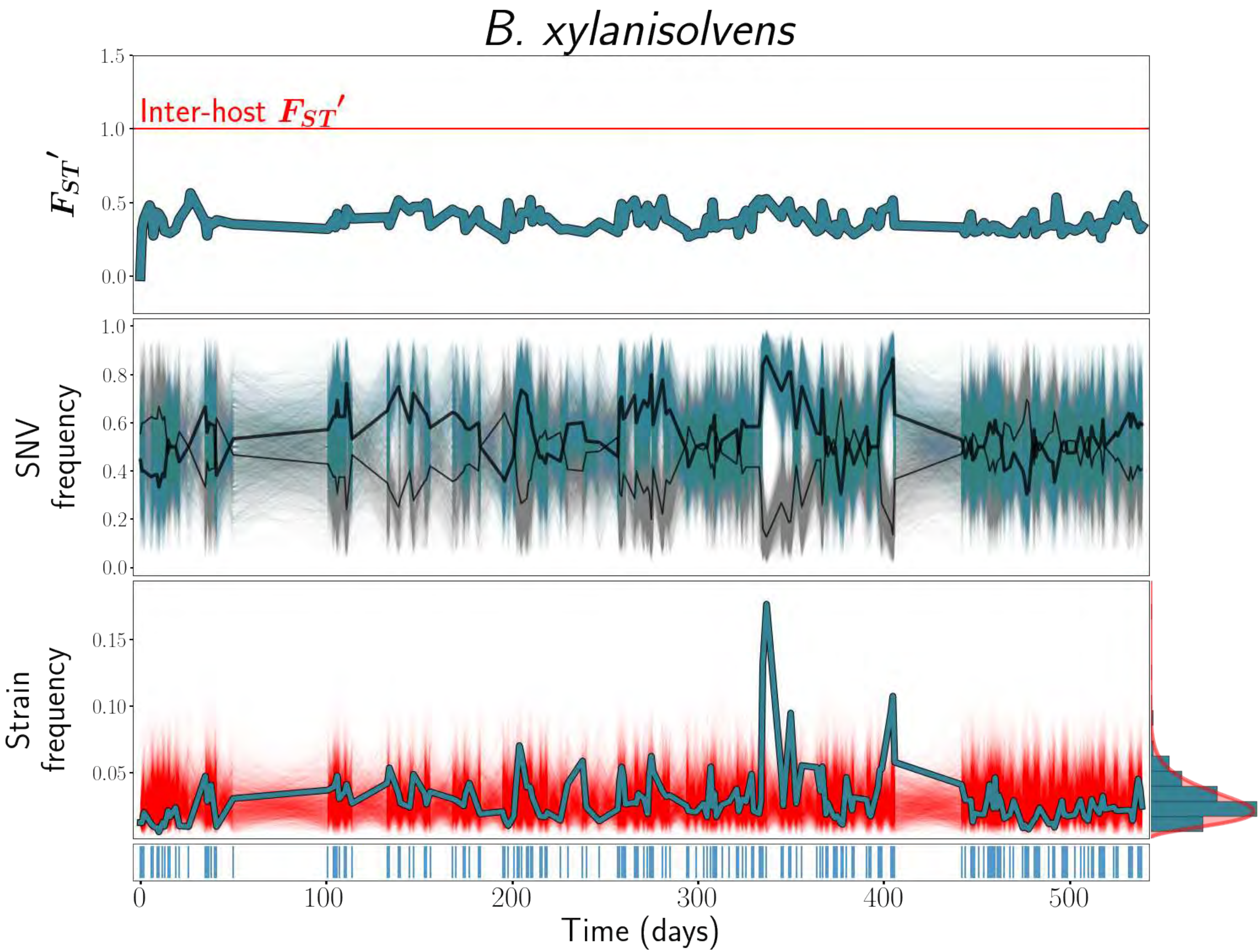
B. xylanisolvens
Inter-host $F_{ST}'$
$F_{ST}'$
1.5
1.0
0.5
0.0
SNV frequency
1.0
0.8
0.6
0.4
0.2
0.0
Strain frequency
0.15
0.10
0.05
0
100
200
300
400
500
Time (days)

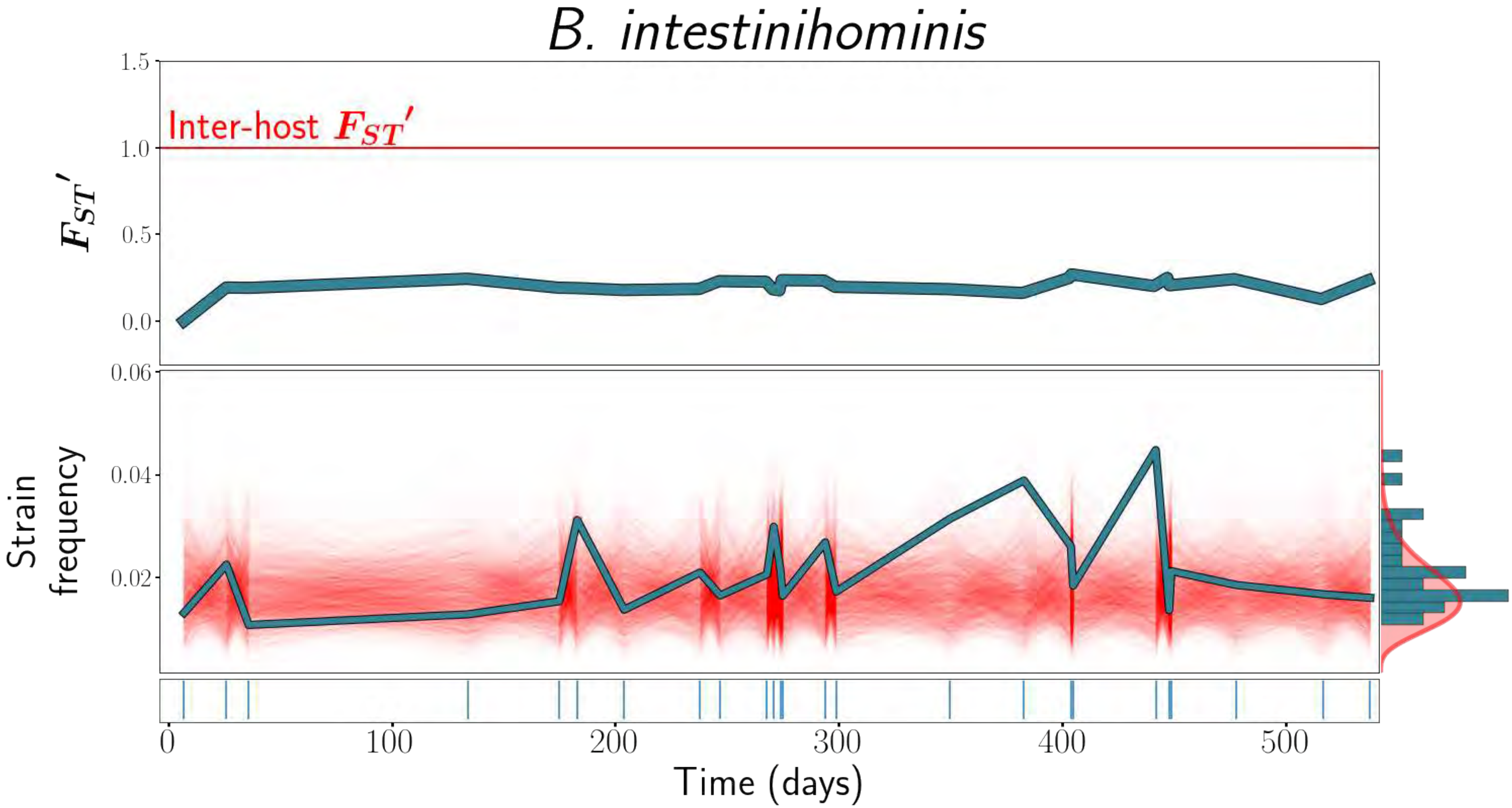

B. intestinihominis
Inter-host $F_{ST}'$
$F_{ST}'$
1.5
1.0
0.5
0.0
Strain frequency
0.06
0.04
0.02
0
100
200
300
400
500
Time (days)

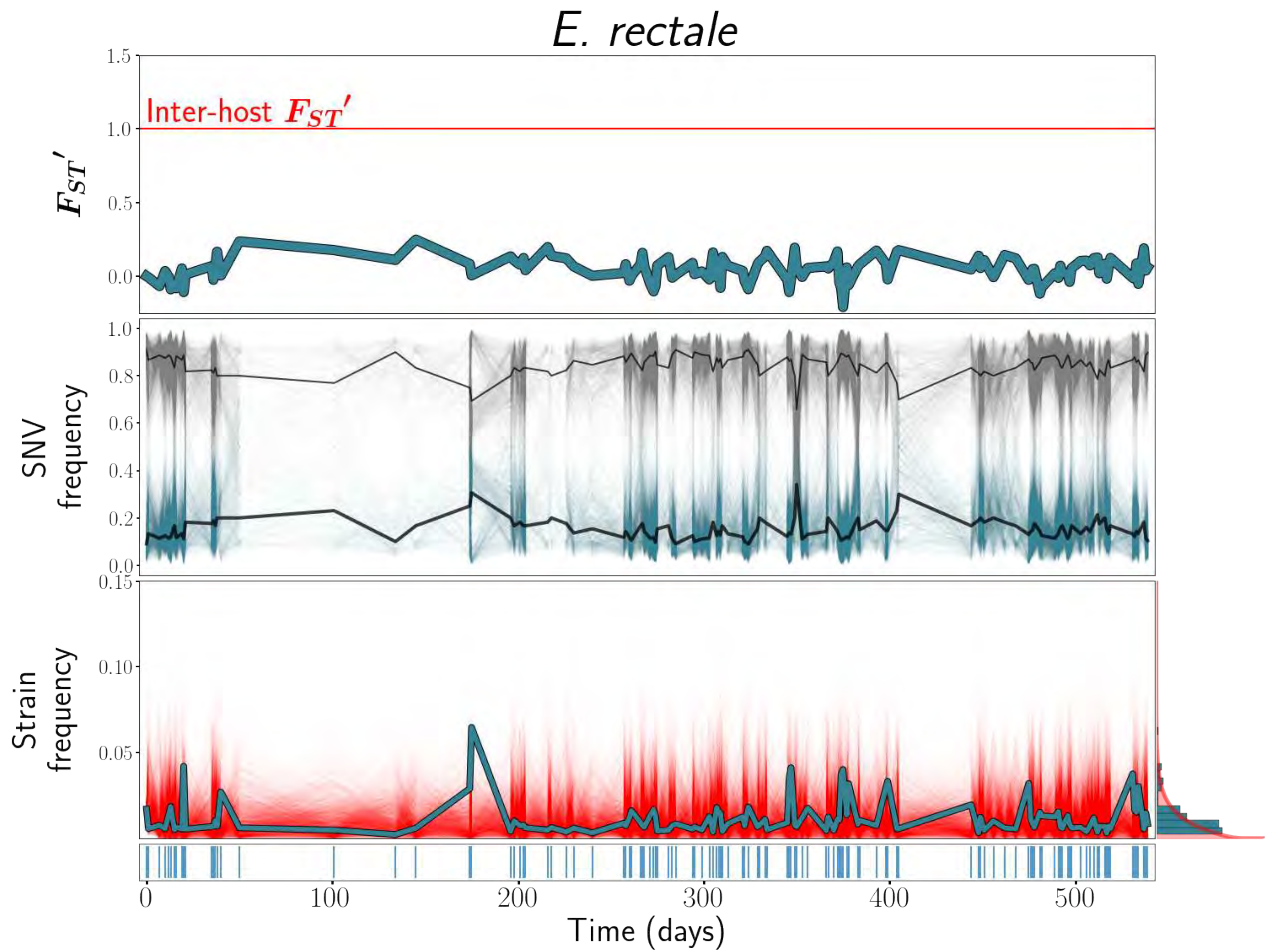

E. rectale
Inter-host $F_{ST}'$
$F_{ST}'$
1.5
1.0
0.5
0.0
SNV frequency
1.0
0.8
0.6
0.4
0.2
0.0
Strain frequency
0.15
0.10
0.05
0
100
200
300
400
500
Time (days)

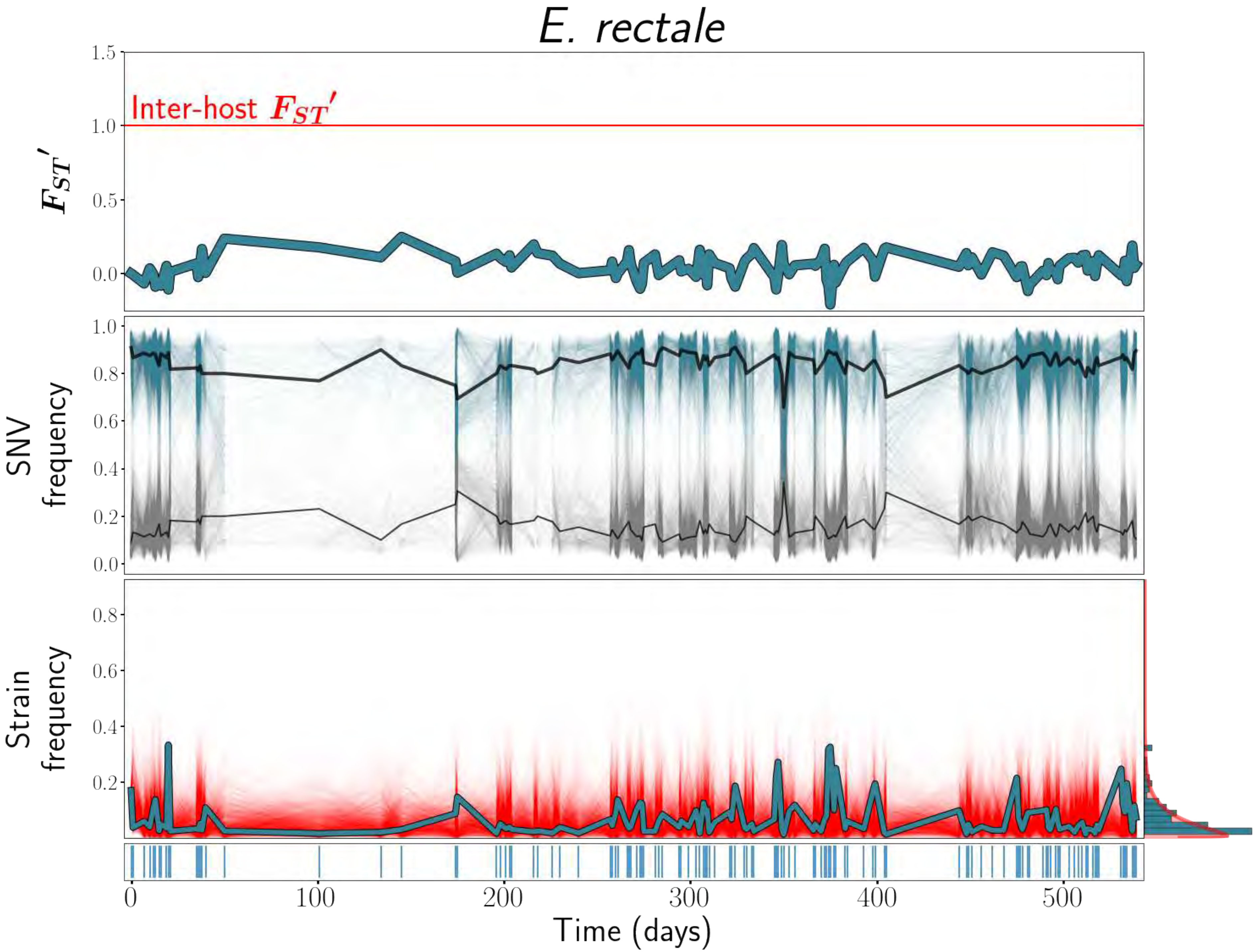

E. rectale
Inter-host $F_{ST}'$
$F_{ST}'$
SNV frequency
Strain frequency
Time (days)

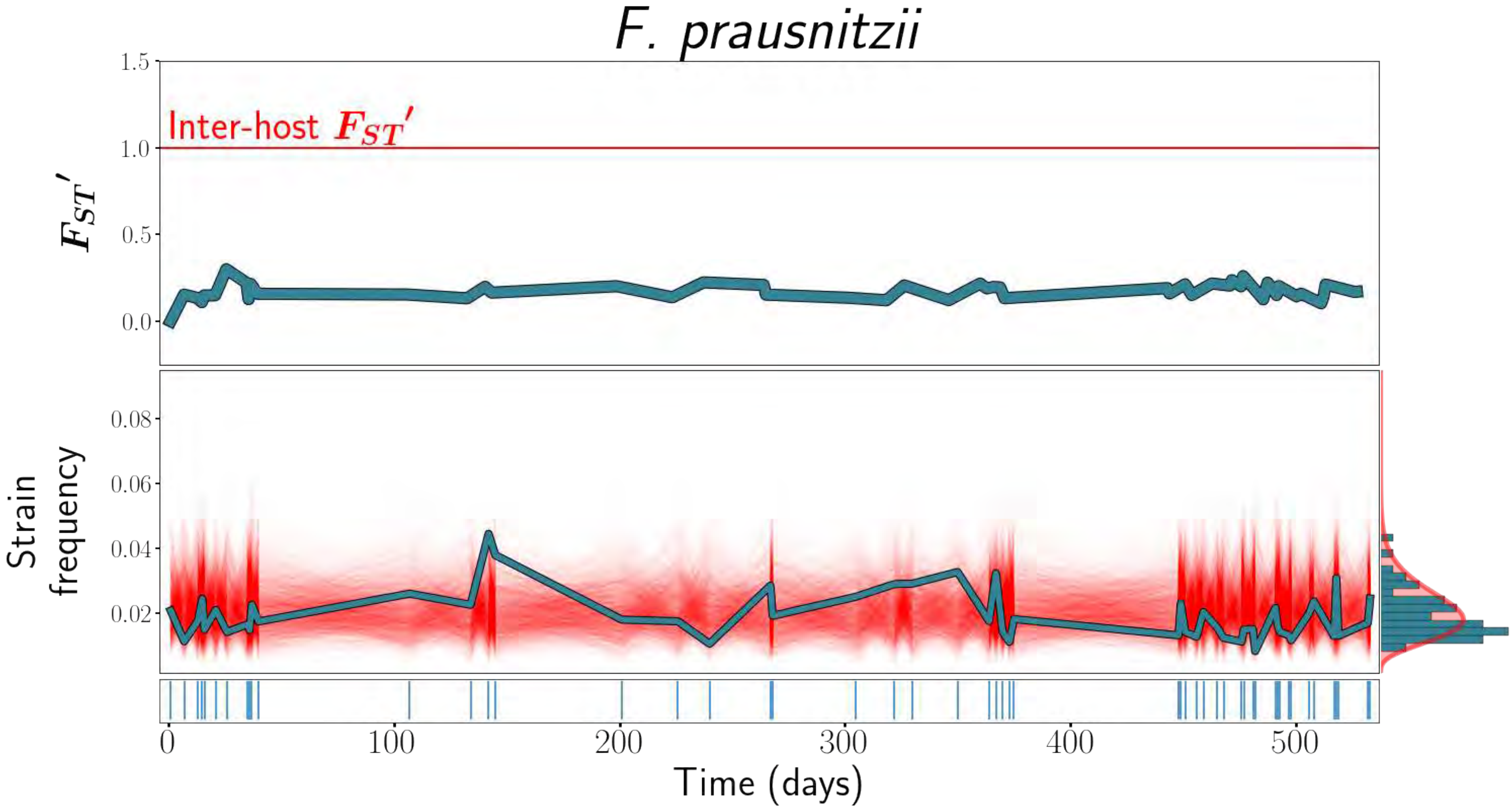

F. prausnitzii
Inter-host $F_{ST}'$
$F_{ST}'$
1.5
1.0
0.5
0.0
Strain frequency
0.08
0.06
0.04
0.02
0
100
200
300
400
500
Time (days)

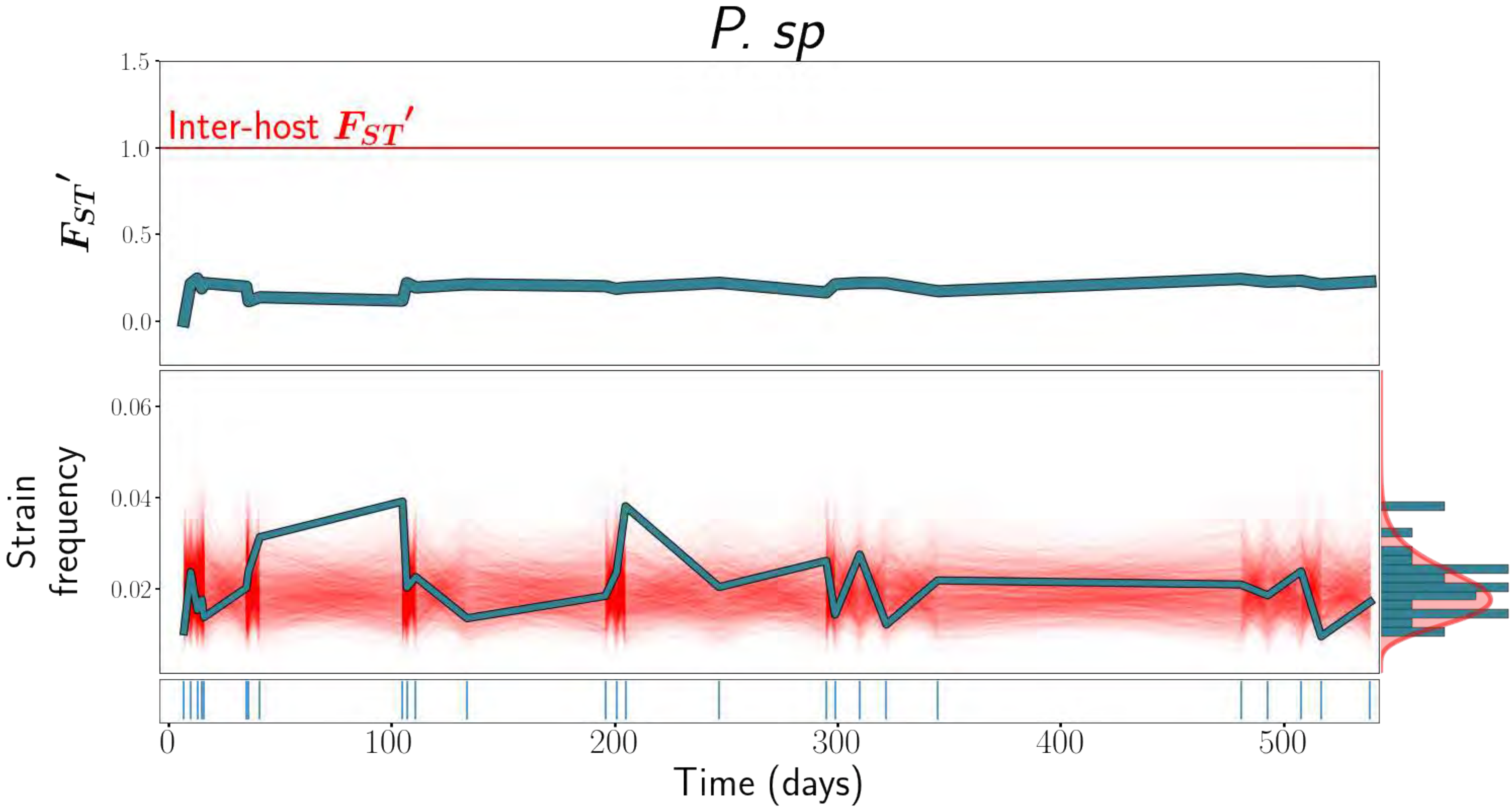

P. sp
Inter-host $F_{ST}'$
$F_{ST}'$
1.5
1.0
0.5
0.0
Strain frequency
0.06
0.04
0.02
0
100
200
300
400
500
Time (days)

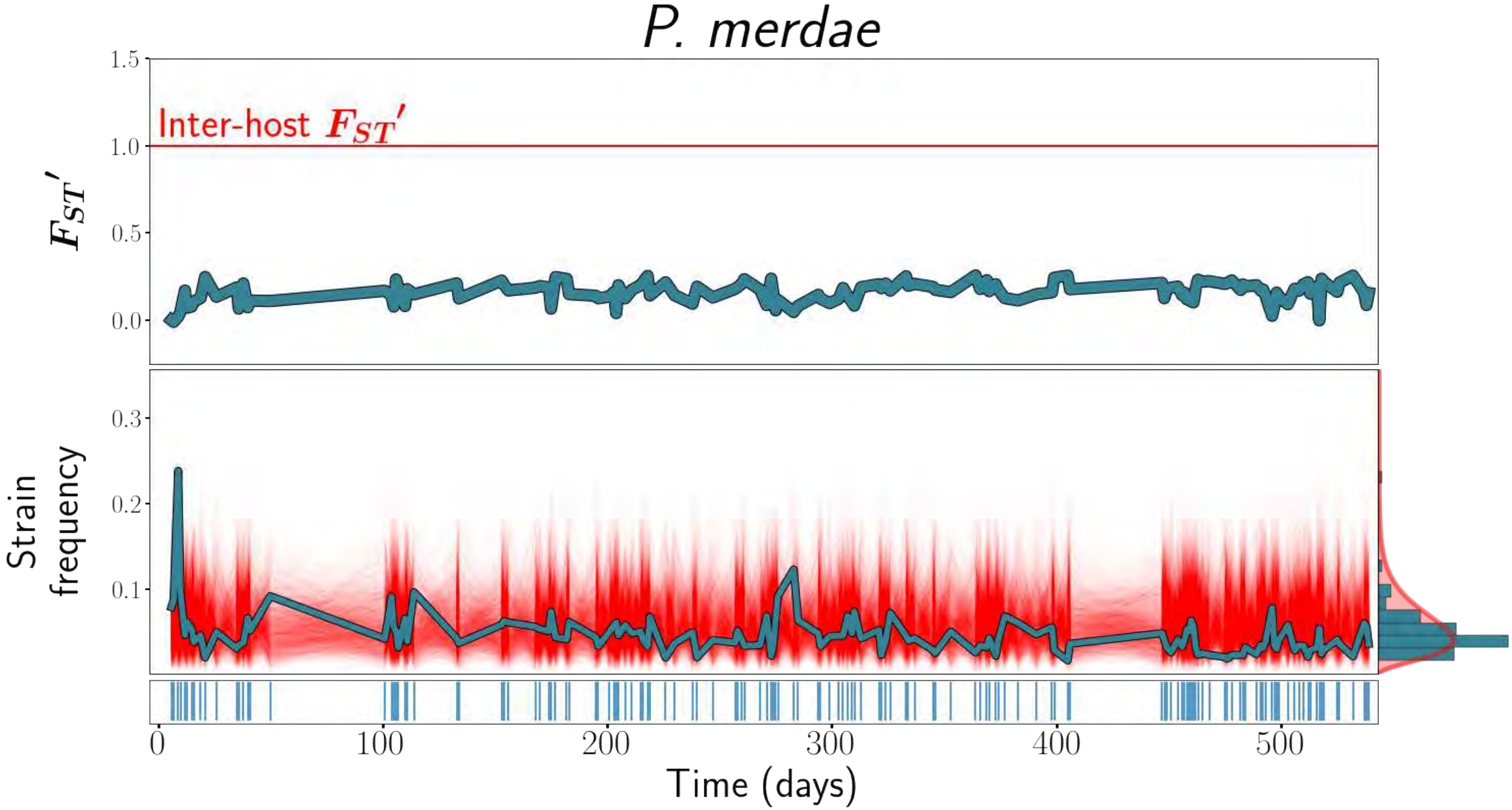

P. merdae
Inter-host $F_{ST}'$
$F_{ST}'$
1.5
1.0
0.5
0.0
Strain frequency
0.3
0.2
0.1
0
100
200
300
400
500
Time (days)

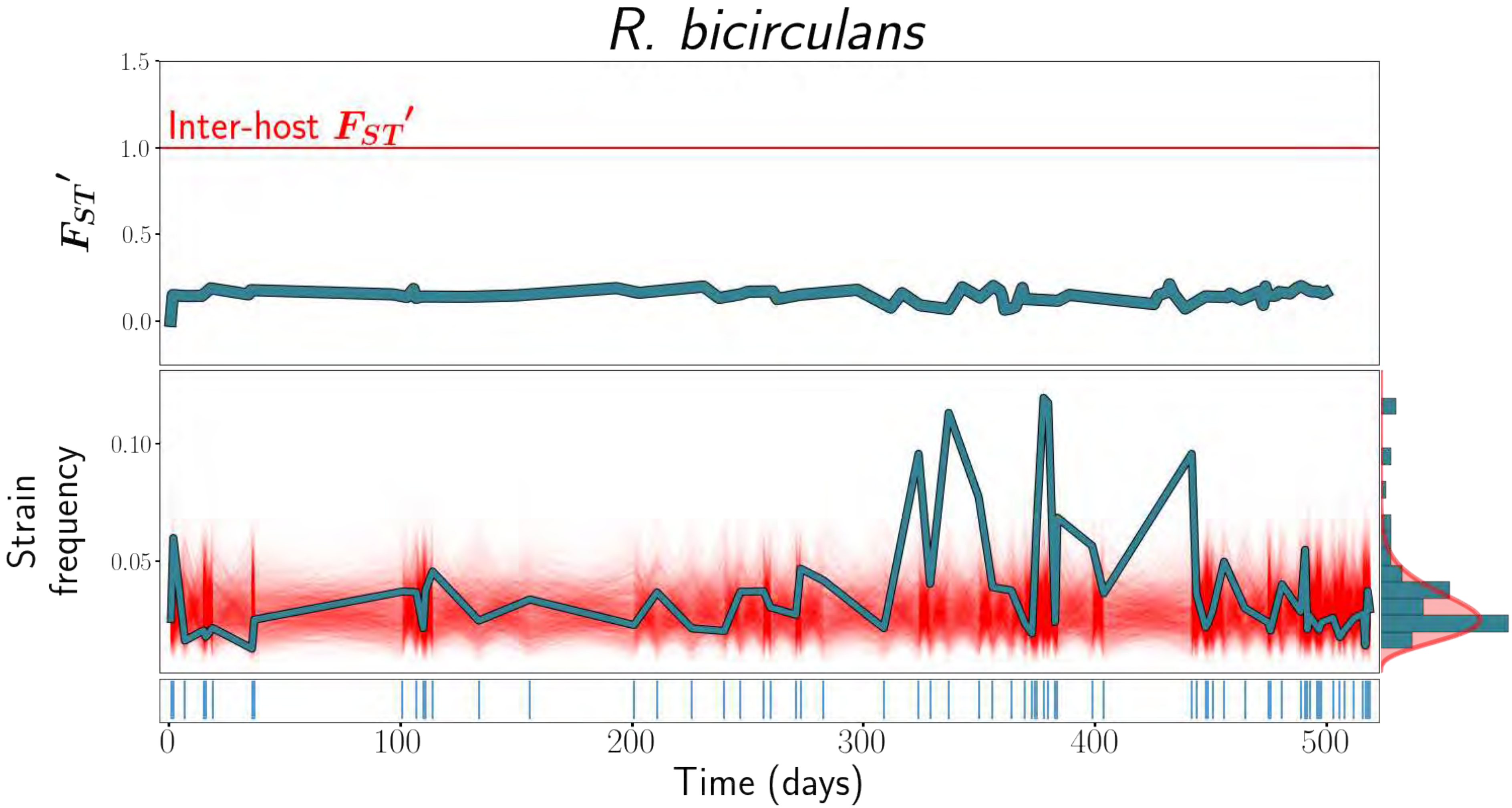

R. bicirculans
1.5
1.0
0.5
0.0
Inter-host $F_{ST}'$
$F_{ST}'$
0.10
0.05
Strain frequency
0
100
200
300
400
500
Time (days)

### *R. bromii*

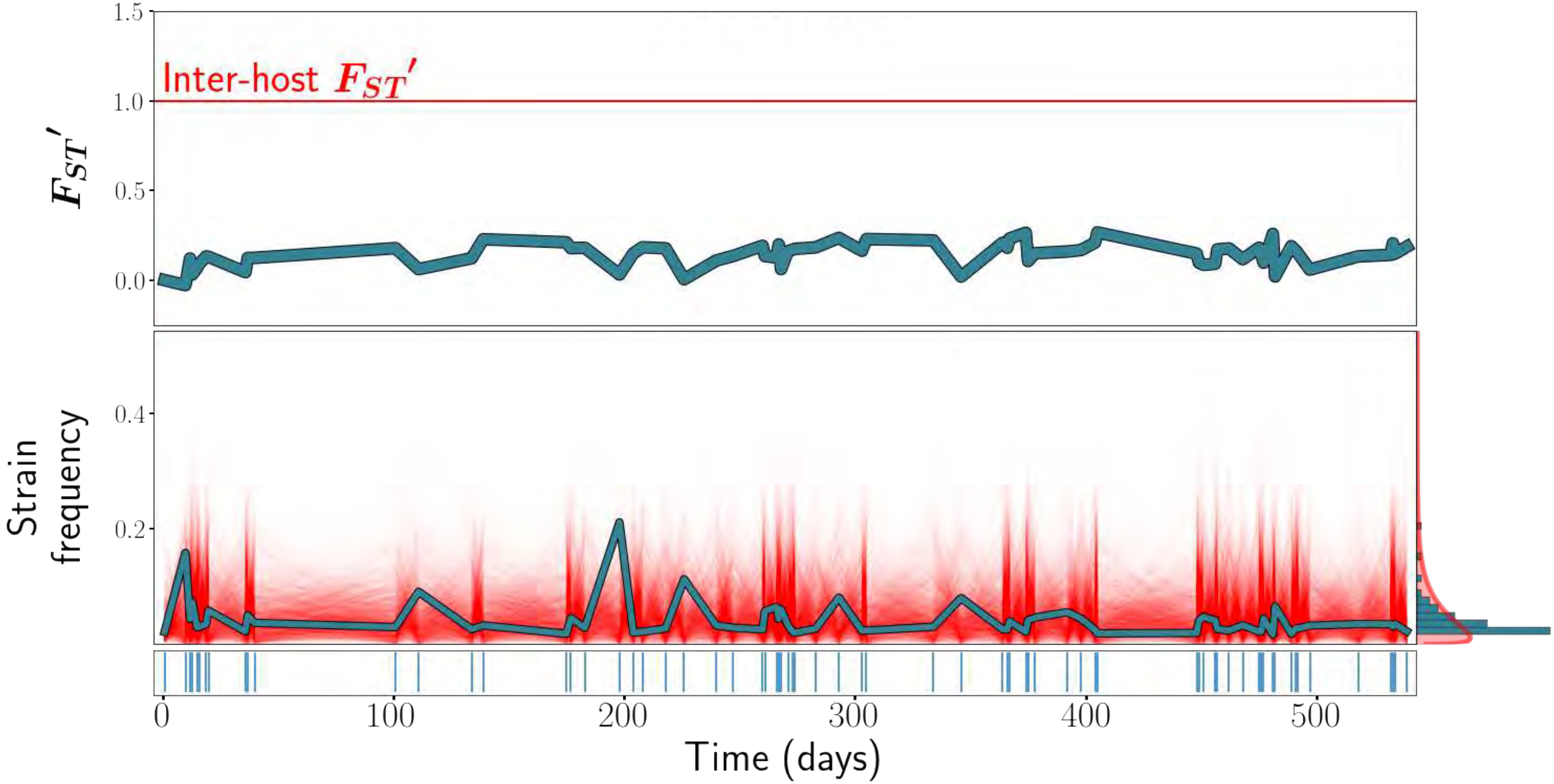

Inter-host $F_{ST}'$
$F_{ST}'$
1.5
1.0
0.5
0.0
Strain frequency
0.4
0.2
0
100
200
300
400
500
Time (days)
