## Supplementary material for "Ecological Stability Emerges at the Level of Strains in the Human Gut Microbiome": S3 Text

### S4 Text

*$F_{ST}$'*, strain frequency, and strain abundance dynamics plots for all species analyzed, for host *an.* These plots are analogous to Main Text Figure 1. When only a single strain was detected, only *$F_{ST}$'* and strain abundance dynamics plots, but no strain frequency plot, is included.

*Table of contents*


| | |
|---|---|
| *Alistipes putredinis A* | *1* |
| *Bacteroides clarus A* | *2* |
| *Bacteroides massilensis A* | *3* |
| *Bacteroides uniformis A* | *4* |
| *Bacteroides uniformis B* | *5* |
| *Bacteroides vulgatus A* | *6* |
| *Bacteroides vulgatus B* | *7* |
| *Bacteroides vulgatus C* | *8* |
| *Bacteroides xylanisolvens A* | *9* |
| *Barnesiella intestinihominis A* | *10* |
| *Eubacterium rectale A* | *11* |
| *Eubacterium rectale B* | *12* |
| *Paraprevotella clara A* | *13* |
| *Ruminoccocus bromii A* | *14* |
| *Sutterella wadsworthensis A* | *15* |
| *Sutterella wadsworthensis B* | *16* |

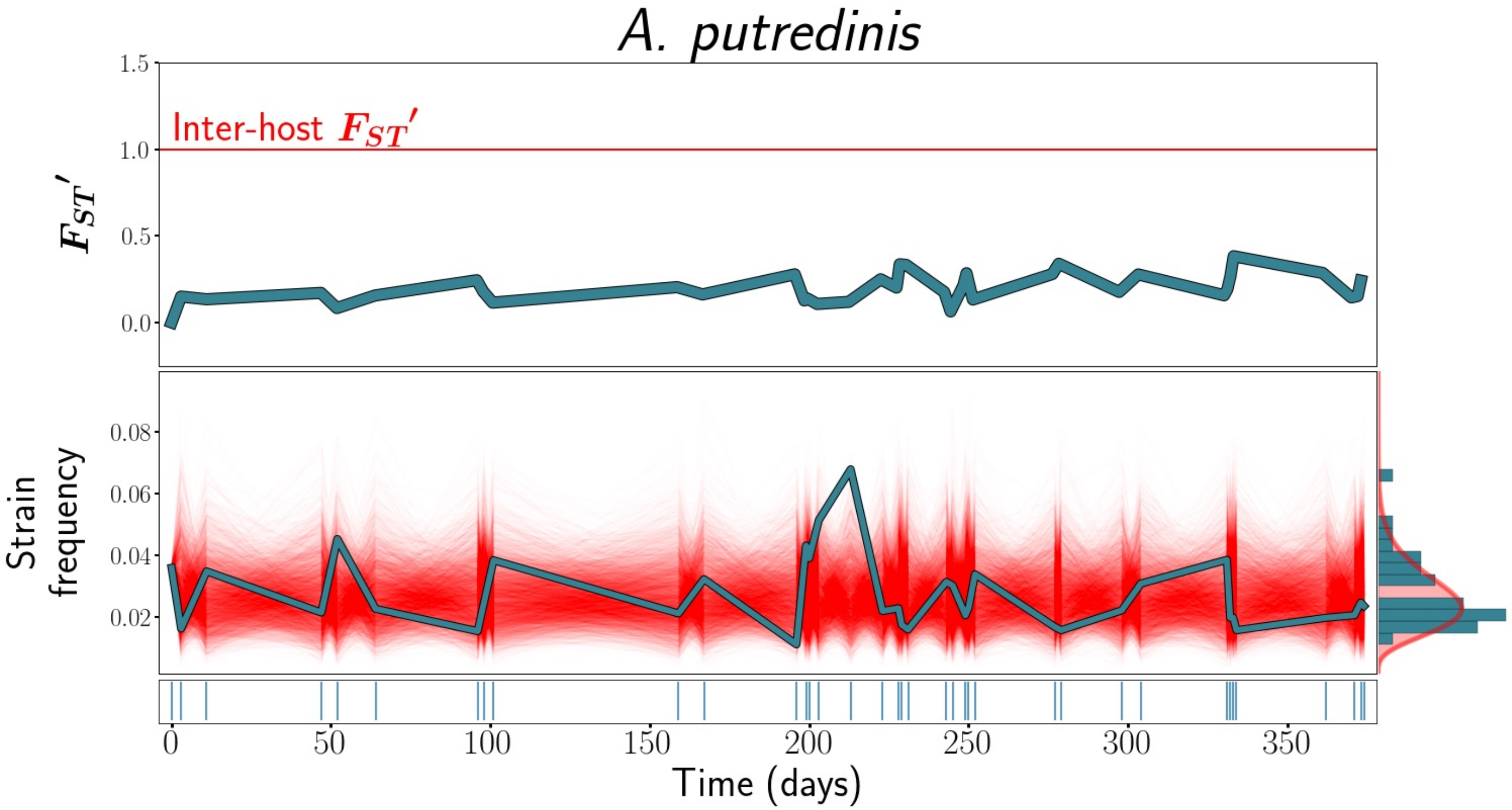

A. putredinis
1.5
Inter-host $F_{ST}'$
1.0
$F_{ST}'$
0.5
0.0
0.08
0.06
Strain frequency
0.04
0.02
0
50
100
150
200
250
300
350
Time (days)

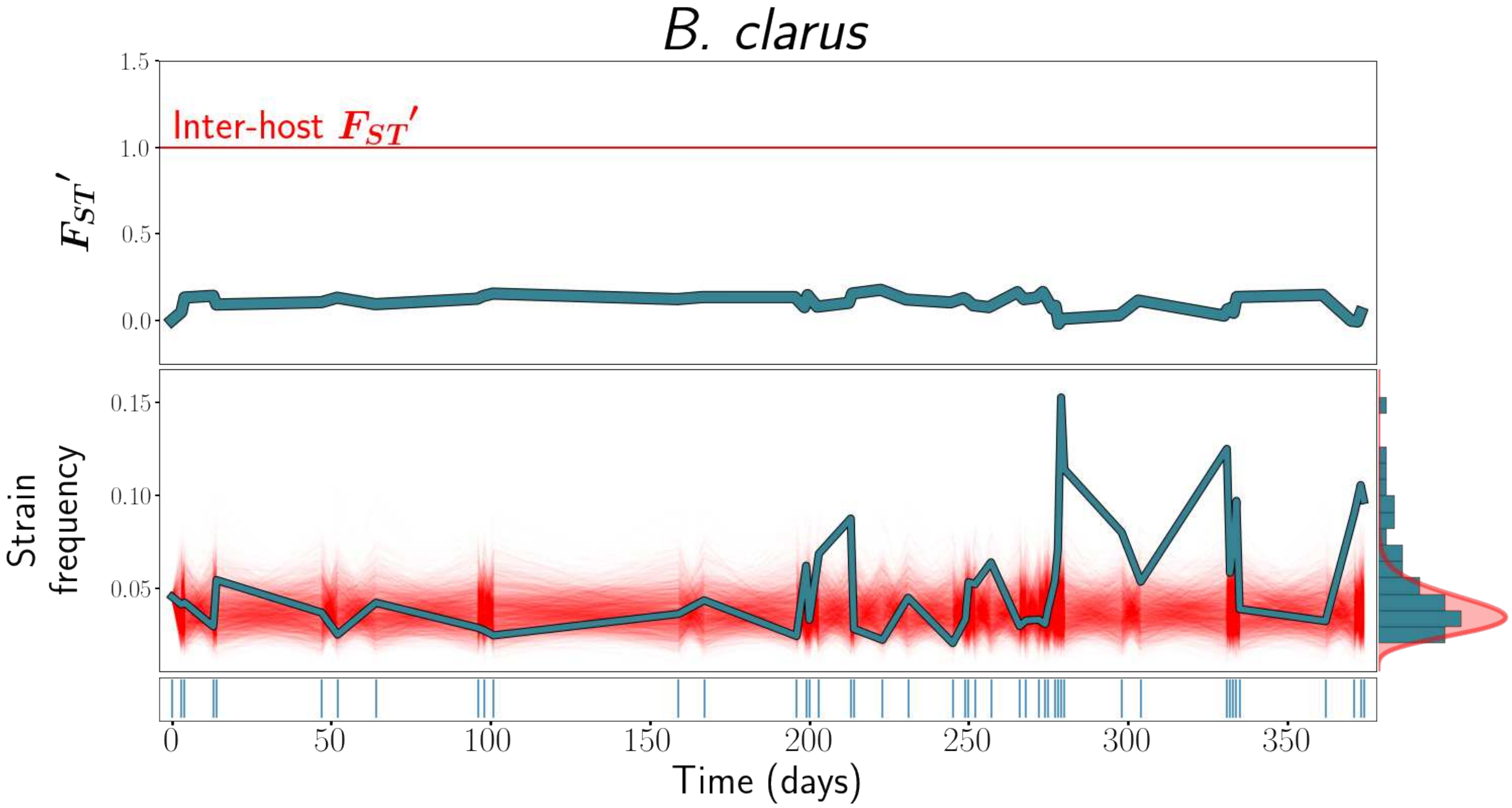

B. clarus
Inter-host $F_{ST}'$
$F_{ST}'$
1.5
1.0
0.5
0.0
Strain frequency
0.15
0.10
0.05
0
50
100
150
200
250
300
350
Time (days)

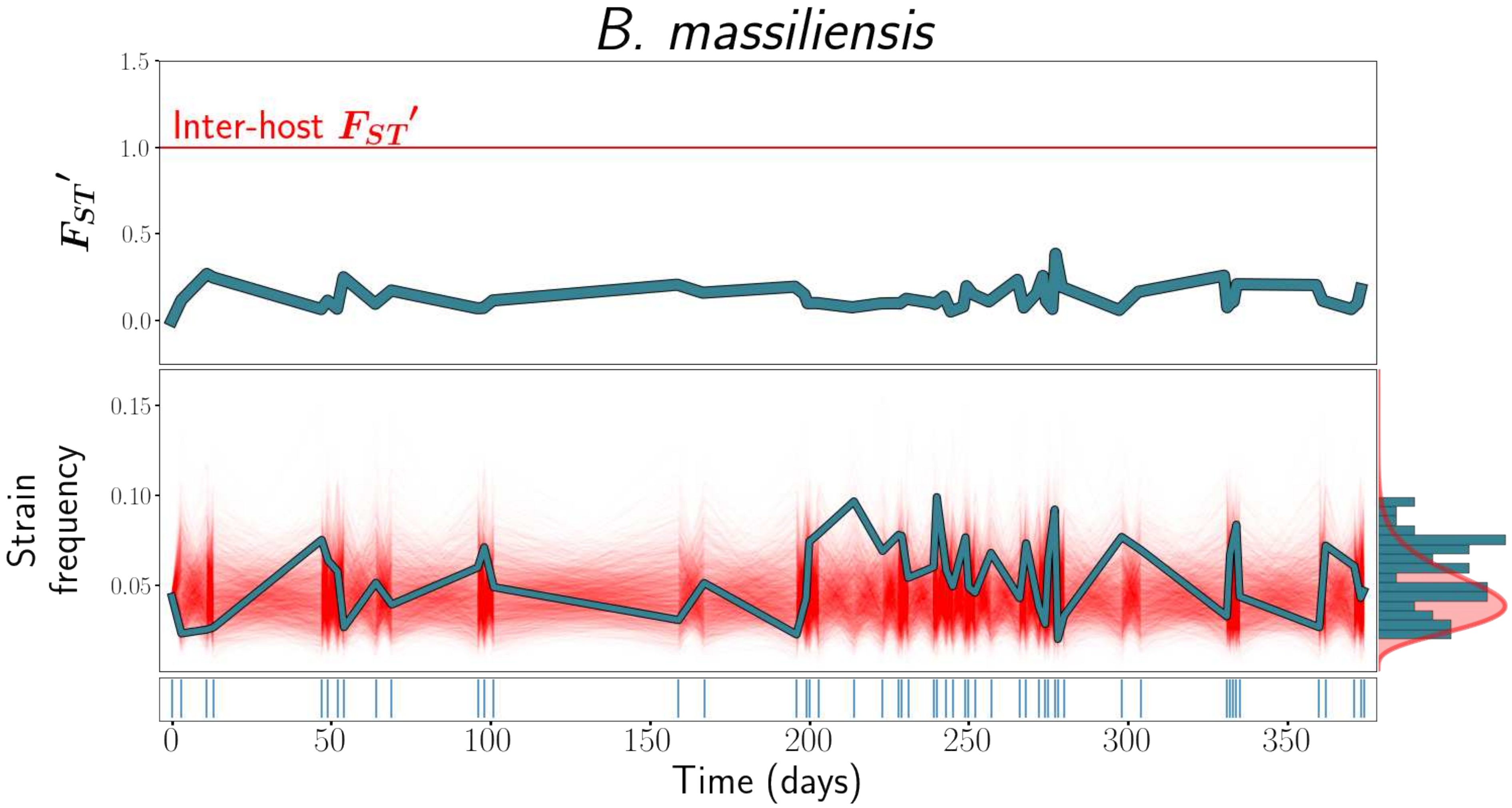
B. massiliensis
Inter-host $F_{ST}'$
$F_{ST}'$
1.5
1.0
0.5
0.0
Strain frequency
0.15
0.10
0.05
0
50
100
150
200
250
300
350
Time (days)

### *B. uniformis*

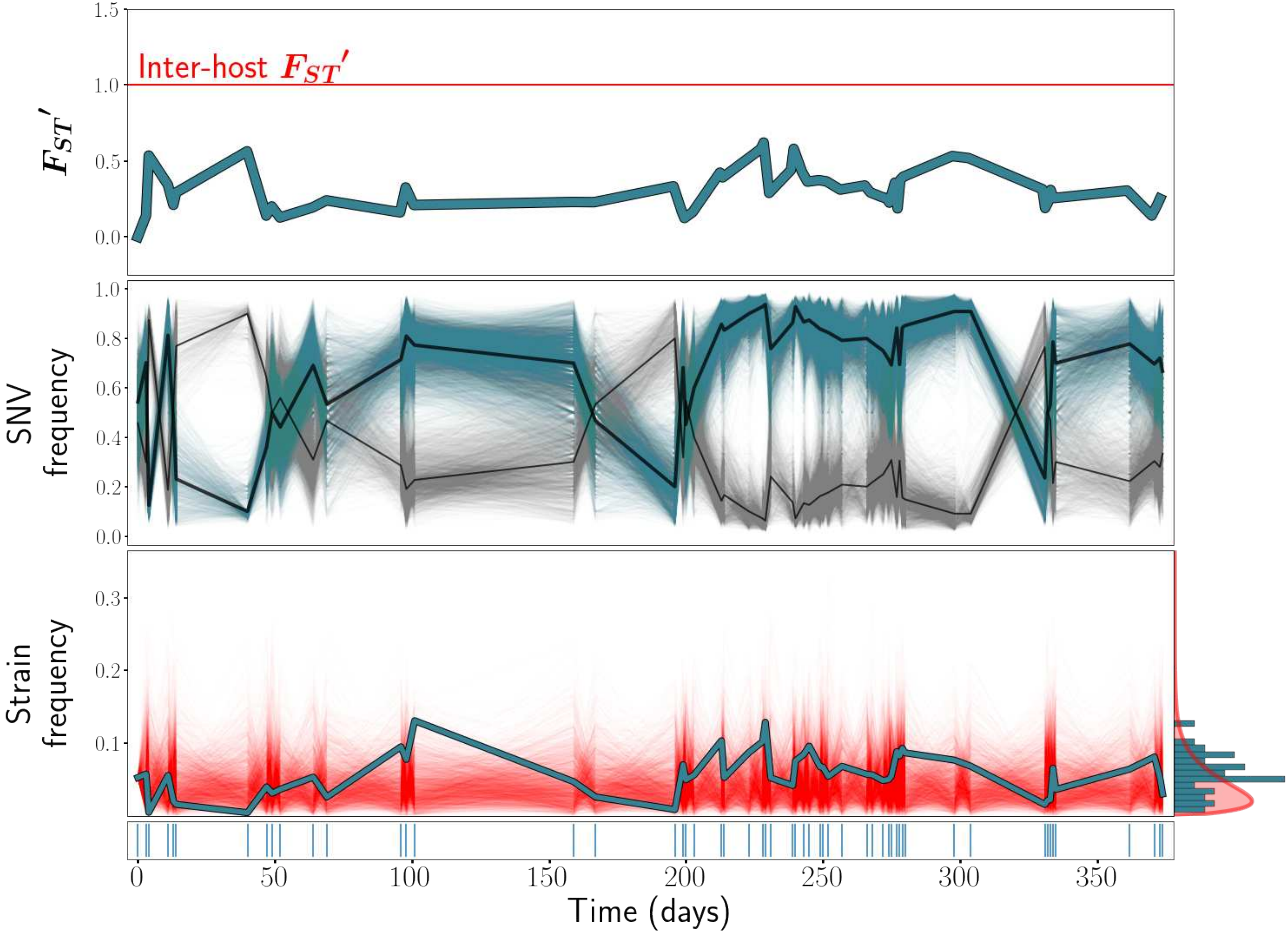

Inter-host $F_{ST}'$
$F_{ST}'$
1.5
1.0
0.5
0.0
SNV frequency
1.0
0.8
0.6
0.4
0.2
0.0
Strain frequency
0.3
0.2
0.1
0
50
100
150
200
250
300
350
Time (days)

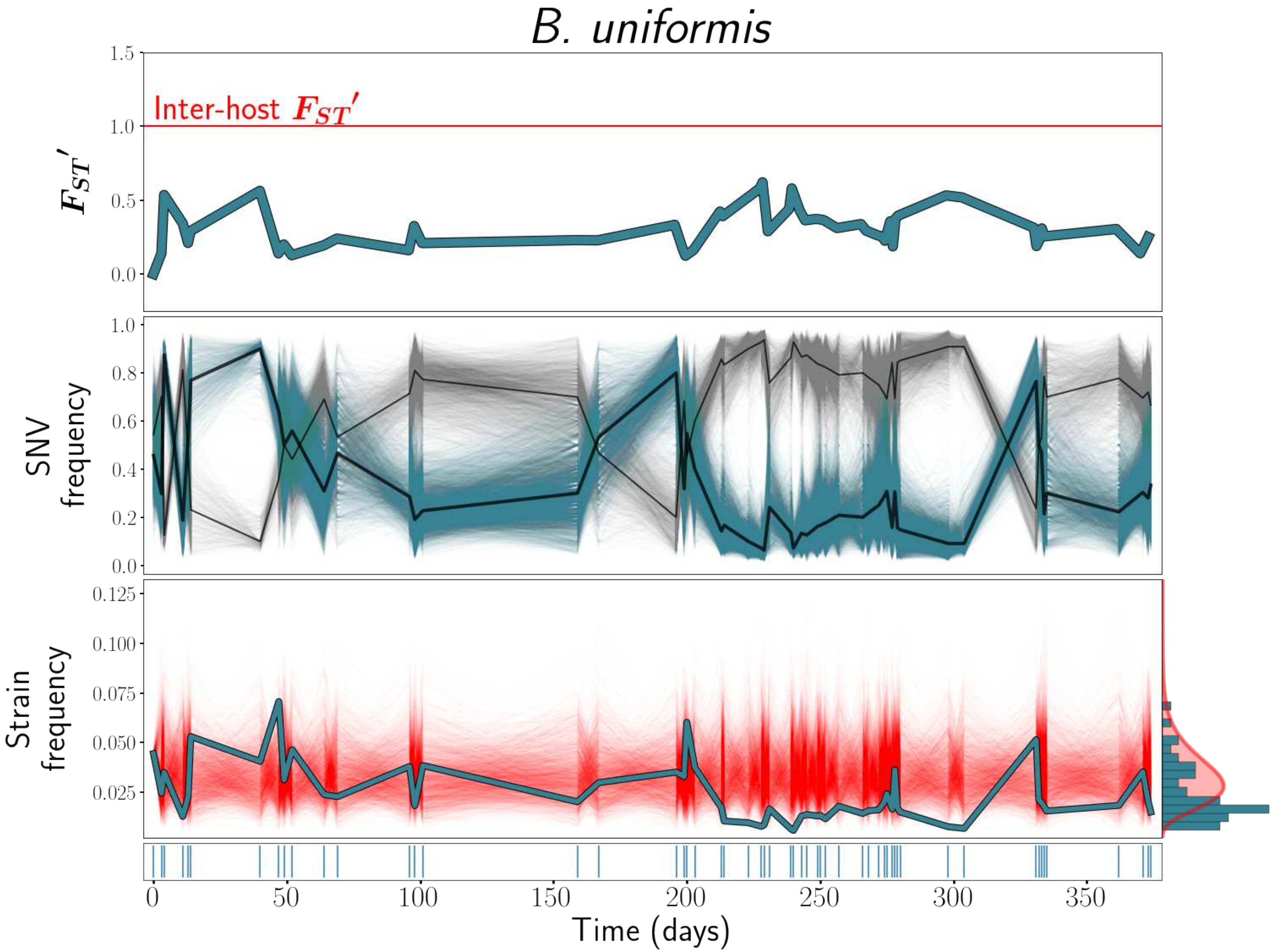

B. uniformis
Inter-host $F_{ST}'$
$F_{ST}'$
1.5
1.0
0.5
0.0
SNV frequency
1.0
0.8
0.6
0.4
0.2
0.0
Strain frequency
0.125
0.100
0.075
0.050
0.025
0
50
100
150
200
250
300
350
Time (days)

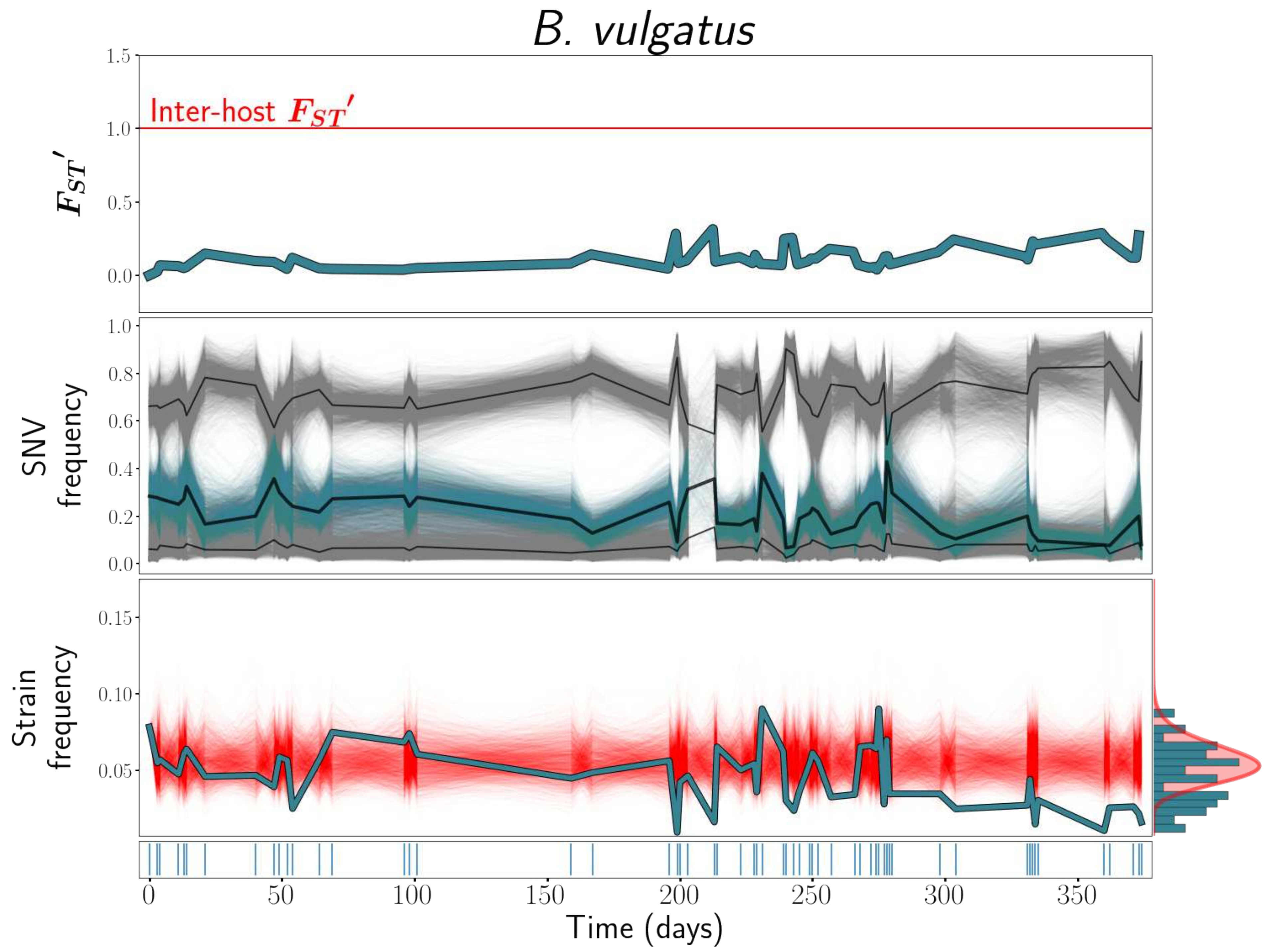
B. vulgatus
Inter-host $F_{ST}'$
$F_{ST}'$
SNV frequency
Strain frequency
Time (days)

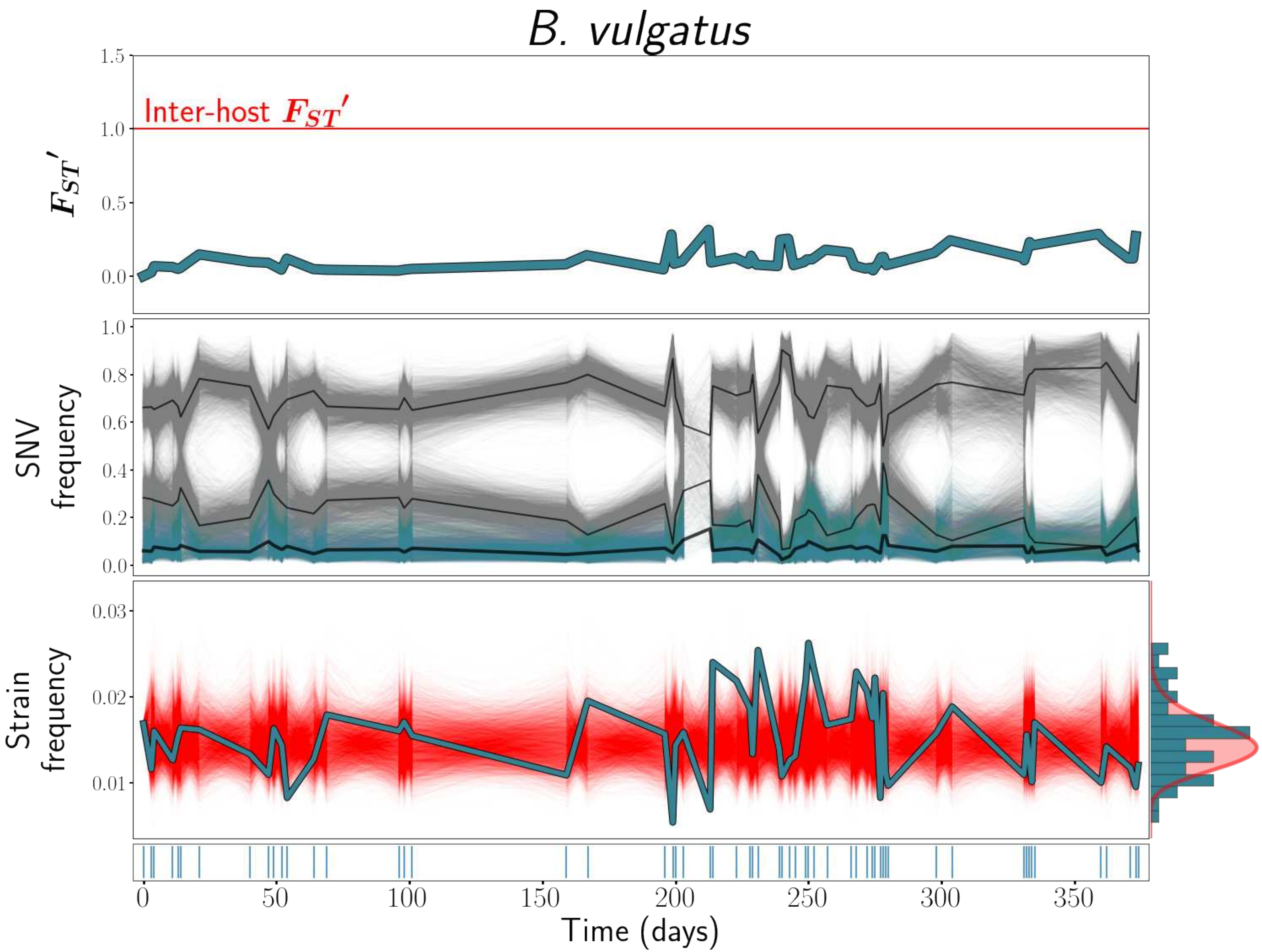

B. vulgatus
Inter-host $F_{ST}'$
$F_{ST}'$
1.5
1.0
0.5
0.0
SNV frequency
1.0
0.8
0.6
0.4
0.2
0.0
Strain frequency
0.03
0.02
0.01
0
50
100
150
200
250
300
350
Time (days)

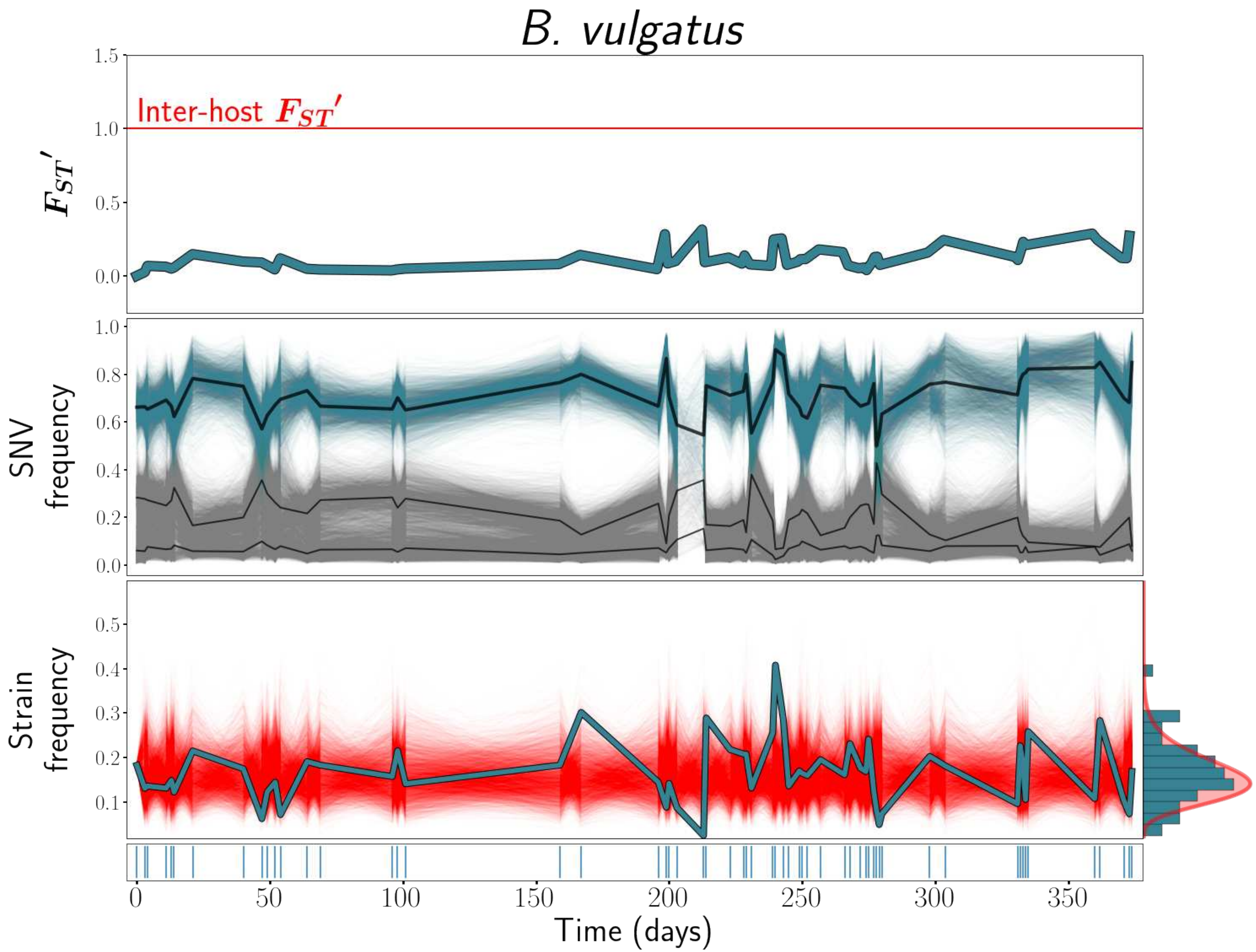

B. vulgatus
Inter-host $F_{ST}'$
$F_{ST}'$
1.5
1.0
0.5
0.0
SNV frequency
1.0
0.8
0.6
0.4
0.2
0.0
Strain frequency
0.5
0.4
0.3
0.2
0.1
0
50
100
150
200
250
300
350
Time (days)

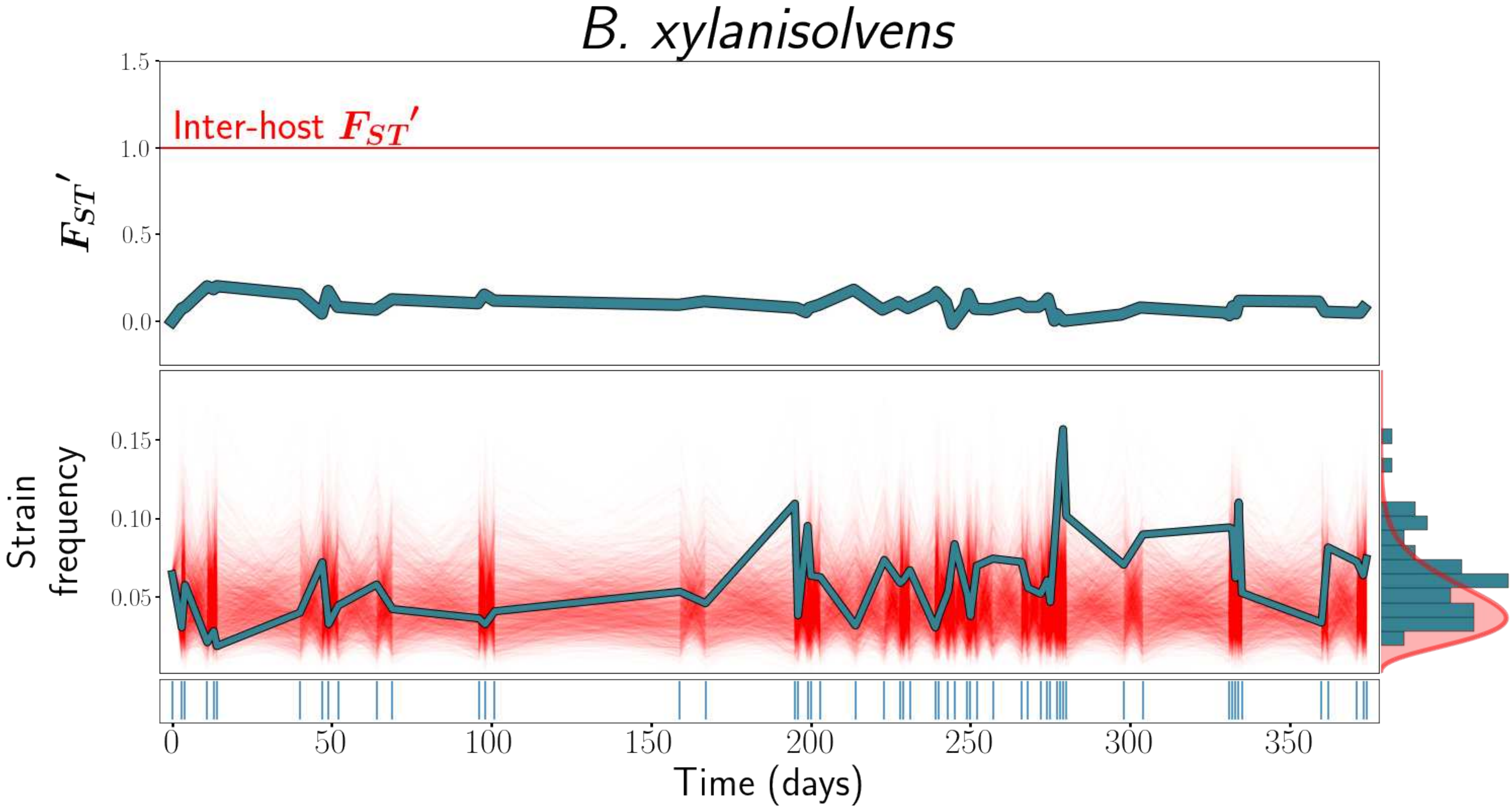
B. xylanisolvens
Inter-host $F_{ST}'$
$F_{ST}'$
1.5
1.0
0.5
0.0
Strain frequency
0.15
0.10
0.05
0
50
100
150
200
250
300
350
Time (days)

### *B. intestinihominis*

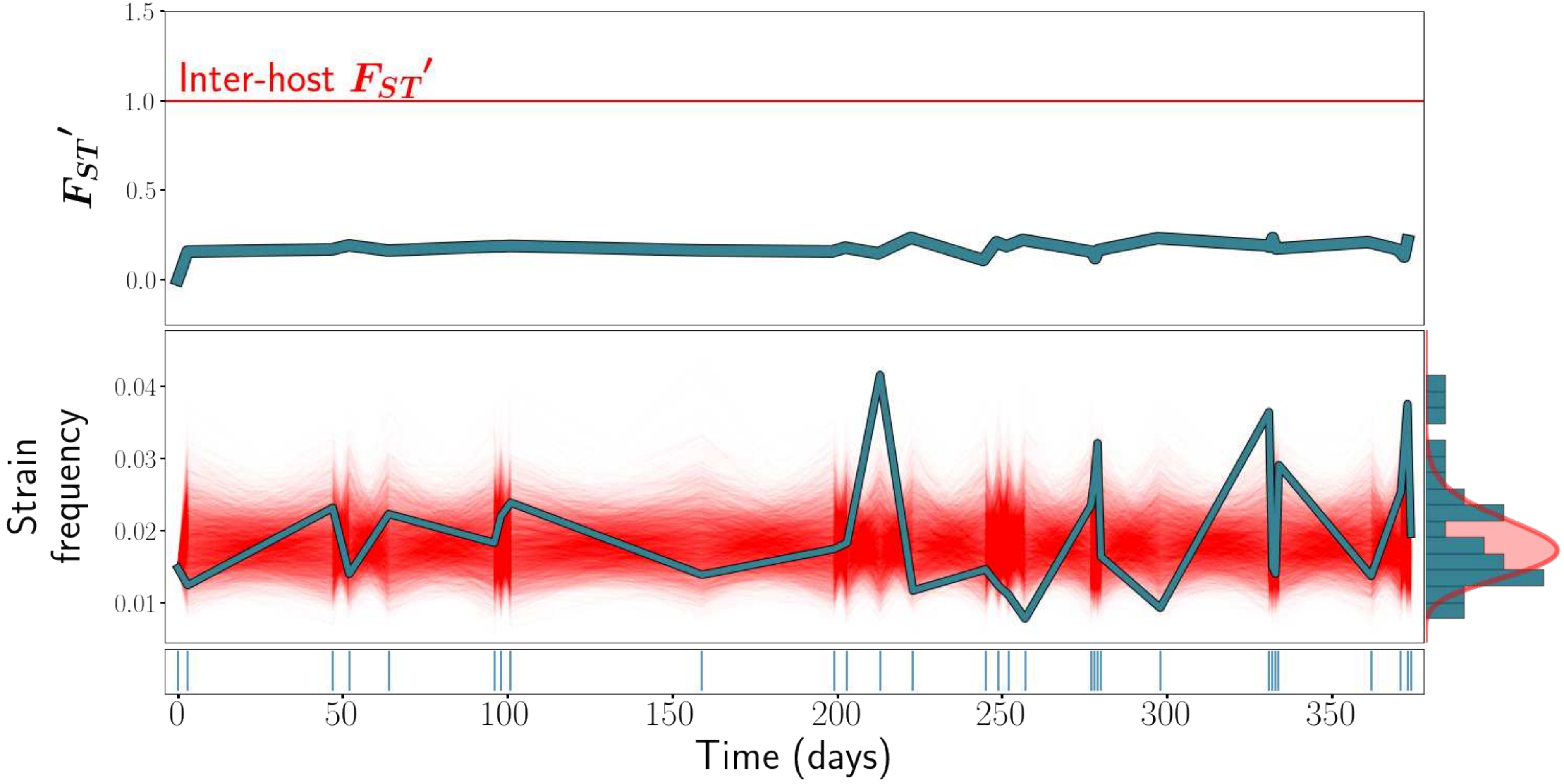

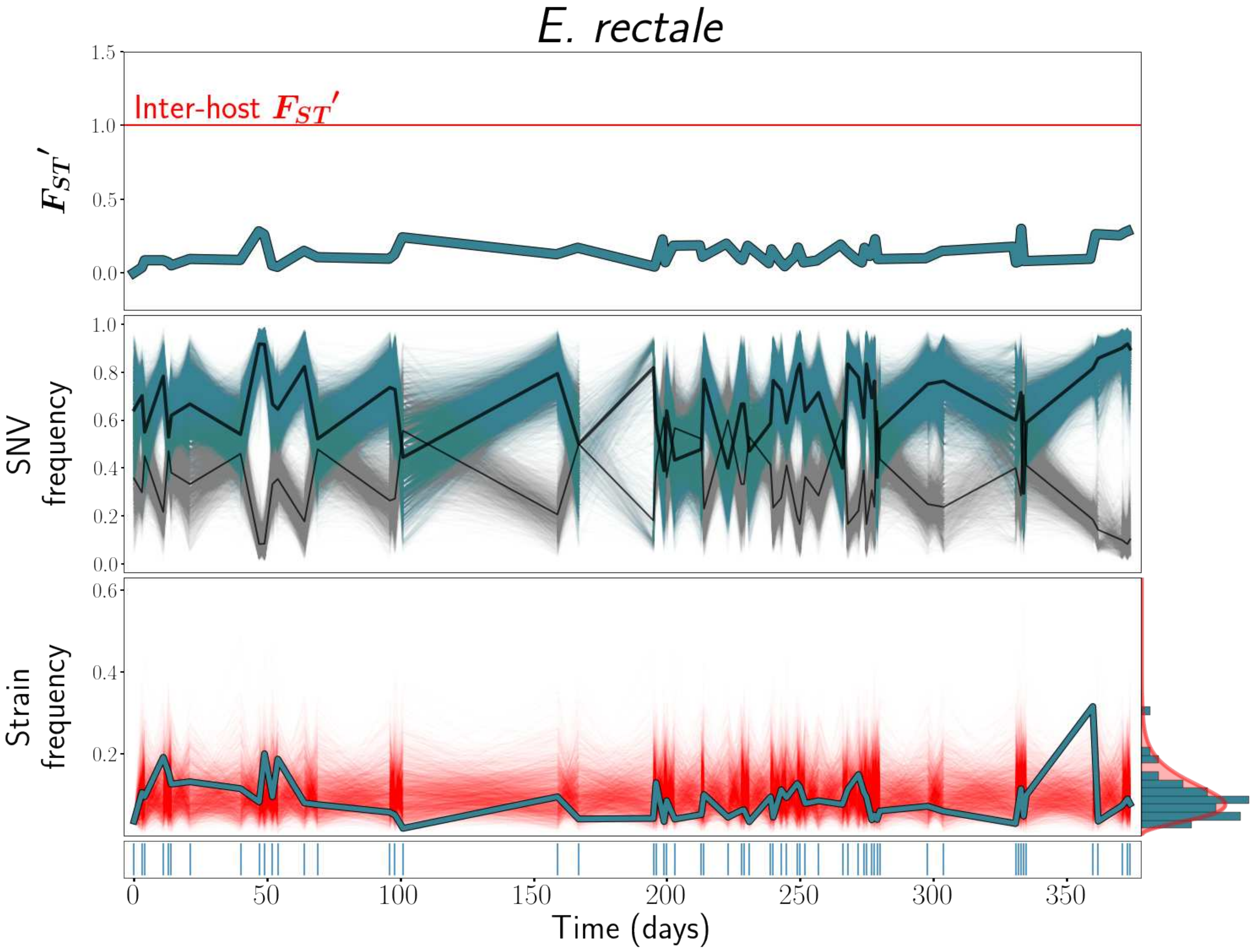
E. rectale
1.5
1.0
0.5
0.0
Inter-host $F_{ST}'$
$F_{ST}'$
1.0
0.8
0.6
0.4
0.2
0.0
SNV frequency
0.6
0.4
0.2
Strain frequency
0
50
100
150
200
250
300
350
Time (days)

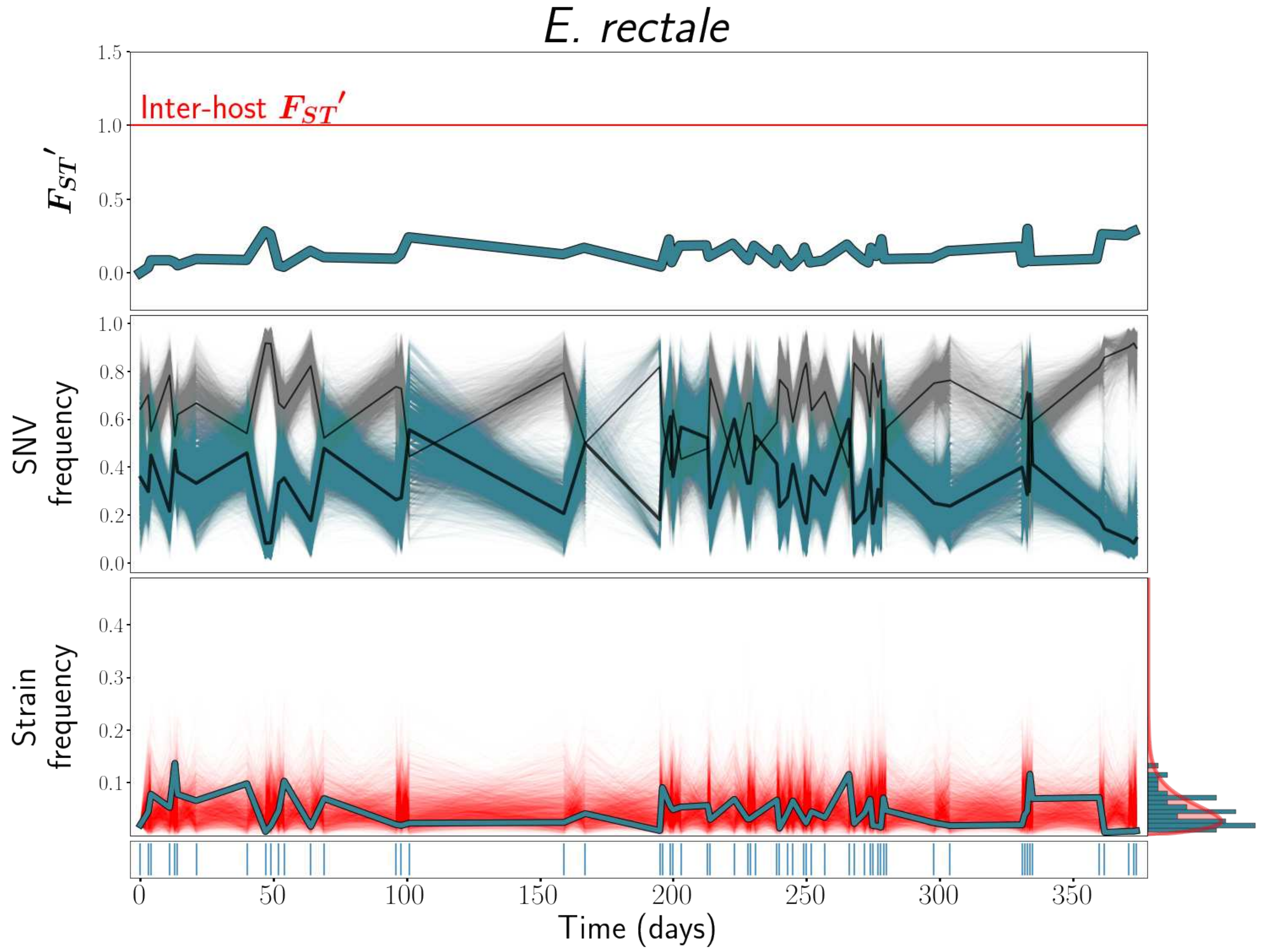
E. rectale
Inter-host $F_{ST}'$
$F_{ST}'$
SNV frequency
Strain frequency
Time (days)
1.5
1.0
0.5
0.0
0.8
0.6
0.4
0.2
0.3
0.1
0
50
100
150
200
250
300
350

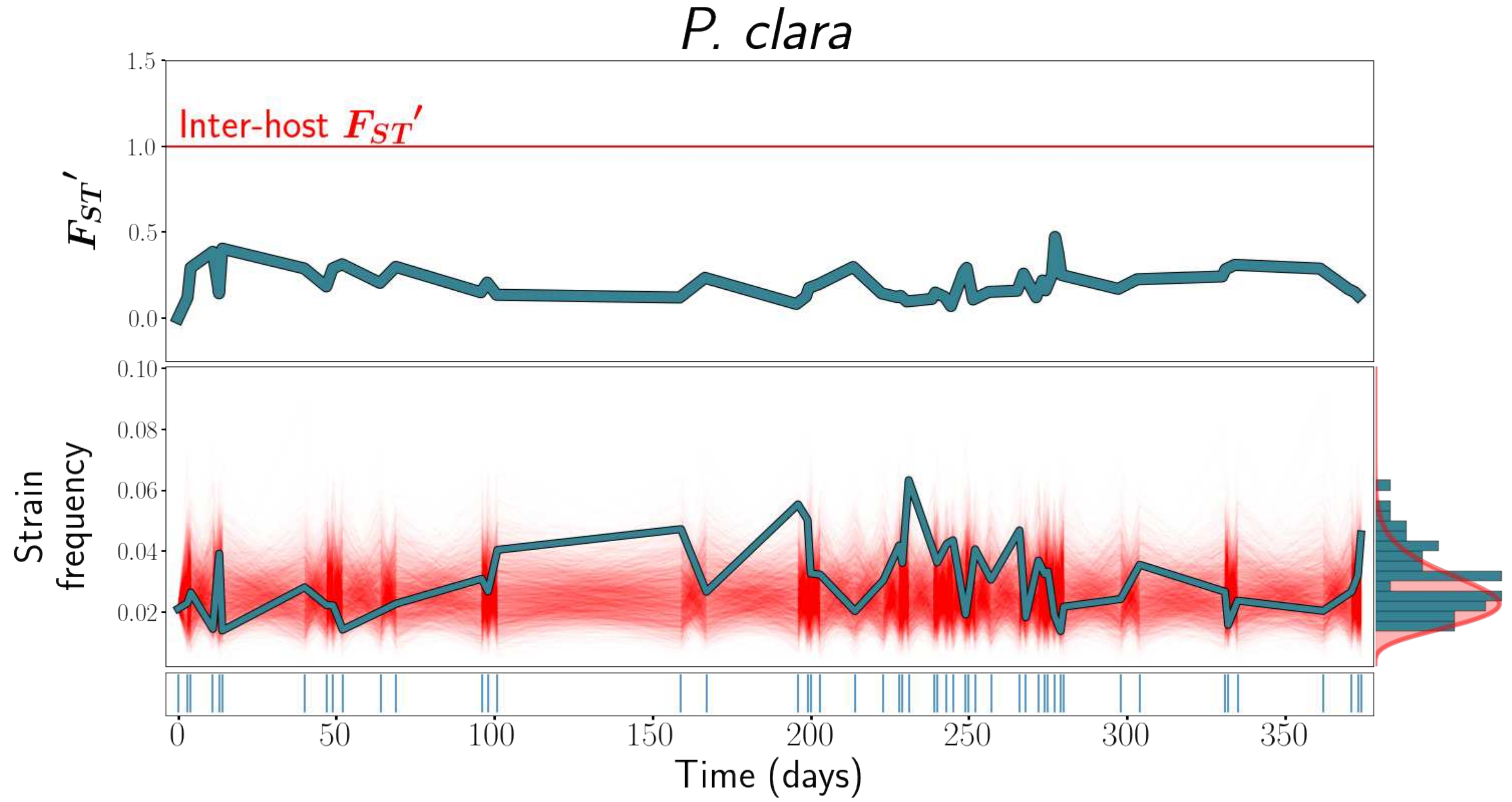

P. clara
1.5
Inter-host $F_{ST}'$
1.0
$F_{ST}'$
0.5
0.0
0.10
0.08
0.06
Strain frequency
0.04
0.02
0
50
100
150
200
250
300
350
Time (days)

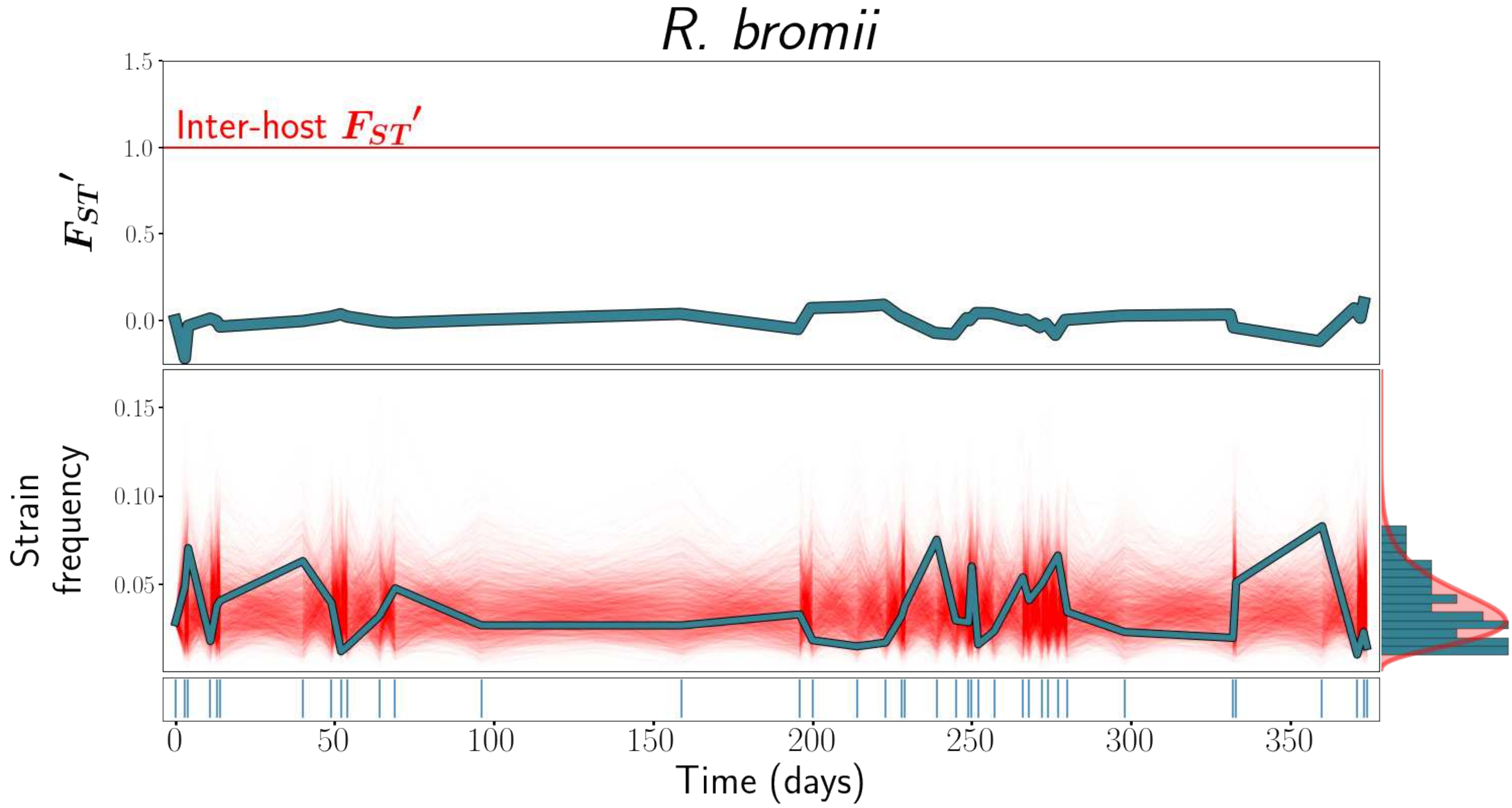
R. bromii
Inter-host $F_{ST}'$
$F_{ST}'$
Strain frequency
Time (days)
1.5
1.0
0.5
0.0
0.15
0.10
0.05
0
50
100
150
200
250
300
350

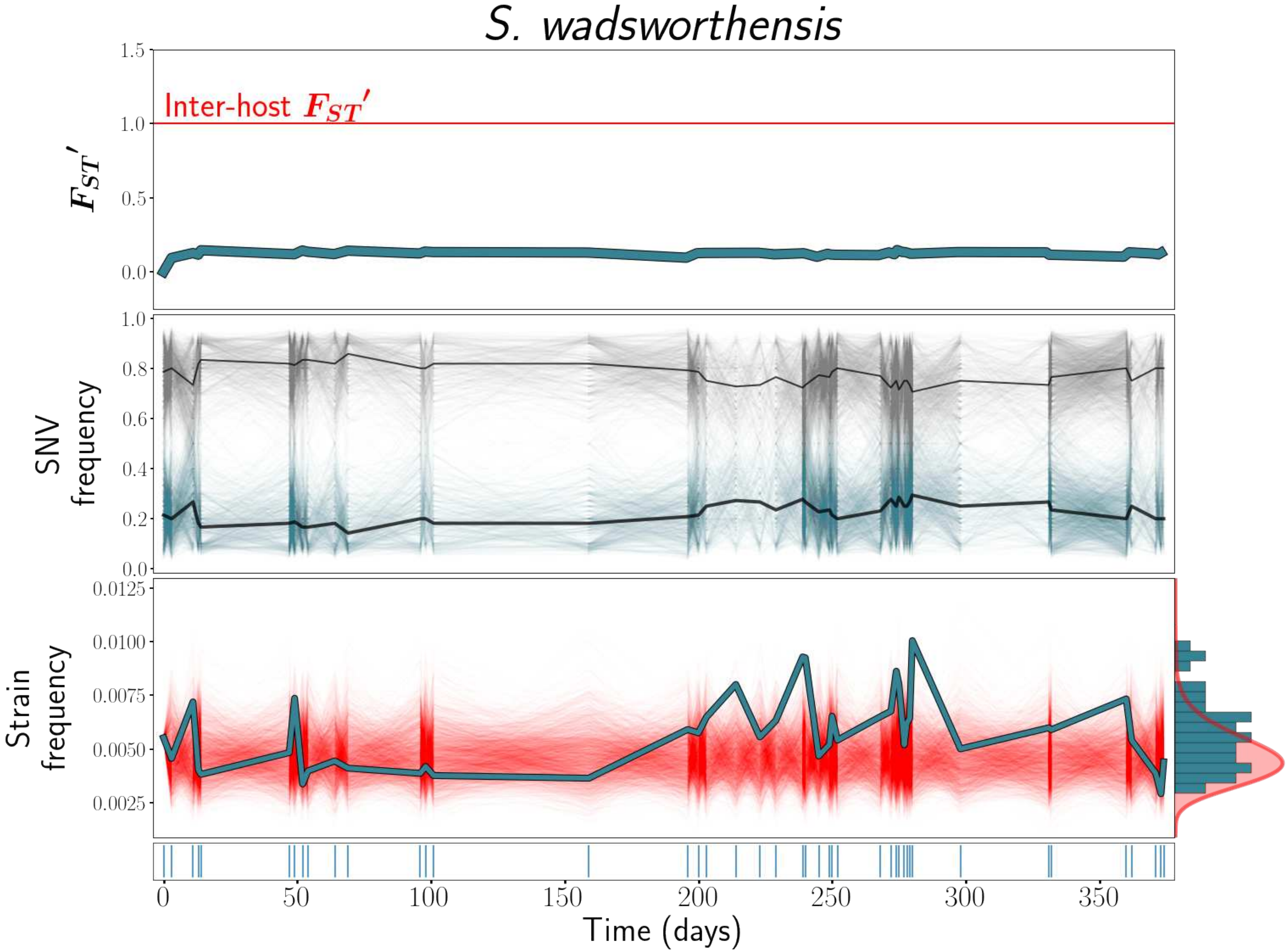
S. wadsworthensis
Inter-host $F_{ST}'$
$F_{ST}'$
SNV frequency
Strain frequency
Time (days)
1.5
1.0
0.5
0.0
0.8
0.6
0.4
0.2
0.0125
0.0100
0.0075
0.0050
0.0025
0
50
100
150
200
250
300
350

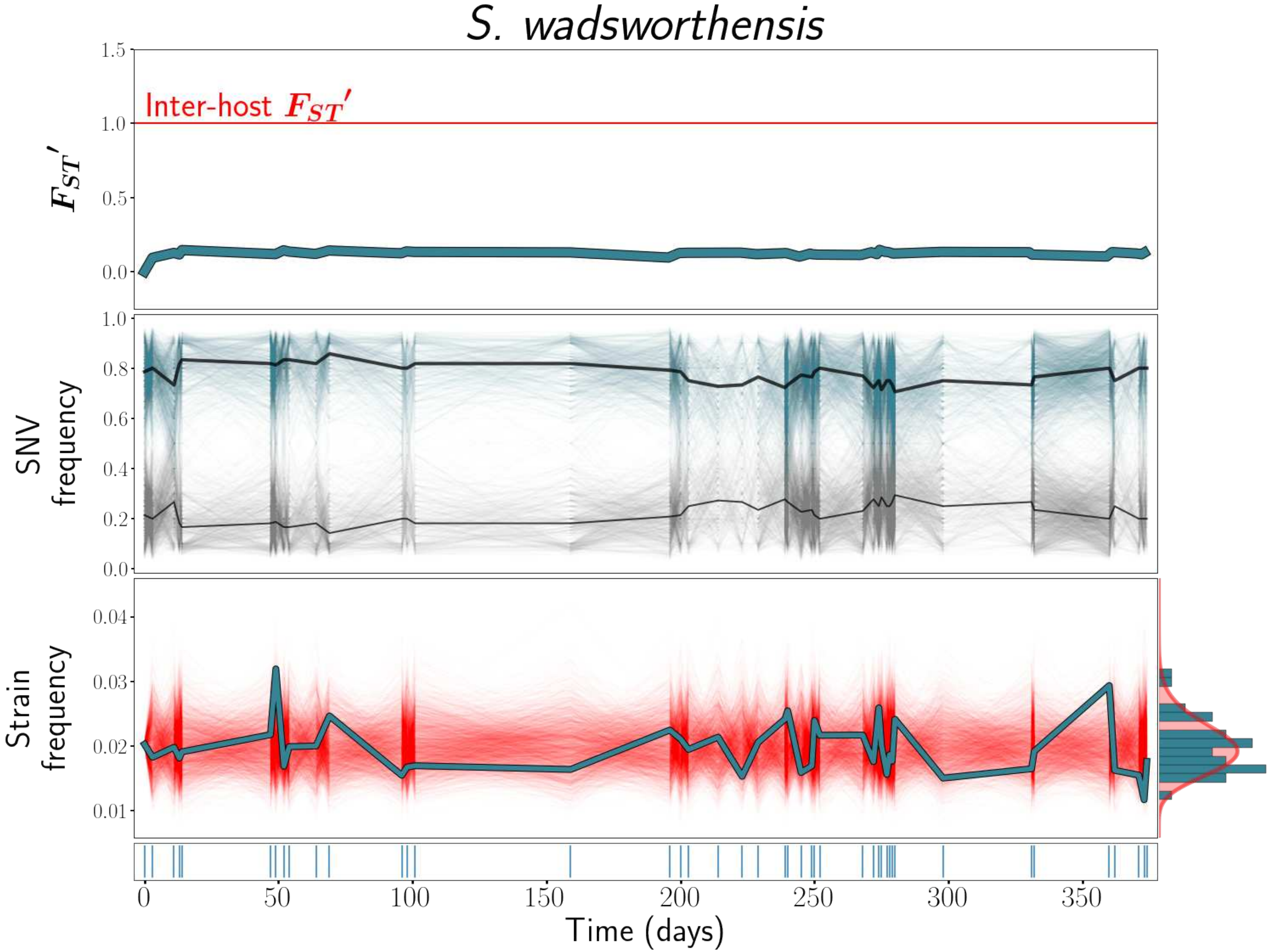
S. wadsworthensis
Inter-host $F_{ST}'$
$F_{ST}'$
SNV frequency
Strain frequency
Time (days)
1.5
1.0
0.5
0.0
0.8
0.6
0.4
0.2
0.04
0.03
0.02
0.01
0
50
100
150
200
250
300
350
