## Supplementary material for "Ecological Stability Emerges at the Level of Strains in the Human Gut Microbiome": S4 Text

### S5 Text

*$F_{ST}$'*, strain frequency, and strain abundance dynamics plots for all species analyzed, for host *ae.* These plots are analogous to Main Text Figure 1. When only a single strain was detected, only *$F_{ST}$'* and strain abundance dynamics plots, but no strain frequency plot, is included.

*Table of contents*


| | |
|---|---|
| *Alistipes onderdonkii A* | *1* |
| *Alistipes onderdonkii B* | *2* |
| *Alistipes putredinis A* | *3* |
| *Bacteroides cellulosilyticus A* | *3* |
| *Bacteroides massiliensis A* | *4* |
| *Bacteroides massiliensis B* | *5* |
| *Bacteroides ovatus A* | *6* |
| *Bacteroides ovatus B* | *7* |
| *Bacteroides thetaiotaomicron A* | *8* |
| *Bacteroides uniformis A* | *9* |
| *Bacteroides vulgatus A* | *10* |
| *Eubacterium rectale A* | *11* |
| *Eubacterium rectale B* | *12* |

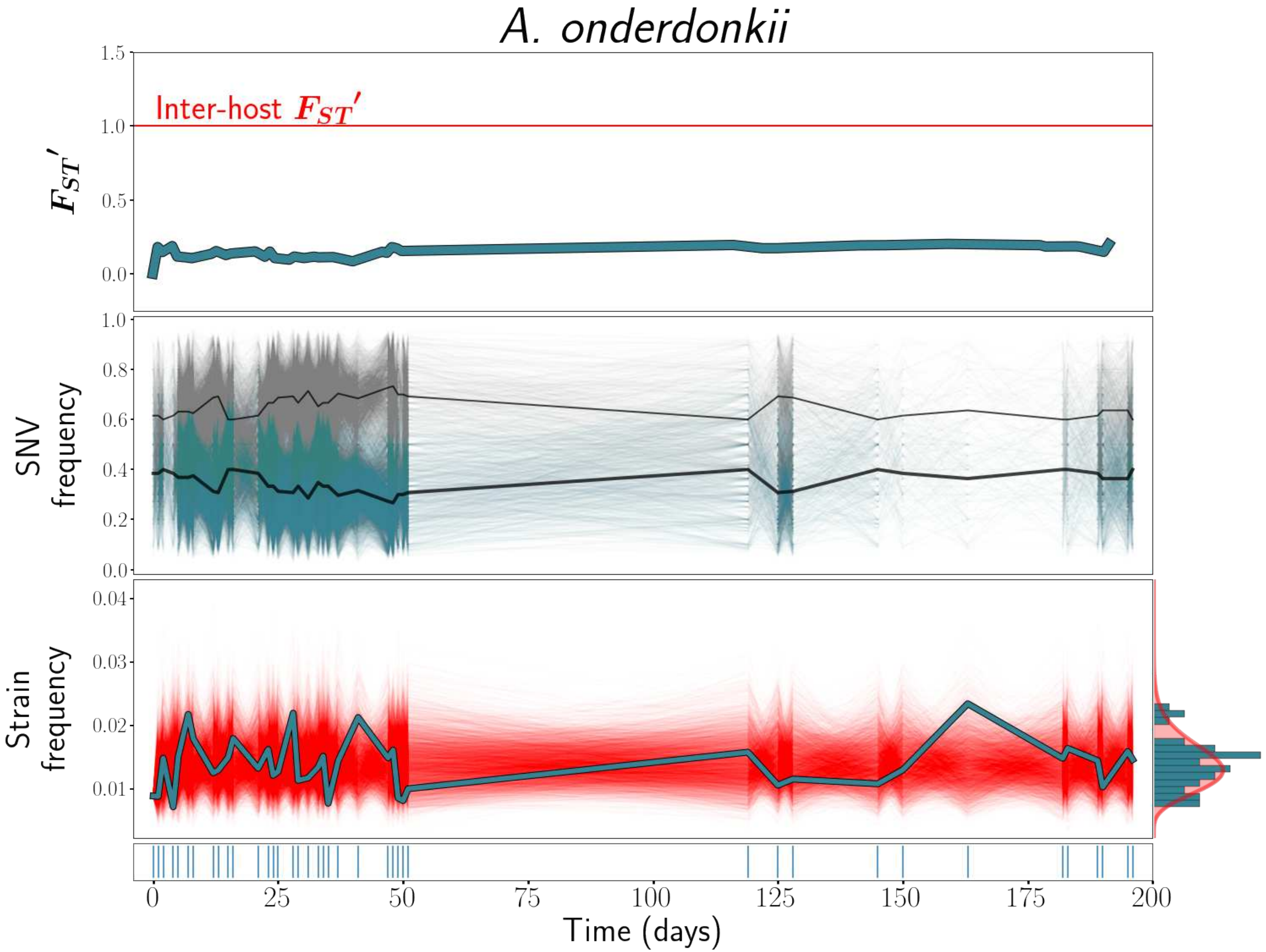

A. onderdonkii
1.5
Inter-host $F_{ST}'$
1.0
$F_{ST}'$
0.5
0.0
1.0
0.8
SNV frequency
0.6
0.4
0.2
0.0
0.04
0.03
Strain frequency
0.02
0.01
0
25
50
75
100
125
150
175
200
Time (days)

### *A. onderdonkii*

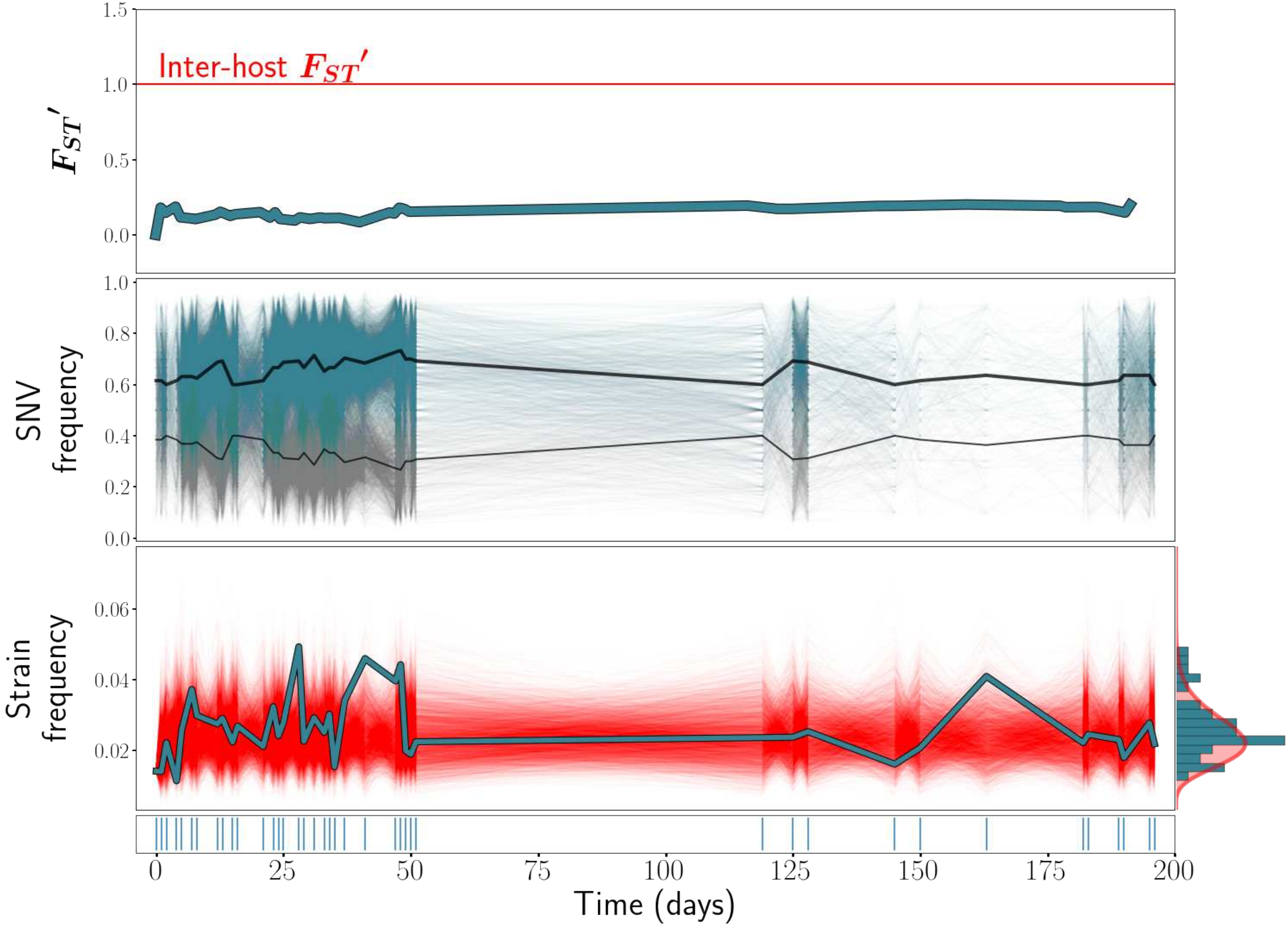

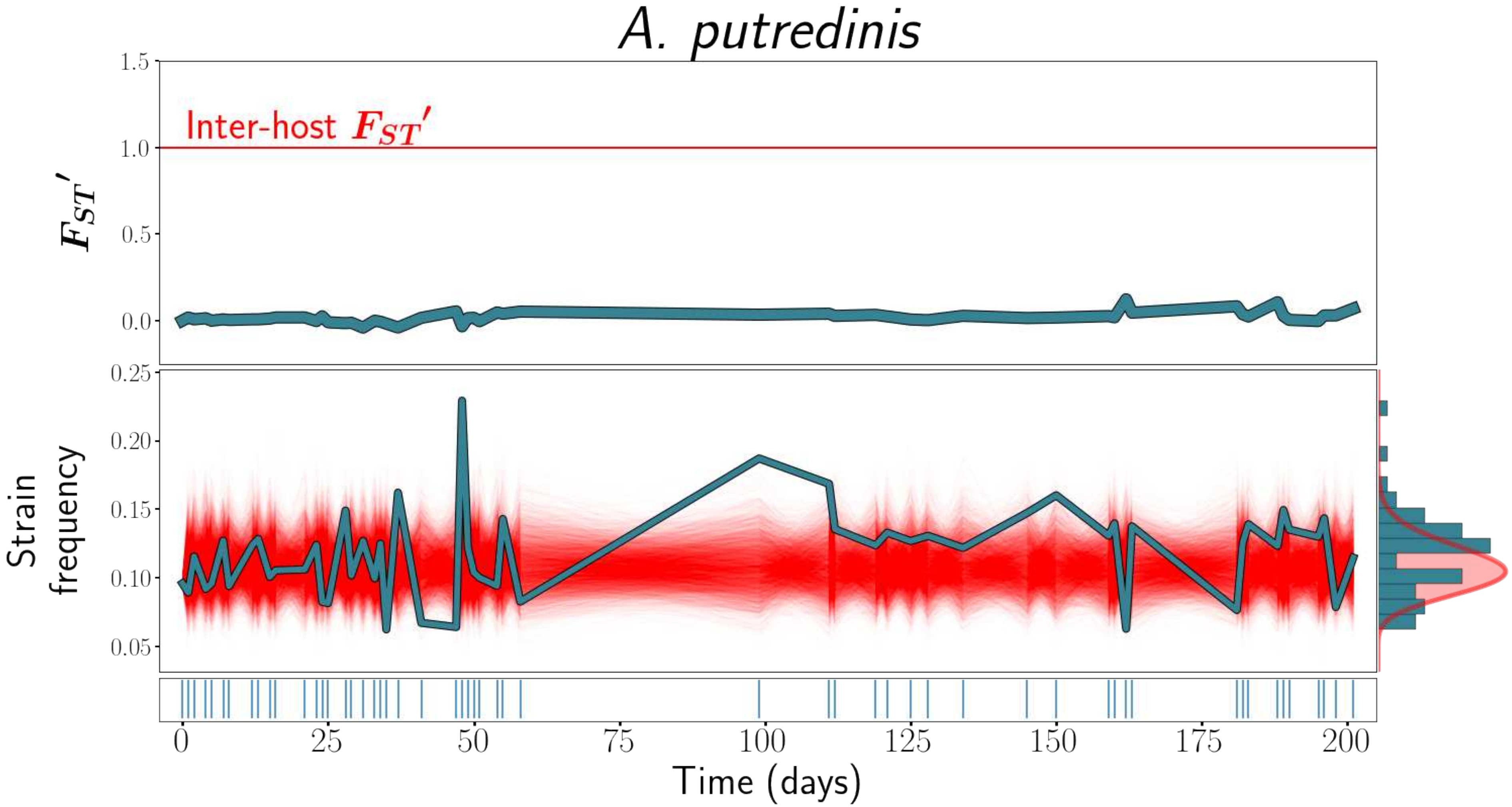

A. putredinis
Inter-host $F_{ST}'$
$F_{ST}'$
1.5
1.0
0.5
0.0
Strain frequency
0.25
0.20
0.15
0.10
0.05
0
25
50
75
100
125
150
175
200
Time (days)

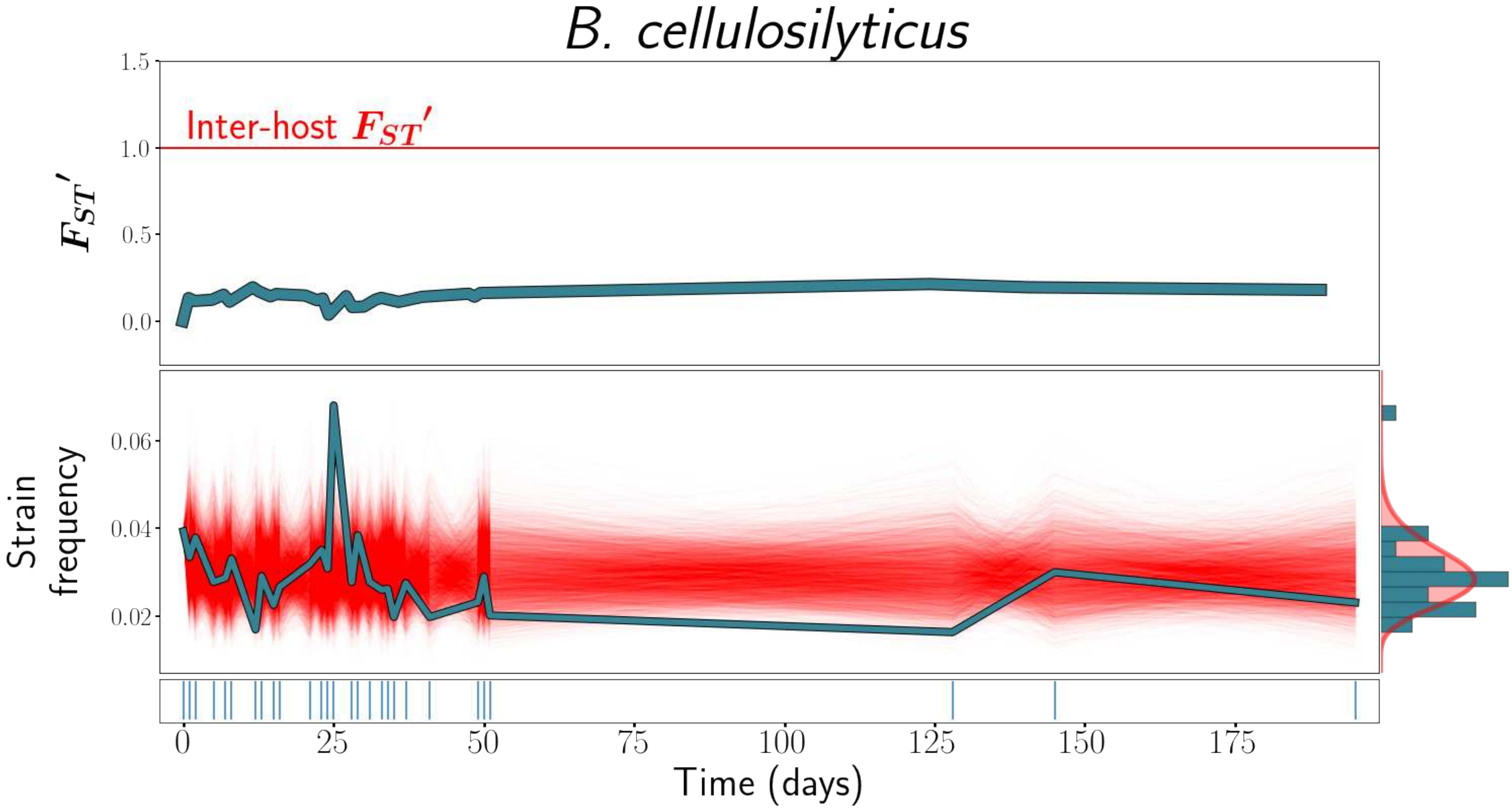
B. cellulosilyticus
Inter-host $F_{ST}'$
$F_{ST}'$
1.5
1.0
0.5
0.0
Strain frequency
0.06
0.04
0.02
0
25
50
75
100
125
150
175
Time (days)

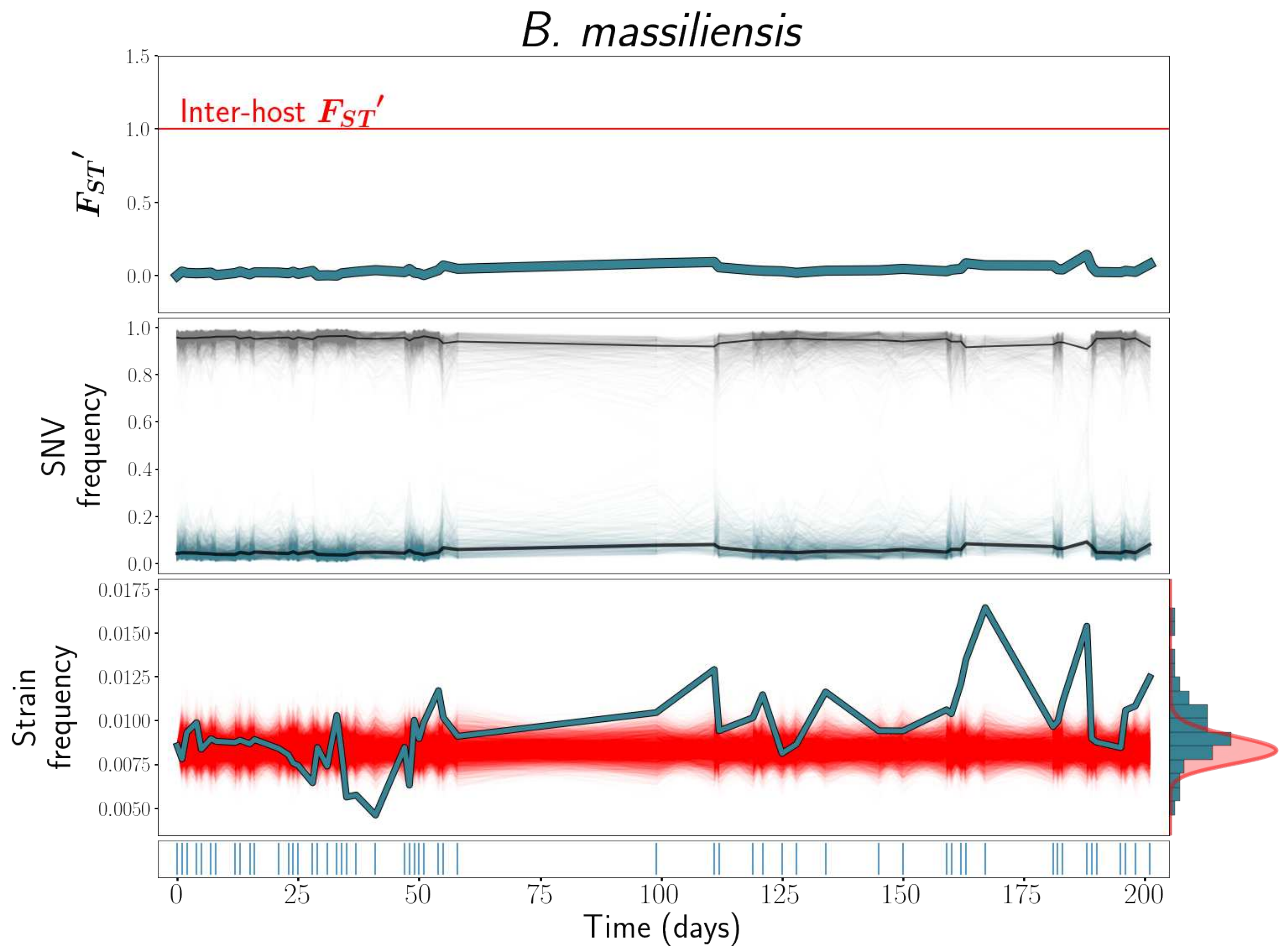

B. massiliensis
Inter-host $F_{ST}'$
$F_{ST}'$
1.5
1.0
0.5
0.0
SNV frequency
1.0
0.8
0.6
0.4
0.2
0.0
Strain frequency
0.0175
0.0150
0.0125
0.0100
0.0075
0.0050
0
25
50
75
100
125
150
175
200
Time (days)

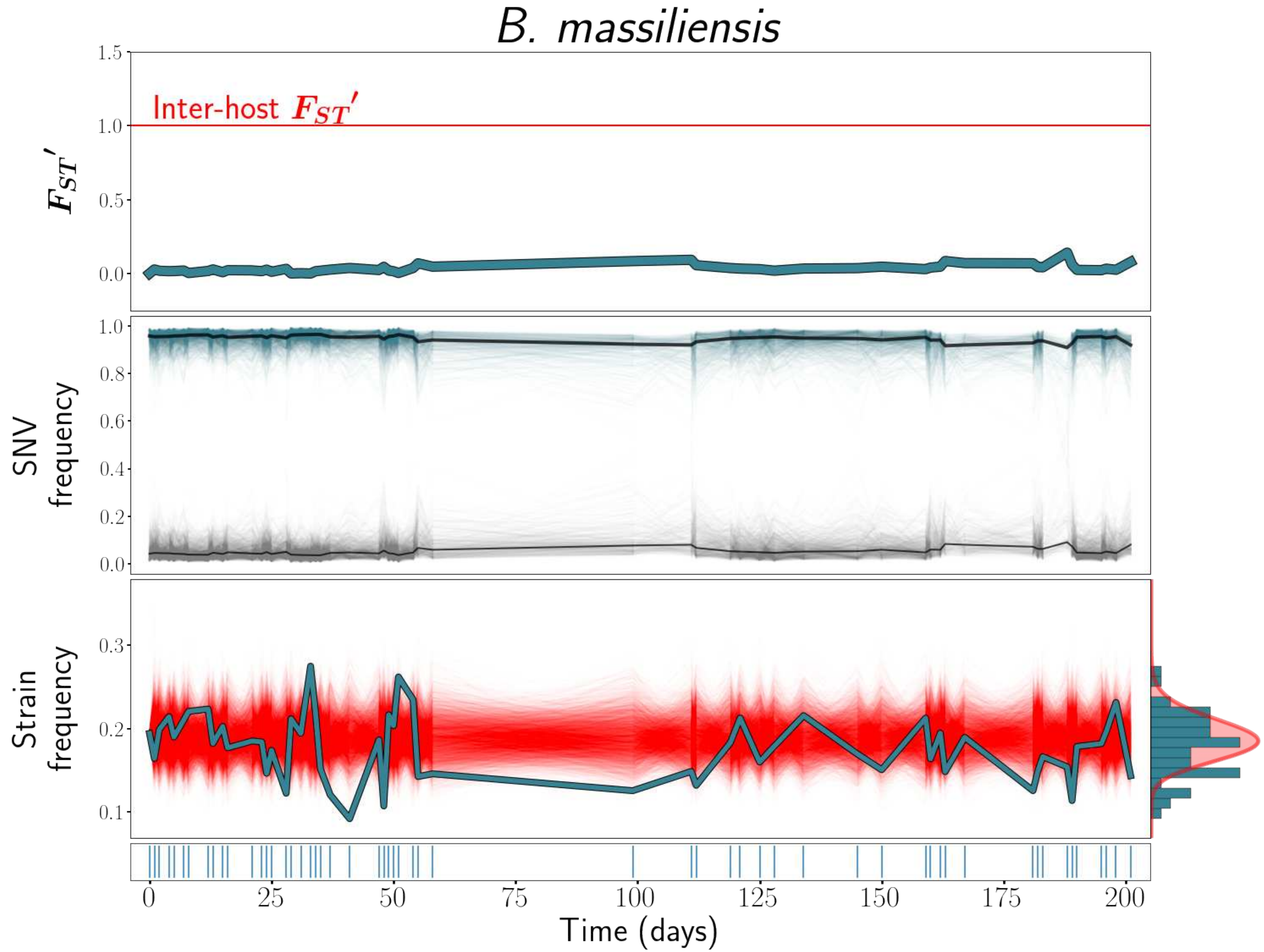

B. massiliensis
Inter-host $F_{ST}'$
$F_{ST}'$
SNV frequency
Strain frequency
Time (days)

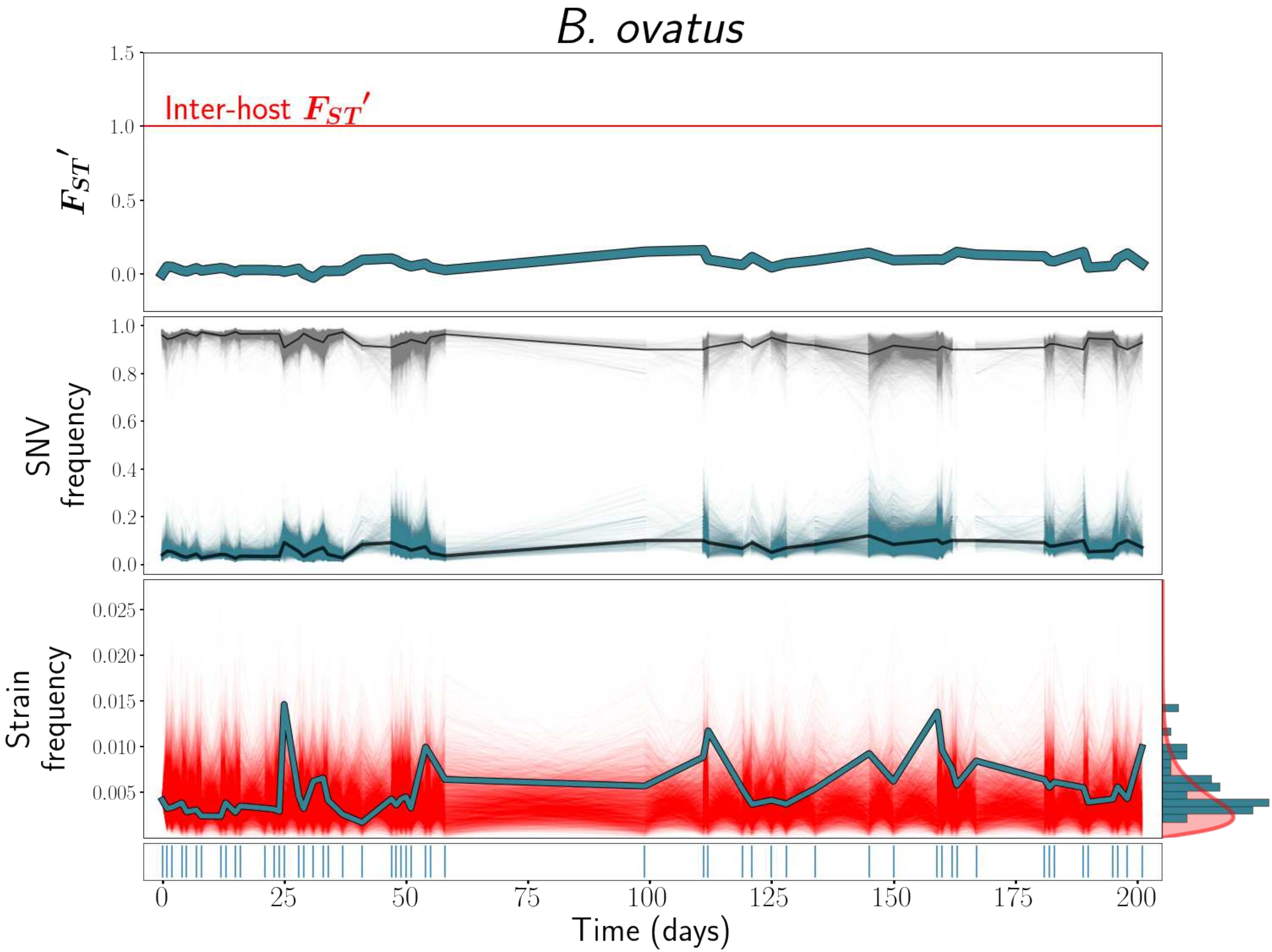

B. ovatus
Inter-host $F_{ST}'$
$F_{ST}'$
SNV frequency
Strain frequency
Time (days)

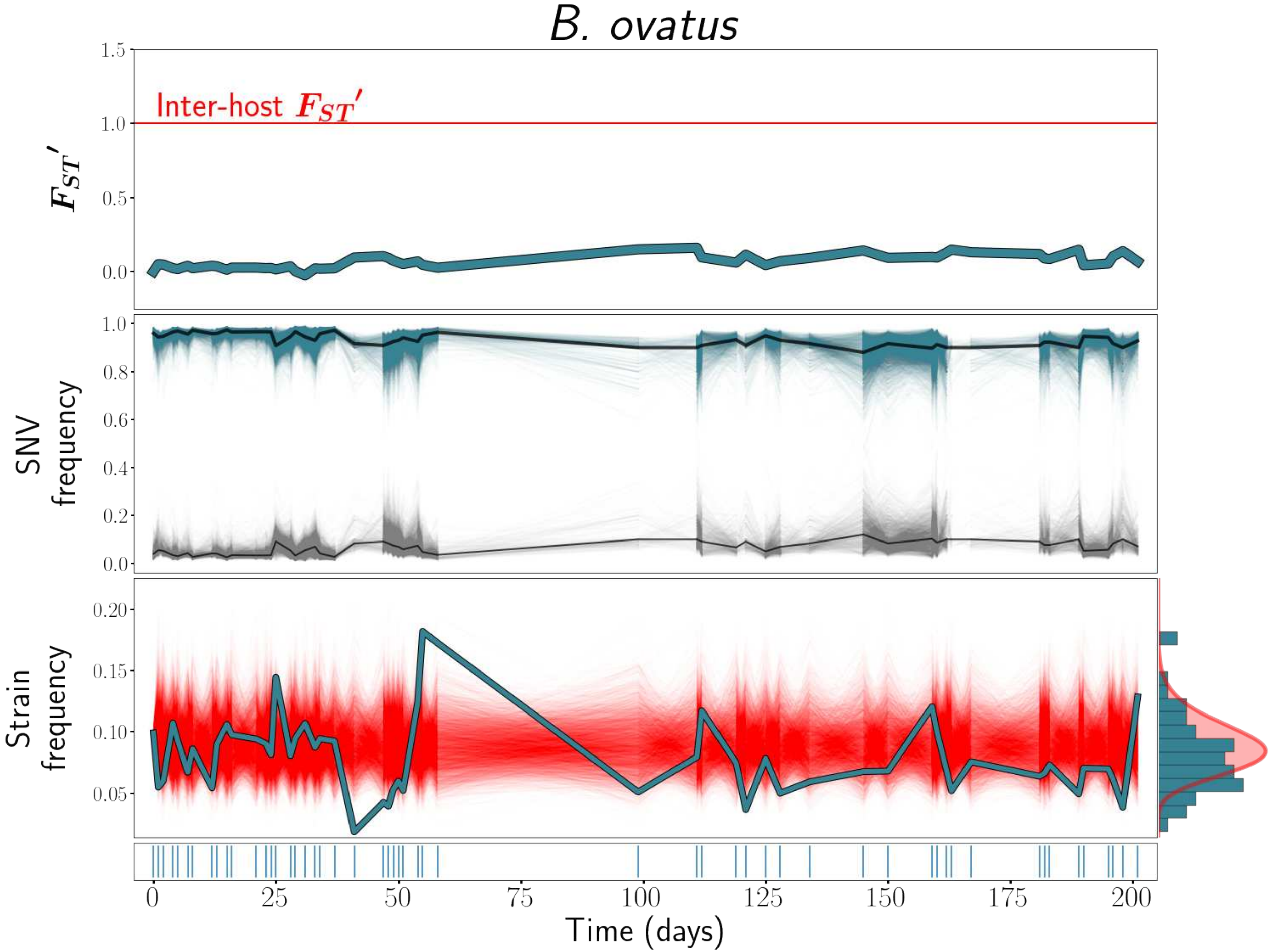
B. ovatus
Inter-host $F_{ST}'$
$F_{ST}'$
SNV frequency
Strain frequency
Time (days)
1.5
1.0
0.5
0.0
0.8
0.6
0.4
0.2
0.20
0.15
0.10
0.05
0
25
50
75
100
125
150
175
200

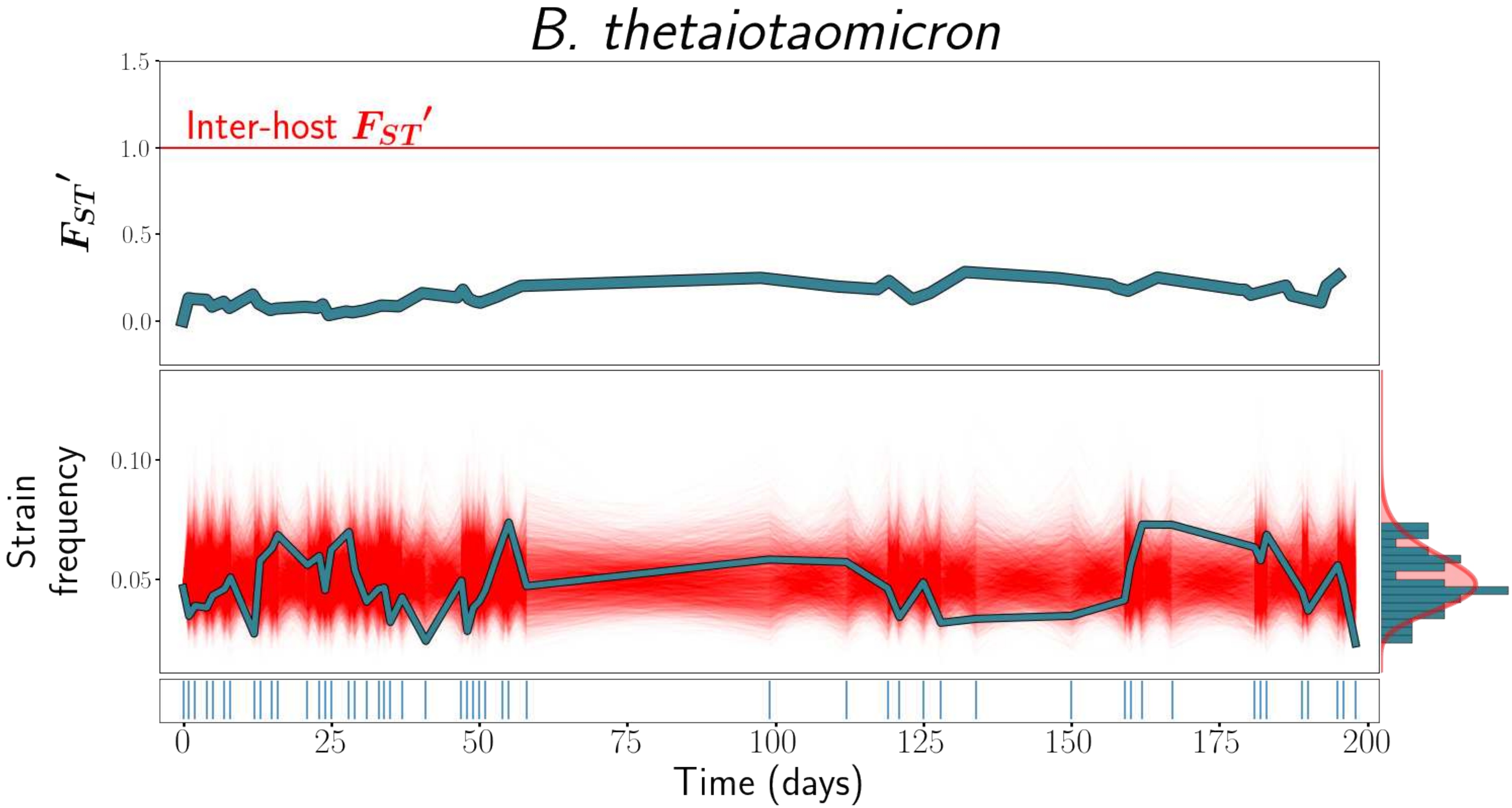

B. thetaiotaomicron
Inter-host $F_{ST}'$
$F_{ST}'$
1.5
1.0
0.5
0.0
Strain frequency
0.10
0.05
0
25
50
75
100
125
150
175
200
Time (days)

### *B. uniformis*

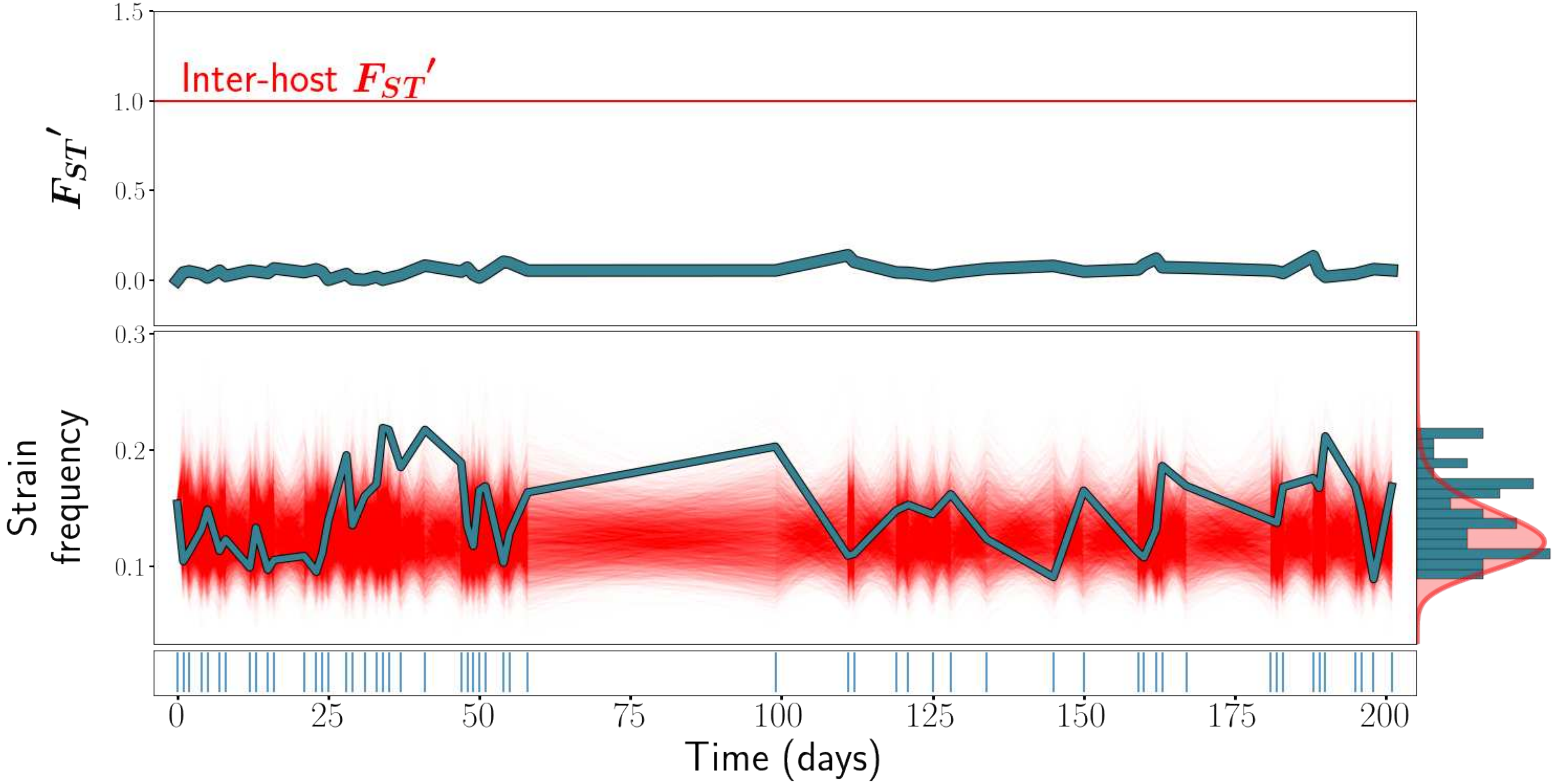

1.5
1.0
0.5
0.0
Inter-host $F_{ST}'$
$F_{ST}'$
0.3
0.2
0.1
Strain frequency
0
25
50
75
100
125
150
175
200
Time (days)

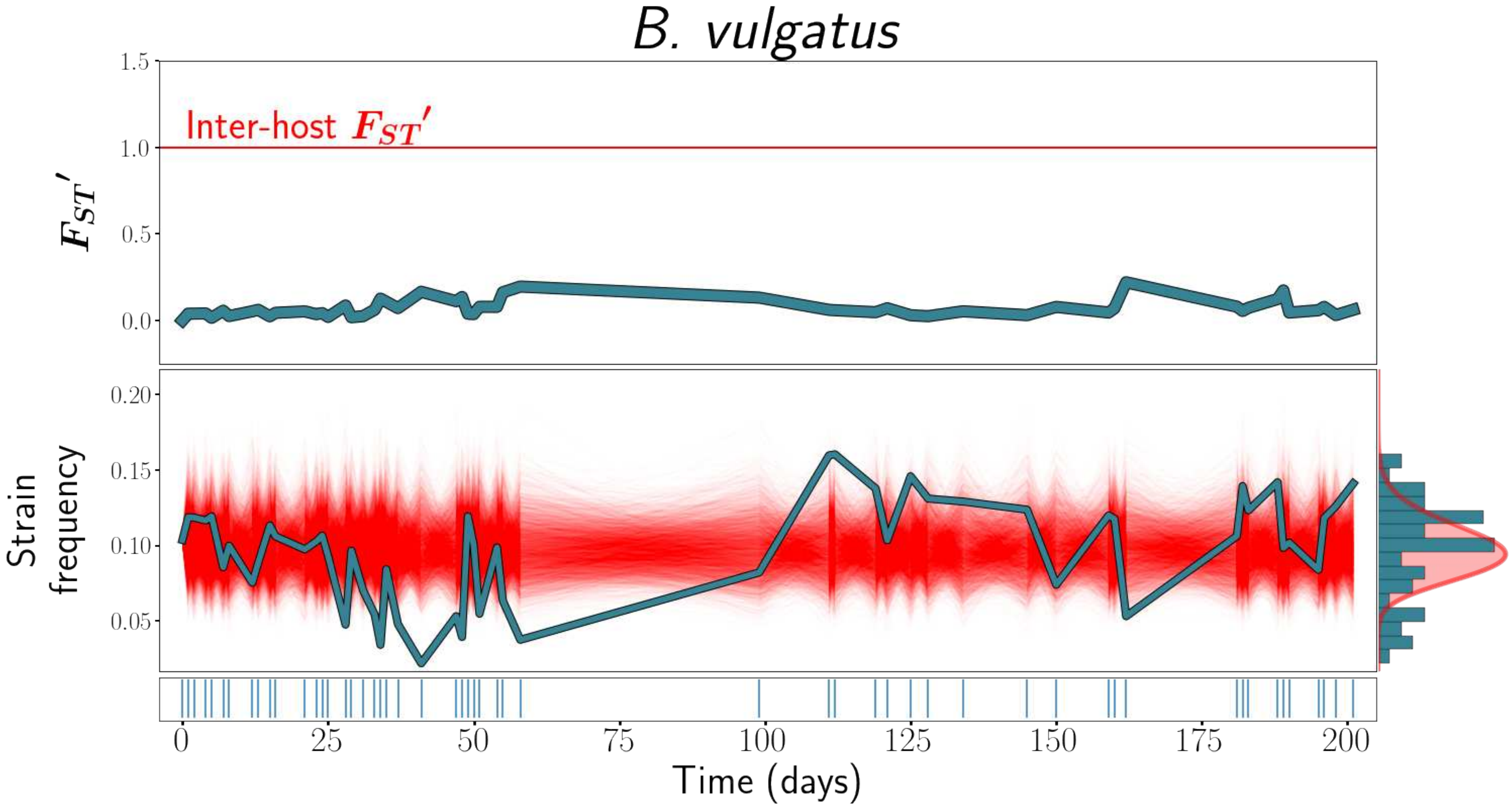
B. vulgatus
Inter-host $F_{ST}'$
$F_{ST}'$
1.5
1.0
0.5
0.0
Strain frequency
0.20
0.15
0.10
0.05
0
25
50
75
100
125
150
175
200
Time (days)

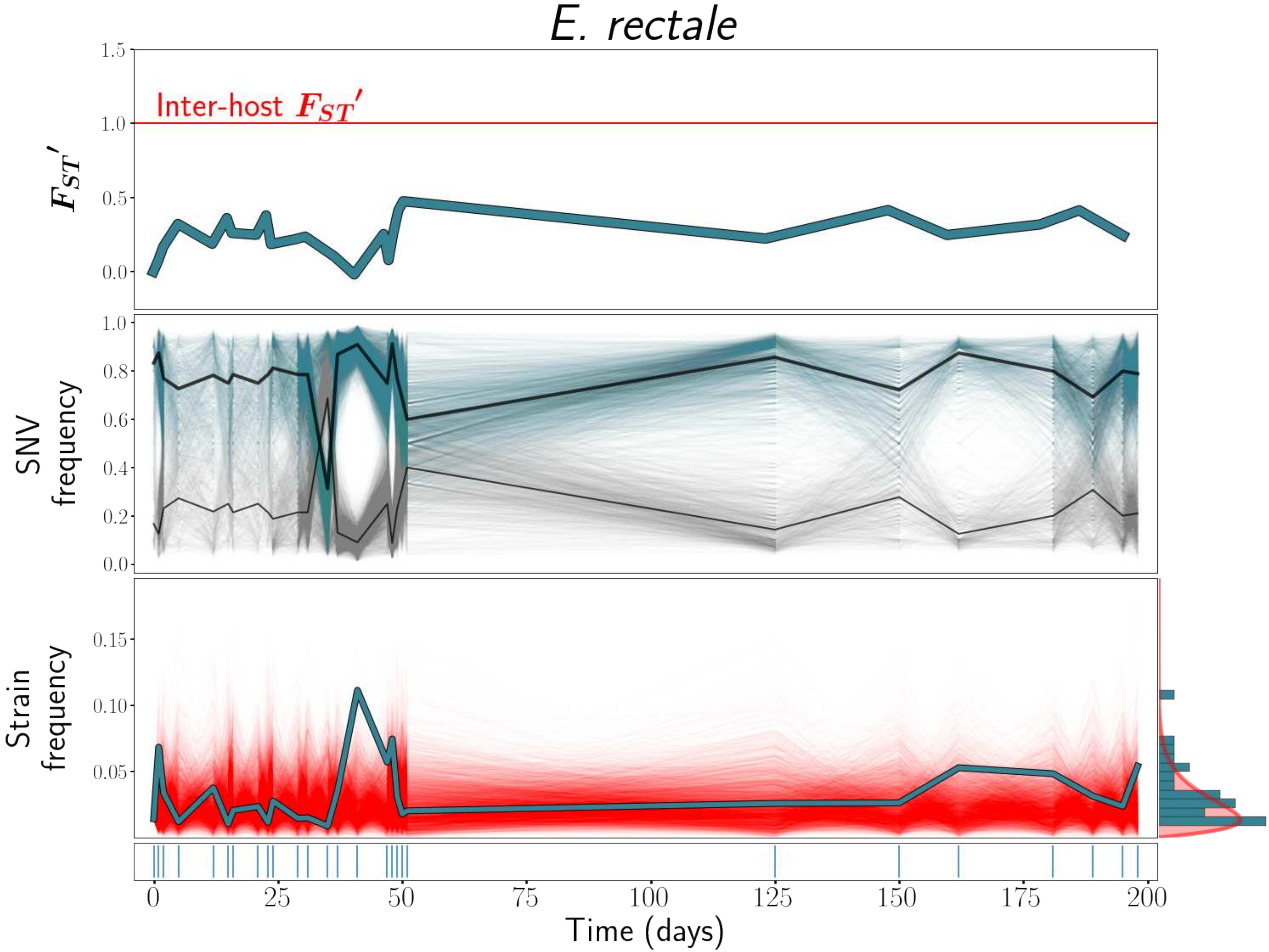

E. rectale
Inter-host $F_{ST}'$
$F_{ST}'$
1.5
1.0
0.5
0.0
SNV frequency
1.0
0.8
0.6
0.4
0.2
0.0
Strain frequency
0.15
0.10
0.05
0
25
50
75
100
125
150
175
200
Time (days)

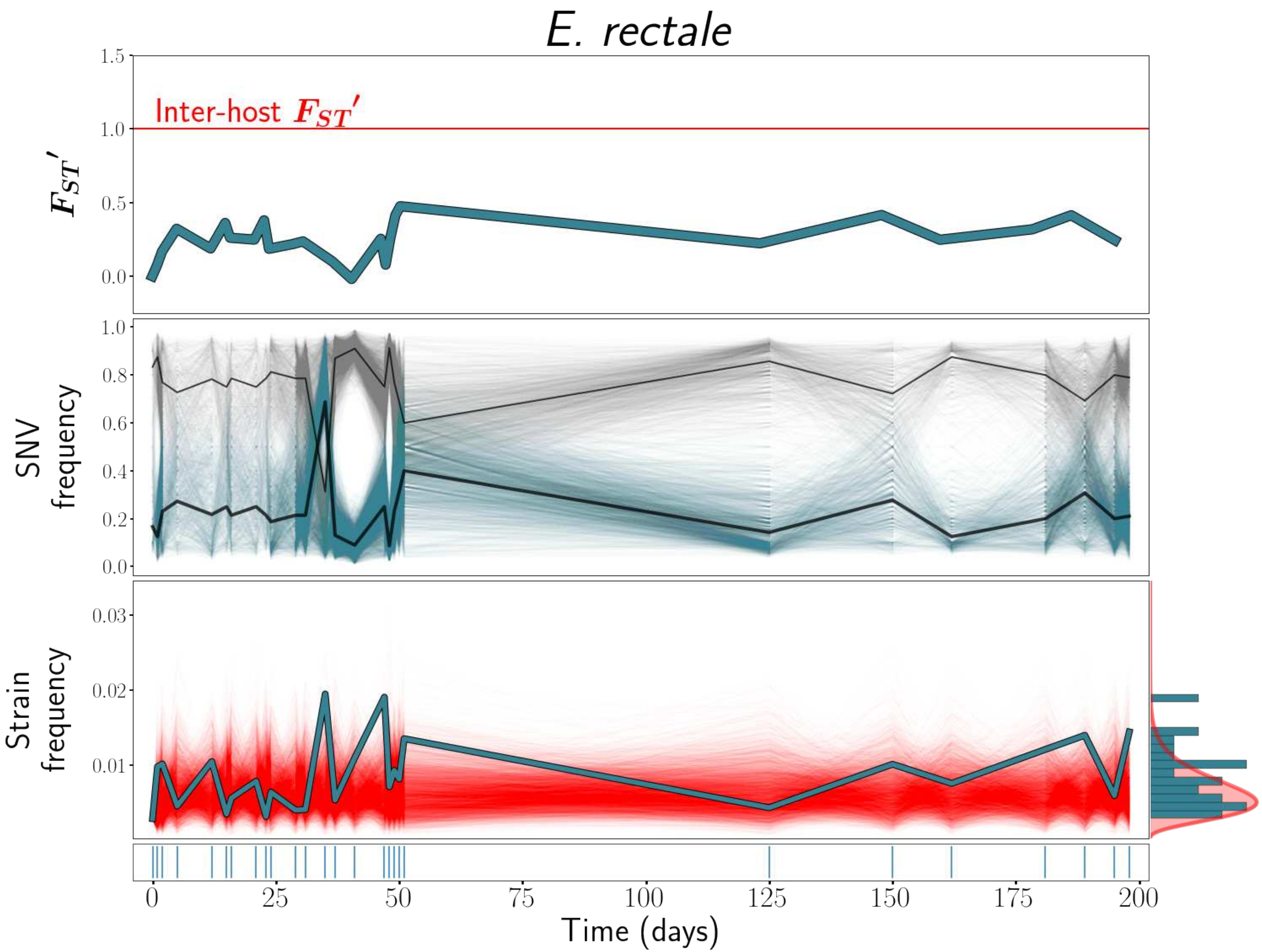

E. rectale
Inter-host $F_{ST}'$
$F_{ST}'$
SNV frequency
Strain frequency
Time (days)
