## Supplementary material for "Ecological Stability Emerges at the Level of Strains in the Human Gut Microbiome": S5 Text

### S3 Text

$F_{ST}$', strain frequency, and strain abundance dynamics plots for all species analyzed, for host *ao.* These plots are analogous to Main Text Figure 1. When only a single strain was detected, only $F_{ST}$' and strain abundance dynamics plots, but no strain frequency plot, is included.

*Table of contents*


| | |
|---|---|
| *Bacteroides fragilis A* | *1* |
| *Bacteroides ovatus A* | *2* |
| *Bacteroides ovatus B* | *3* |
| *Bacteroides uniformis A* | *4* |
| *Bacteroides xylanisolvens A* | *5* |
| *Bacteroides xylanisolvens B* | *6* |
| *Bifidobacterium adolescentis* | *7* |
| *Dialister invisus A* | *8* |
| *Eubacterium eligens A* | *9* |
| *Eubacterium rectale A* | *10* |
| *Faecalibacterium prausnitzii (61481) A* | *11* |
| *Faecalibacterium prausnitzii (61481) B* | *12* |
| *Faecalibacterium prausnitzii (62201) A* | *13* |
| *Ruminoccocus bicirculans A* | *14* |
| *Parabacteroides distasonis A* | *15* |

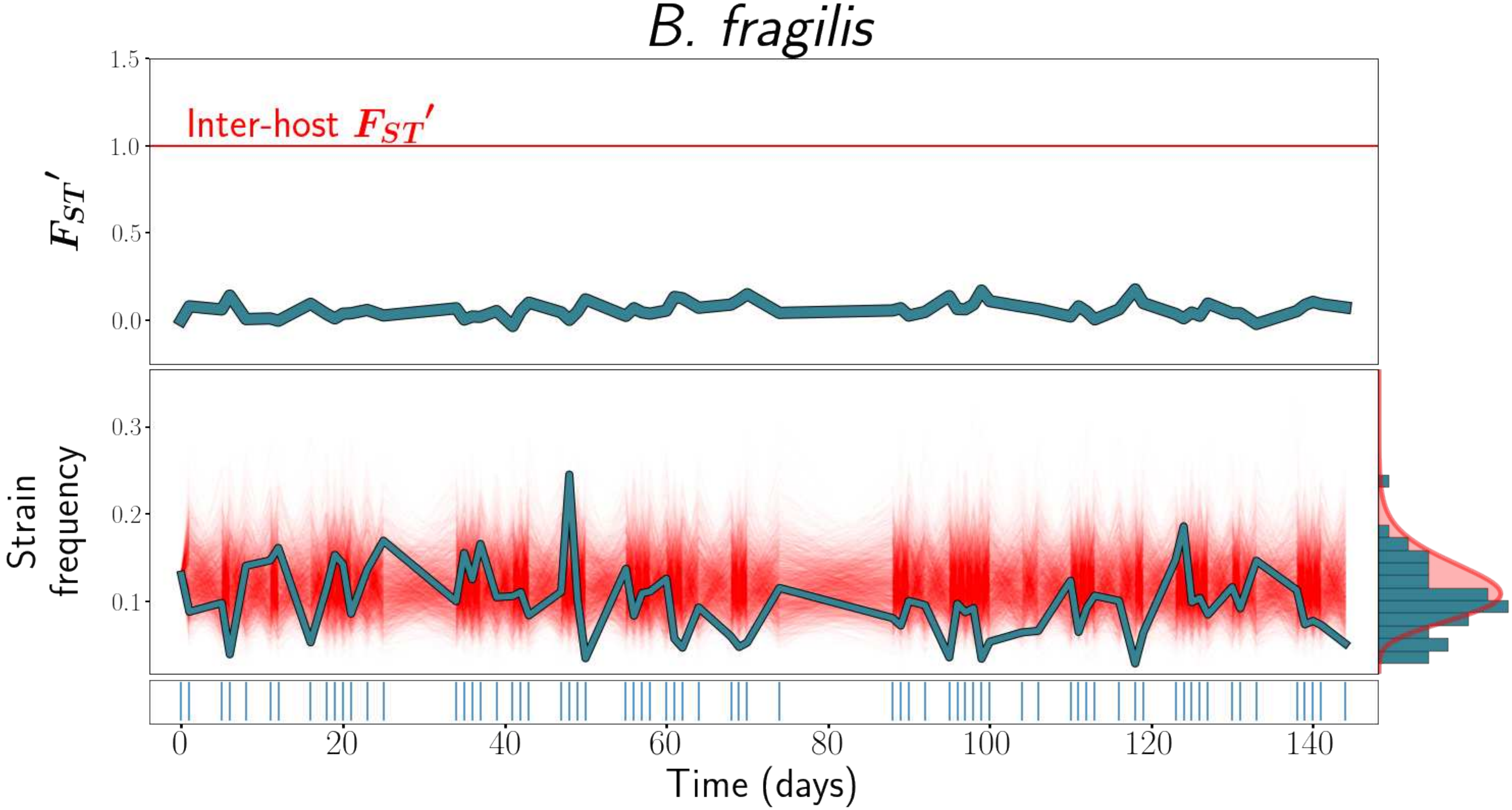
B. fragilis
Inter-host $F_{ST}'$
$F_{ST}'$
1.5
1.0
0.5
0.0
Strain frequency
0.3
0.2
0.1
0
20
40
60
80
100
120
140
Time (days)

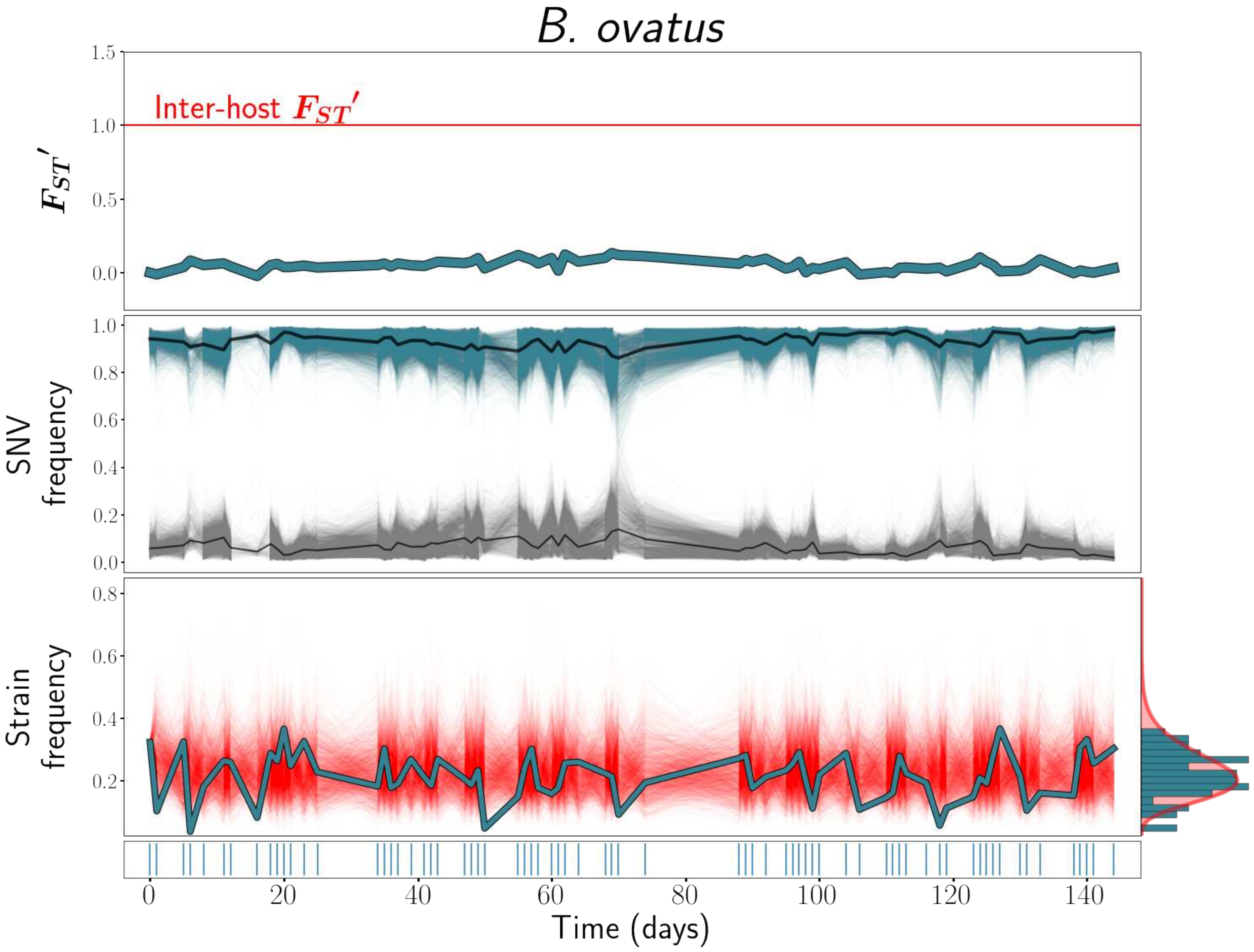

B. ovatus
Inter-host $F_{ST}'$
$F_{ST}'$
1.5
1.0
0.5
0.0
SNV frequency
1.0
0.8
0.6
0.4
0.2
0.0
Strain frequency
0.8
0.6
0.4
0.2
0
20
40
60
80
100
120
140
Time (days)

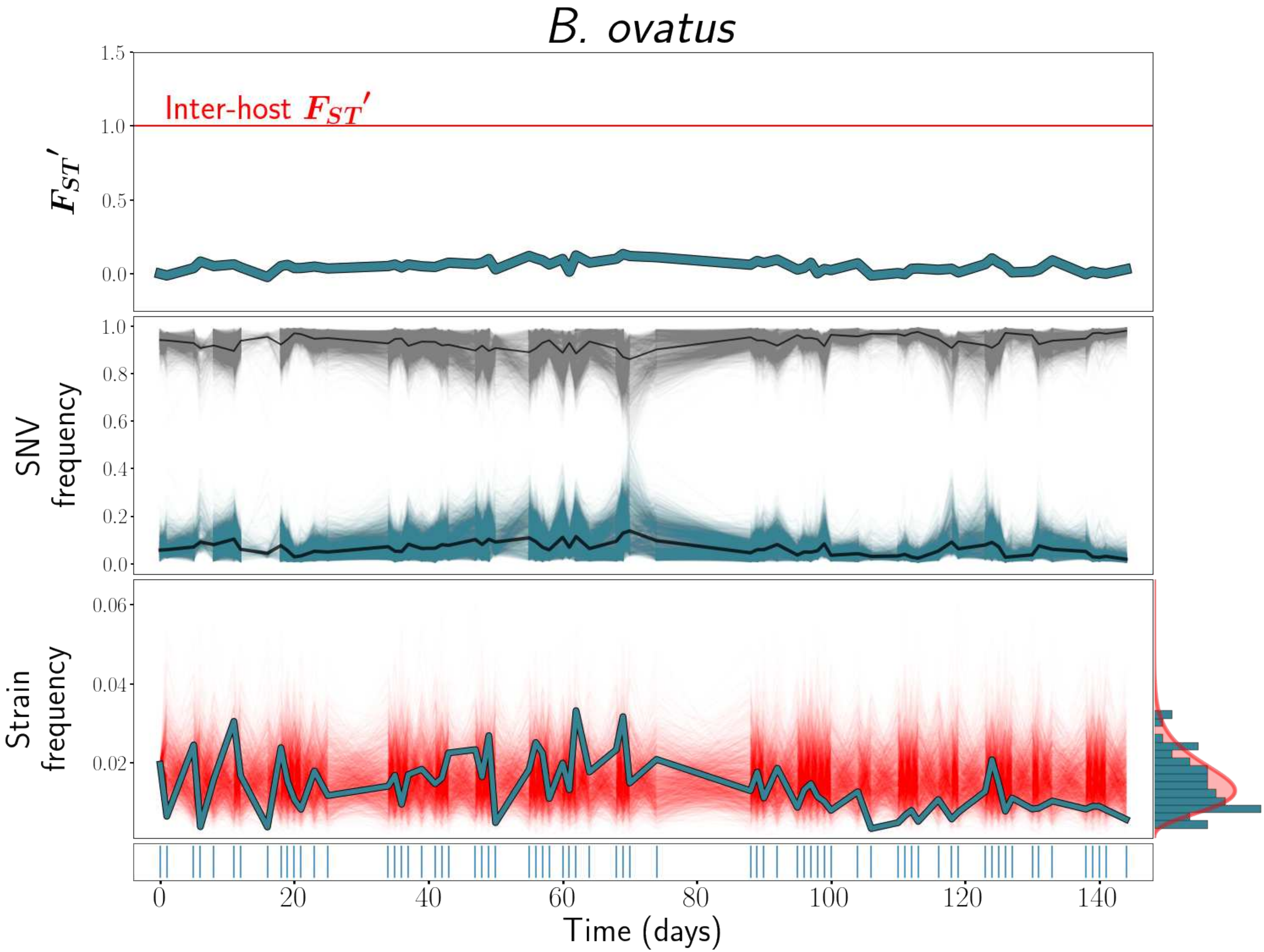

B. ovatus
Inter-host $F_{ST}'$
$F_{ST}'$
SNV frequency
Strain frequency
Time (days)

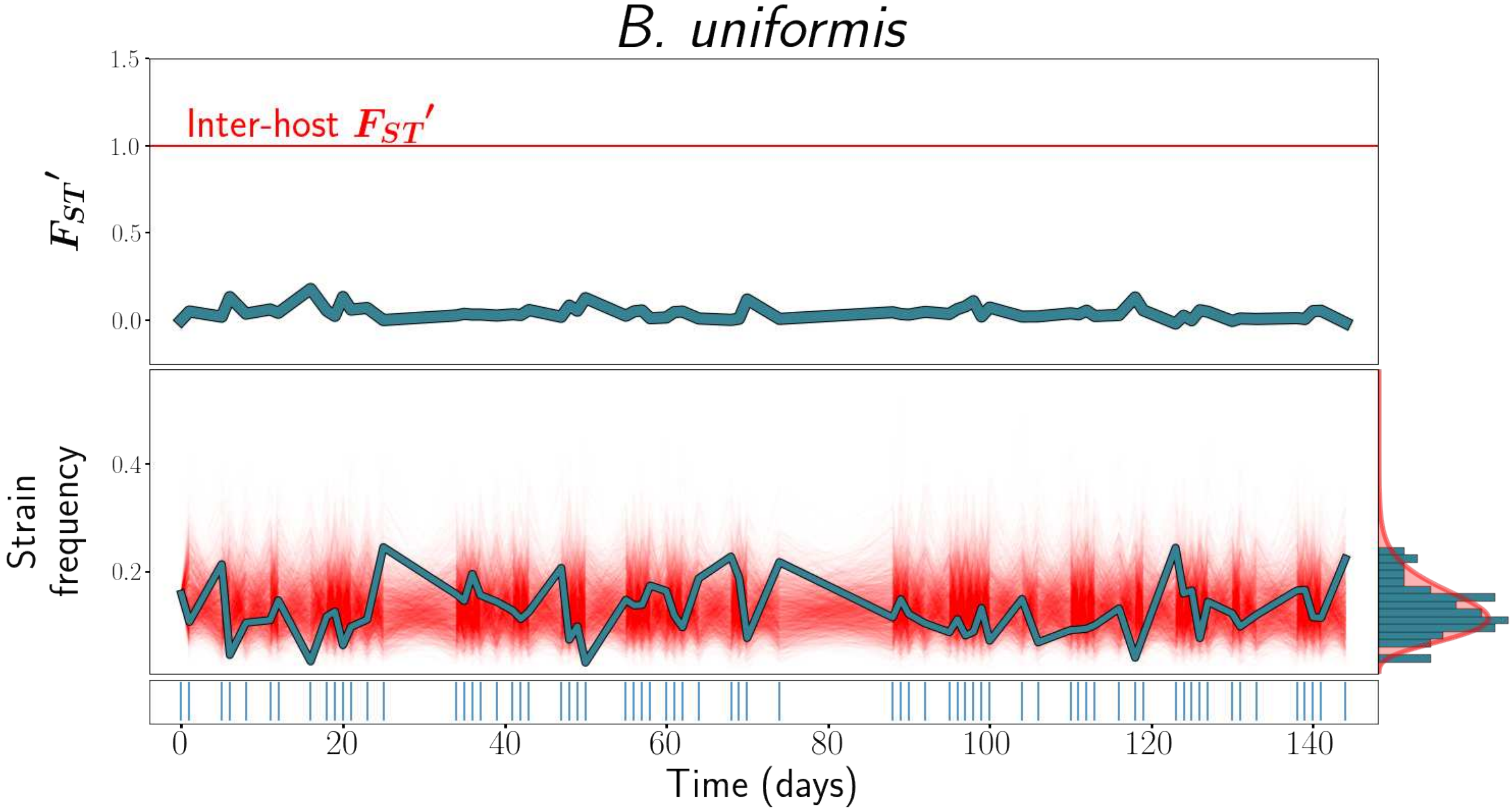

B. uniformis
Inter-host $F_{ST}'$
1.5
1.0
0.5
0.0
$F_{ST}'$
0.4
0.2
Strain frequency
0
20
40
60
80
100
120
140
Time (days)

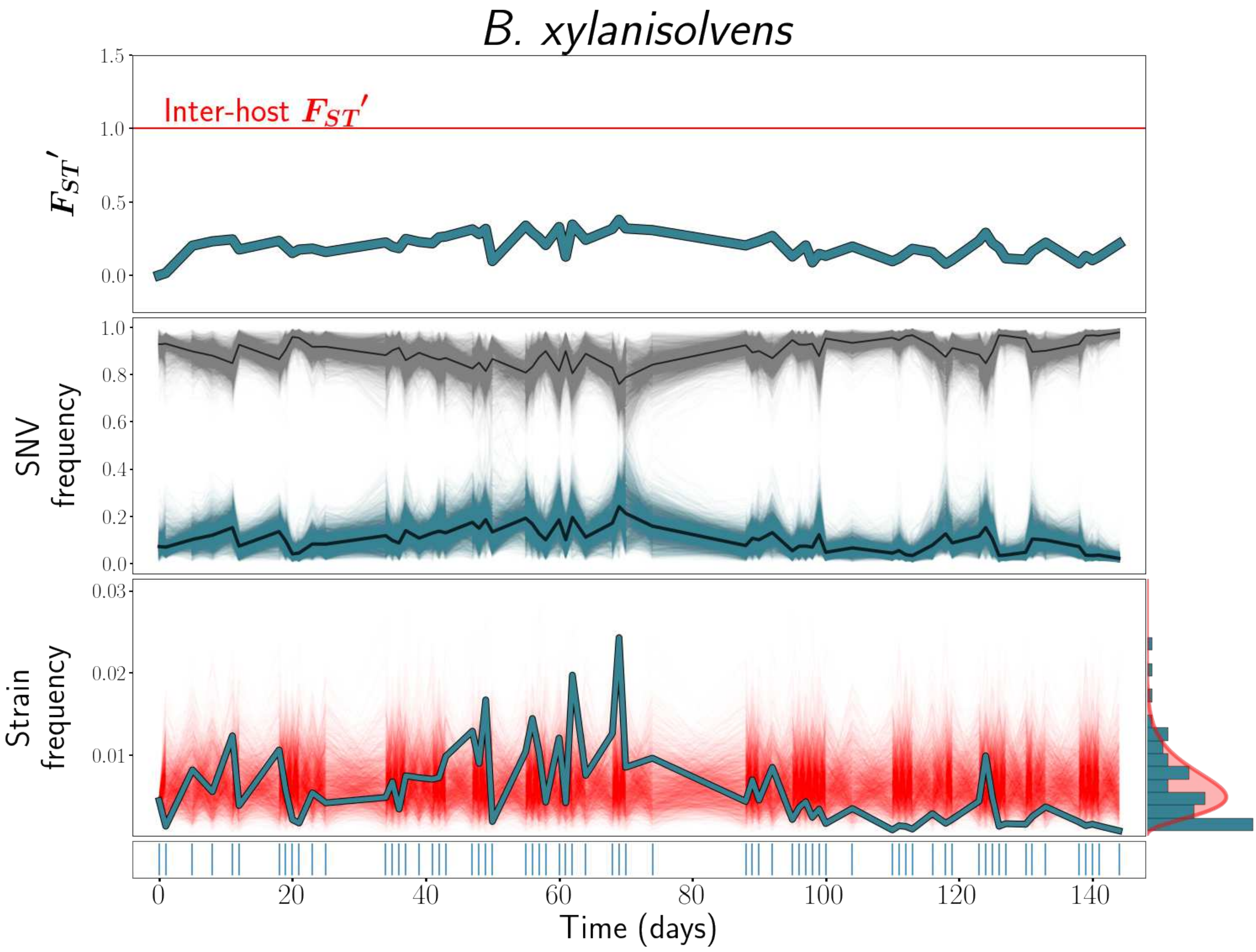

B. xylanisolvens
Inter-host $F_{ST}'$
$F_{ST}'$
SNV frequency
Strain frequency
Time (days)

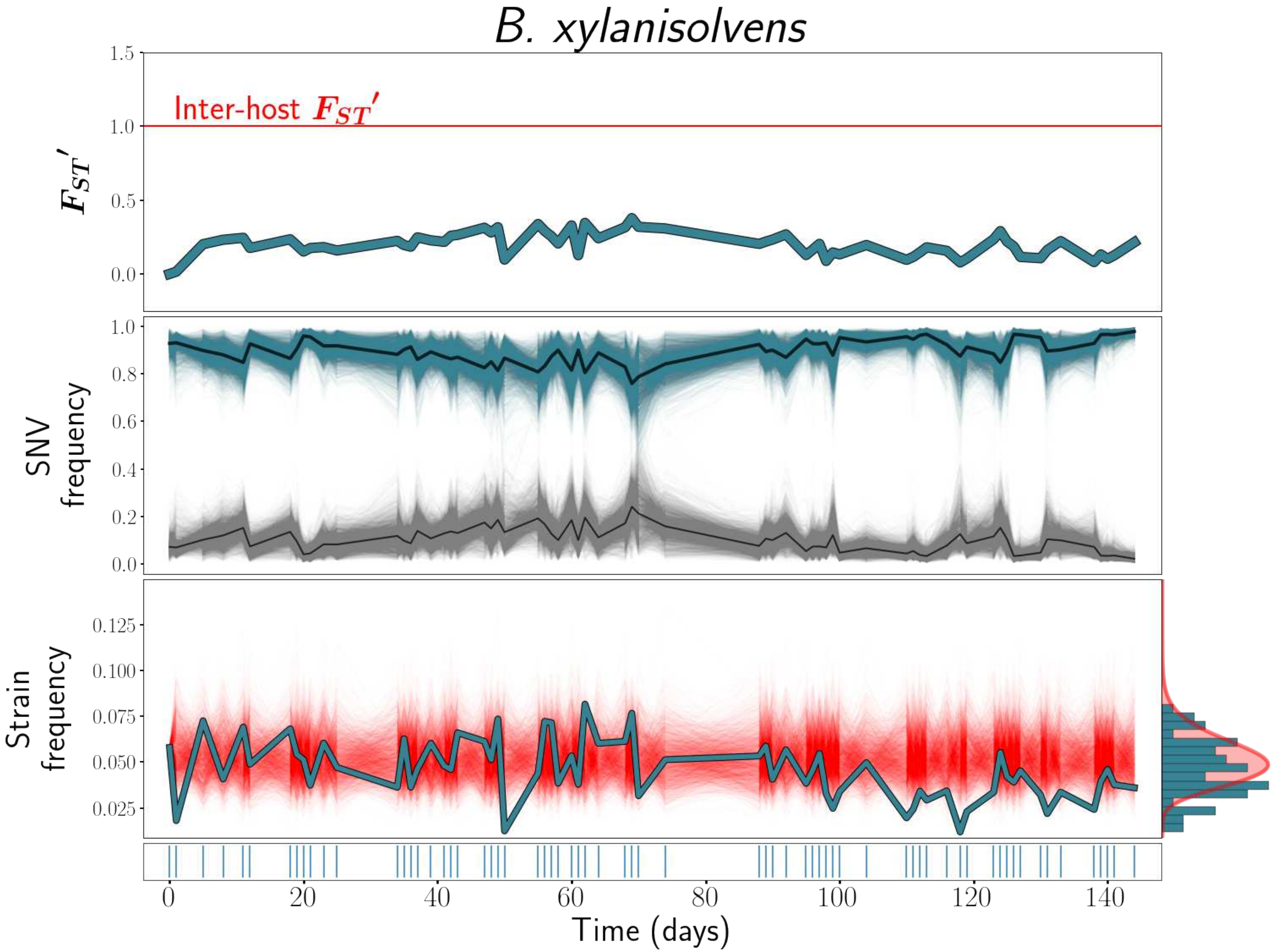

B. xylanisolvens
Inter-host $F_{ST}'$
$F_{ST}'$
1.5
1.0
0.5
0.0
SNV frequency
1.0
0.8
0.6
0.4
0.2
0.0
Strain frequency
0.125
0.100
0.075
0.050
0.025
0
20
40
60
80
100
120
140
Time (days)

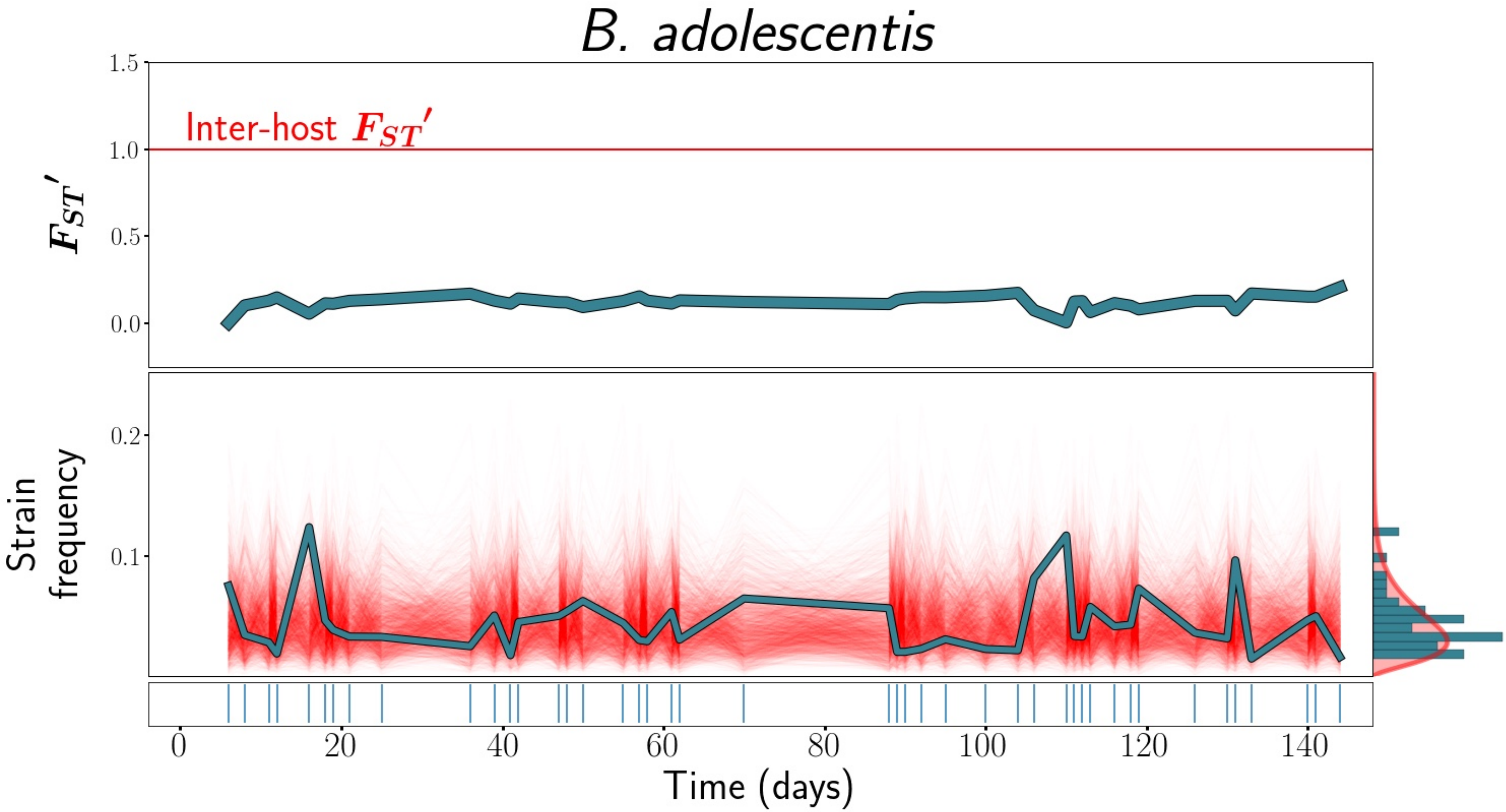

B. adolescentis
Inter-host $F_{ST}'$
$F_{ST}'$
1.5
1.0
0.5
0.0
Strain frequency
0.2
0.1
0
20
40
60
80
100
120
140
Time (days)

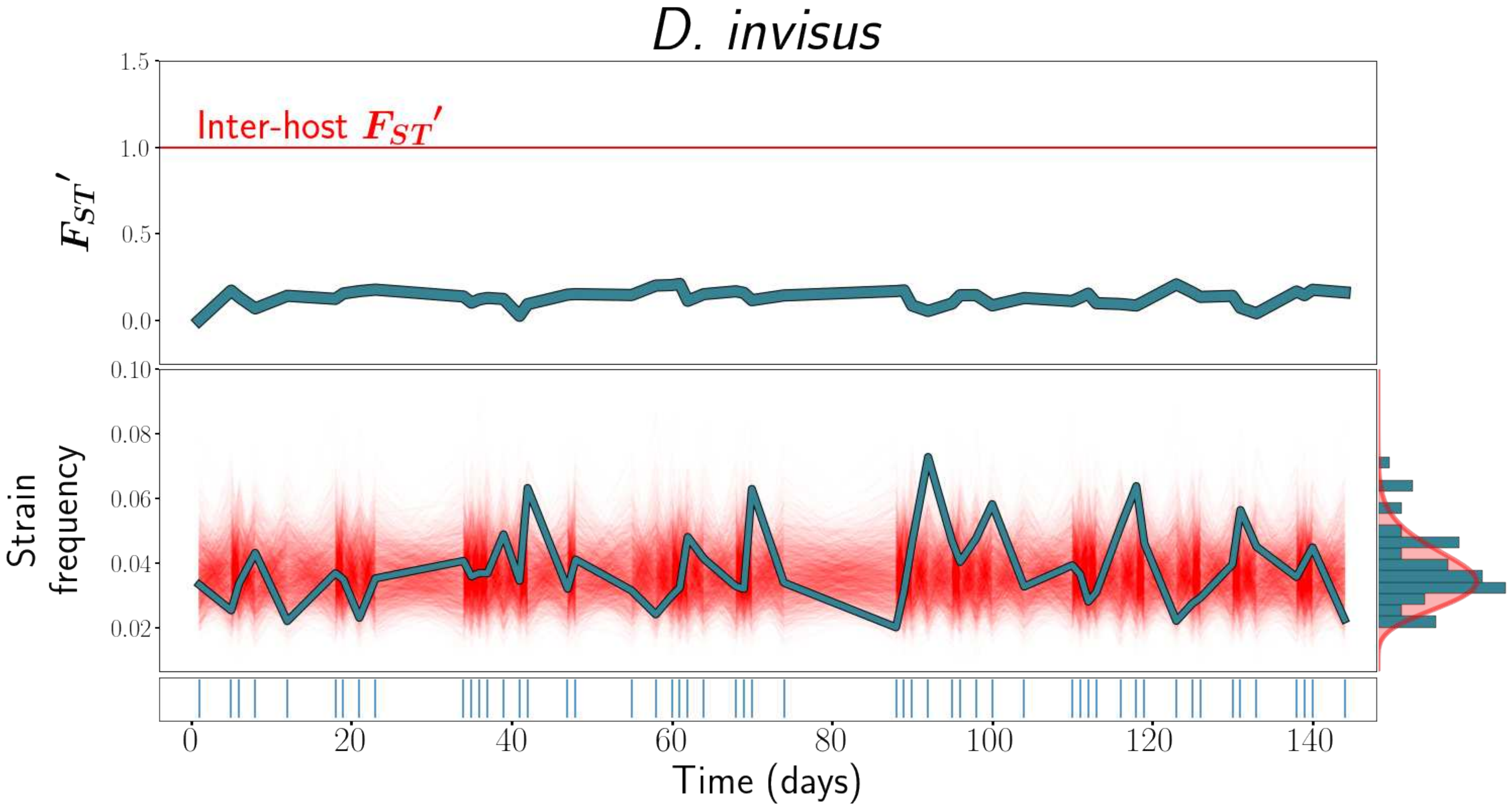

D. invisus
Inter-host $F_{ST}'$
$F_{ST}'$
1.5
1.0
0.5
0.0
0.10
0.08
0.06
0.04
0.02
Strain frequency
0
20
40
60
80
100
120
140
Time (days)

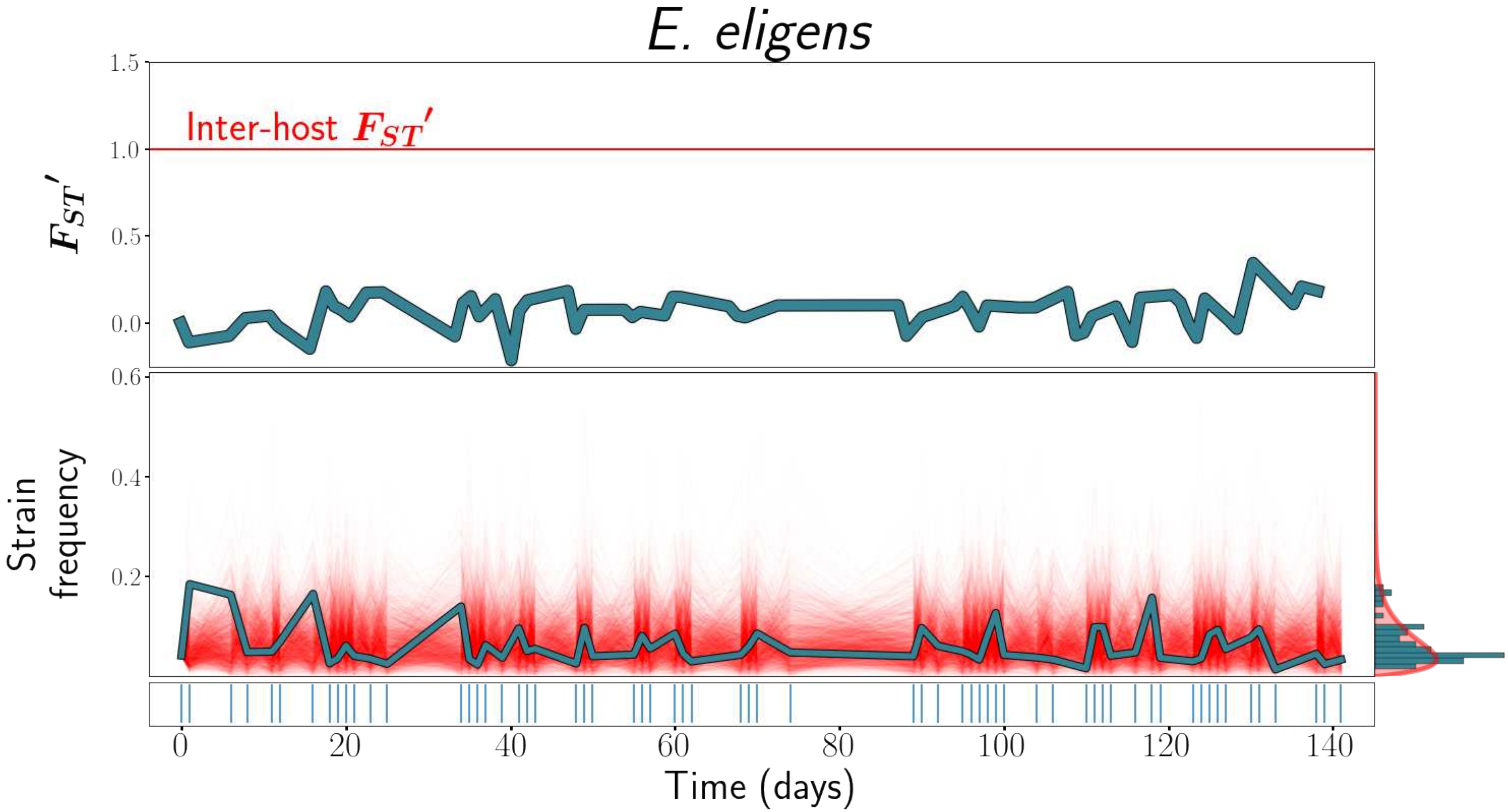
E. eligens
1.5
1.0
0.5
0.0
Inter-host $F_{ST}'$
$F_{ST}'$
0.6
0.4
0.2
Strain frequency
0
20
40
60
80
100
120
140
Time (days)

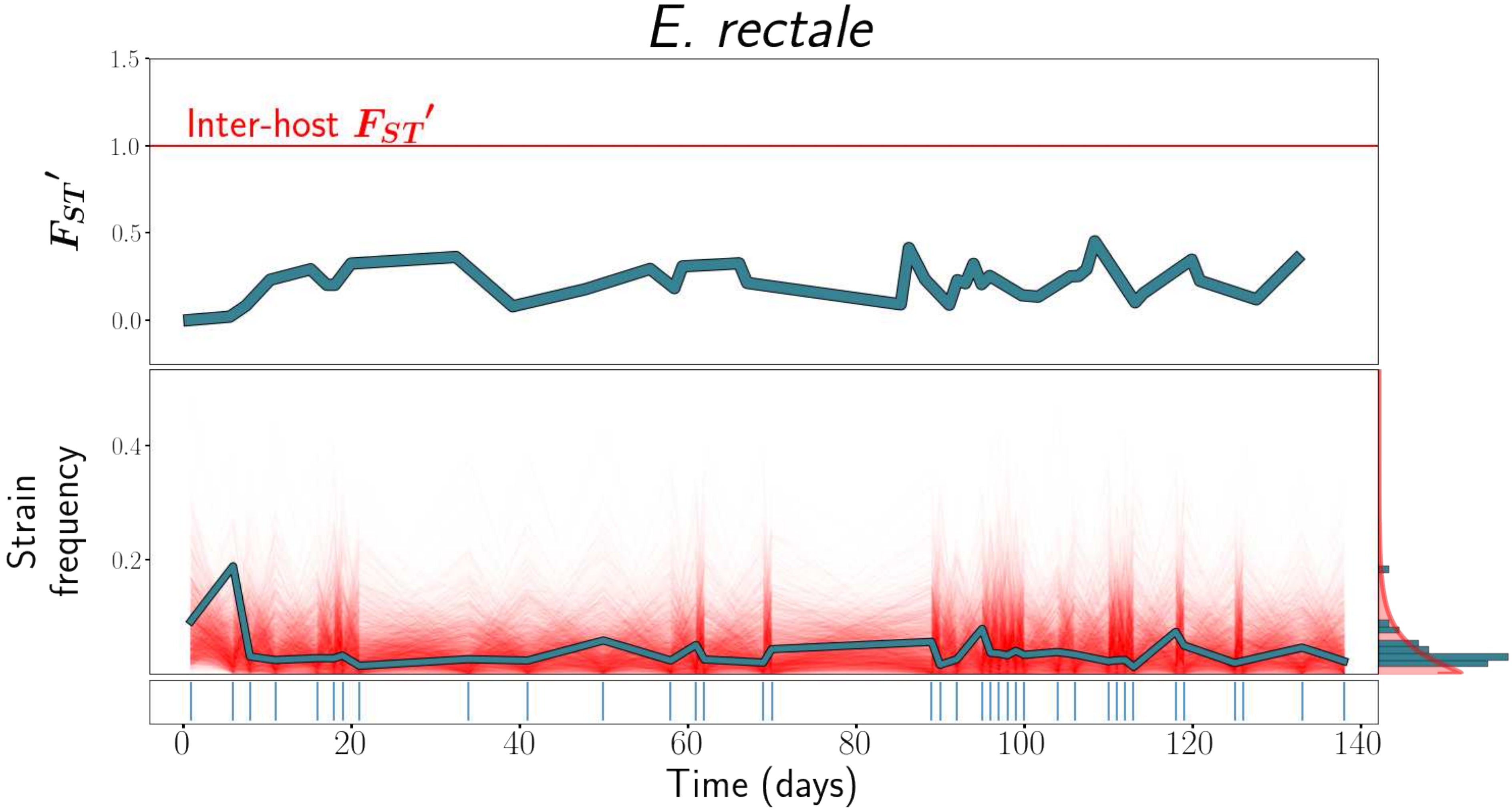

E. rectale
1.5
Inter-host $F_{ST}'$
1.0
$F_{ST}'$
0.5
0.0
0.4
Strain frequency
0.2
0
20
40
60
80
100
120
140
Time (days)

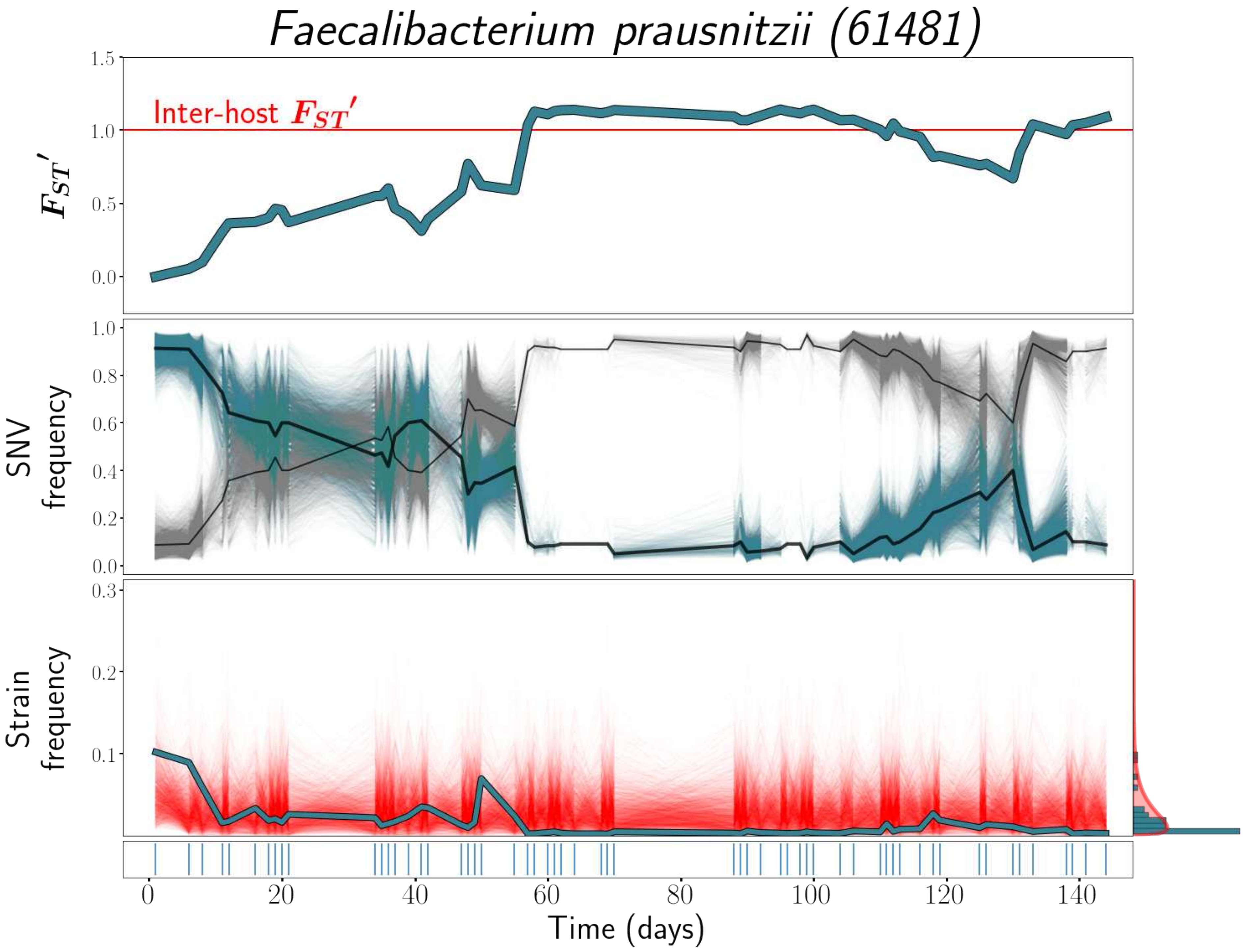
Faecalibacterium prausnitzii (61481)
Inter-host $F_{ST}'$
$F_{ST}'$
1.5
1.0
0.5
0.0
SNV frequency
1.0
0.8
0.6
0.4
0.2
0.0
Strain frequency
0.3
0.2
0.1
0
20
40
60
80
100
120
140
Time (days)

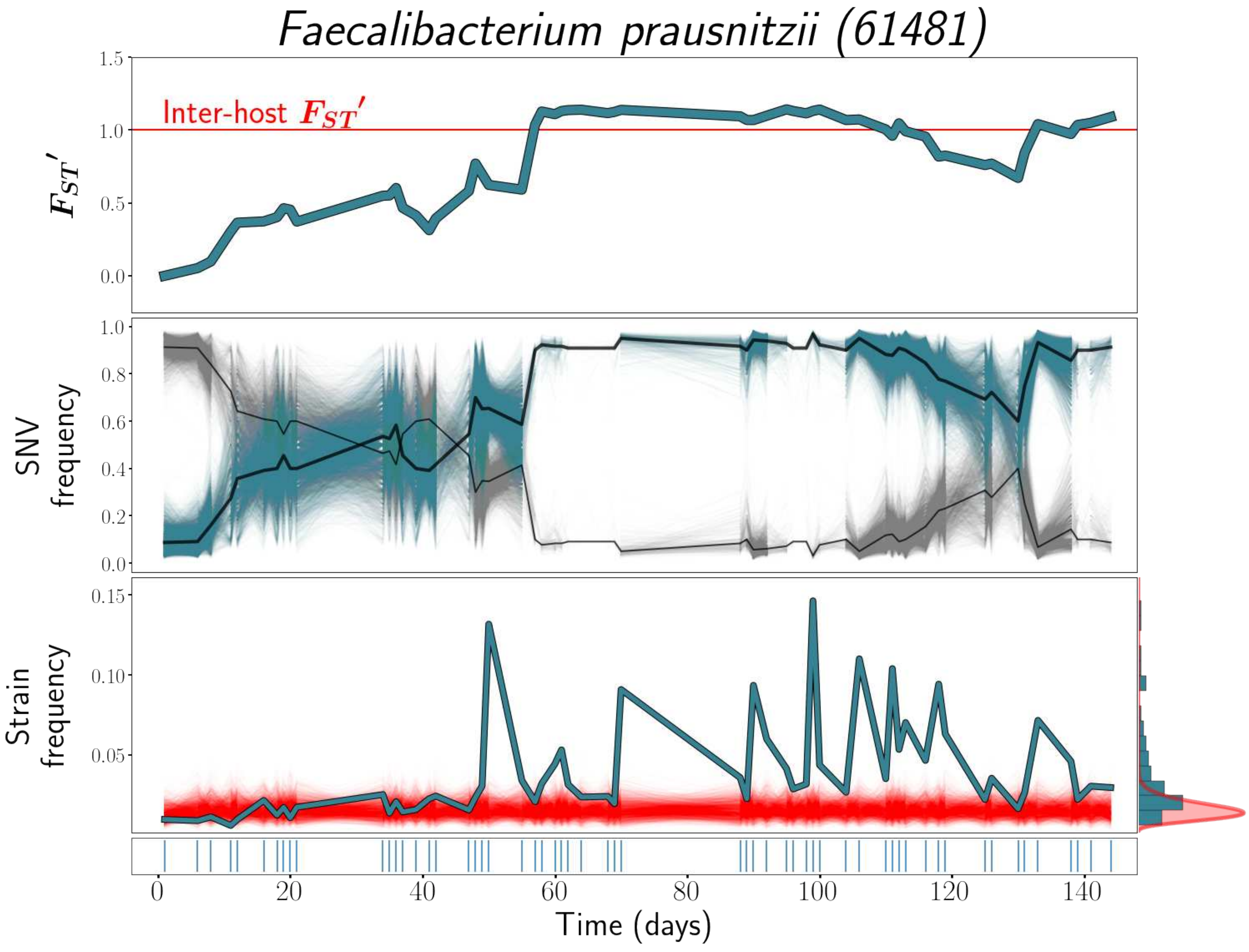

Faecalibacterium prausnitzii (61481)
Inter-host $F_{ST}'$
$F_{ST}'$
1.5
1.0
0.5
0.0
SNV frequency
1.0
0.8
0.6
0.4
0.2
0.0
Strain frequency
0.15
0.10
0.05
0
20
40
60
80
100
120
140
Time (days)

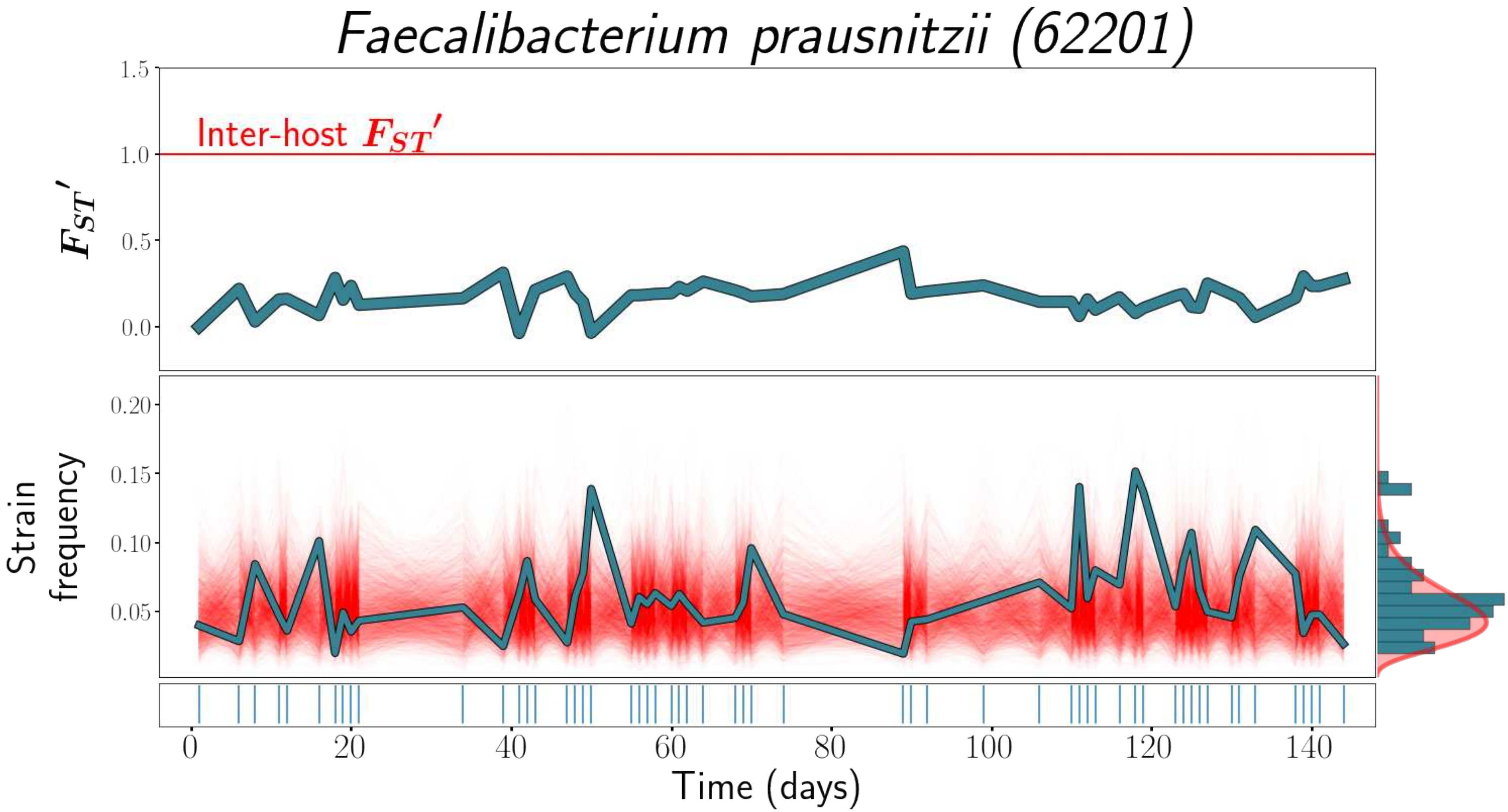

Faecalibacterium prausnitzii (62201)
Inter-host $F_{ST}'$
$F_{ST}'$
1.5
1.0
0.5
0.0
Strain frequency
0.20
0.15
0.10
0.05
0
20
40
60
80
100
120
140
Time (days)

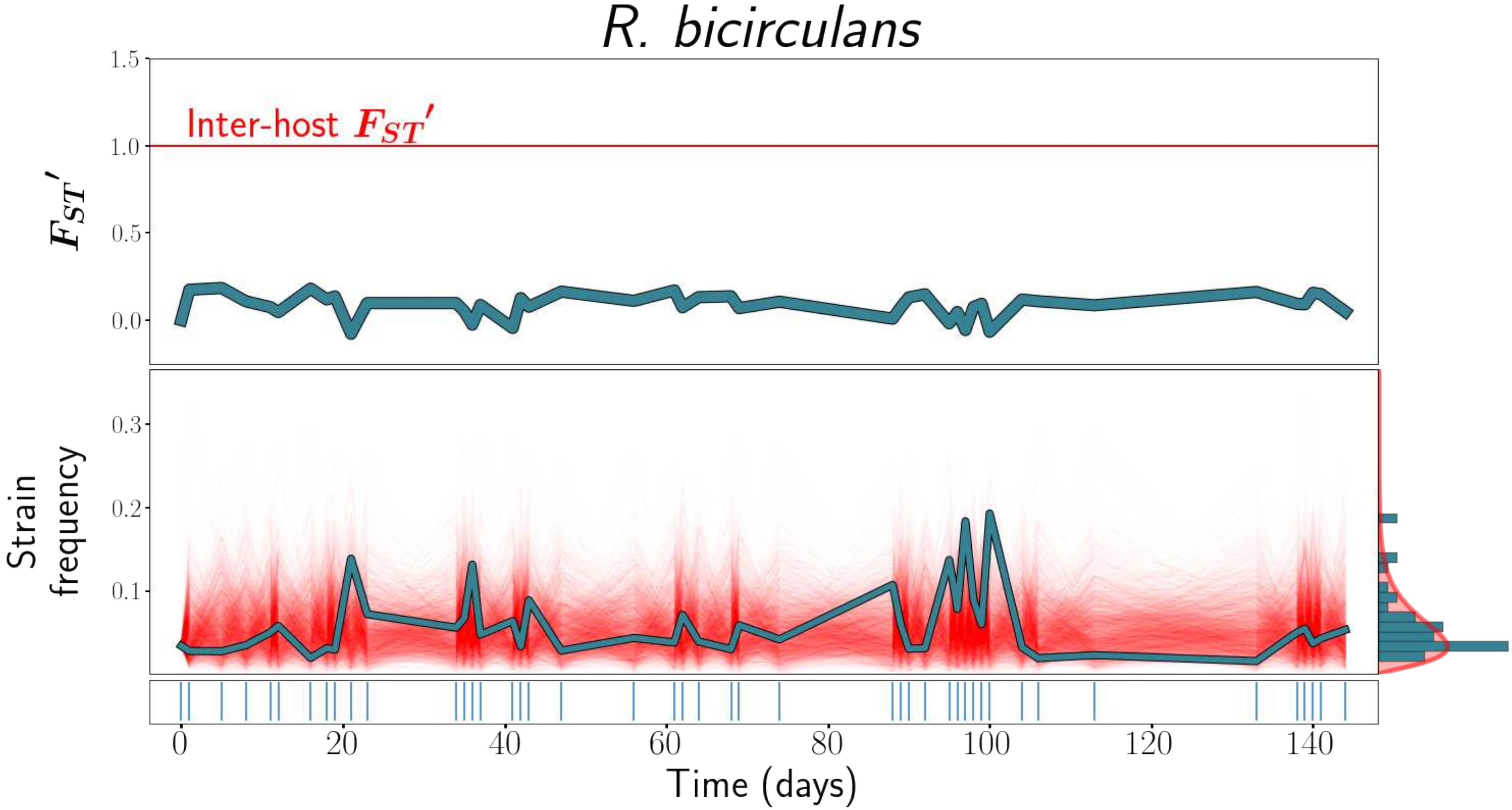

R. bicirculans
1.5
Inter-host $F_{ST}'$
1.0
$F_{ST}'$
0.5
0.0
0.3
Strain frequency
0.2
0.1
0
20
40
60
80
100
120
140
Time (days)

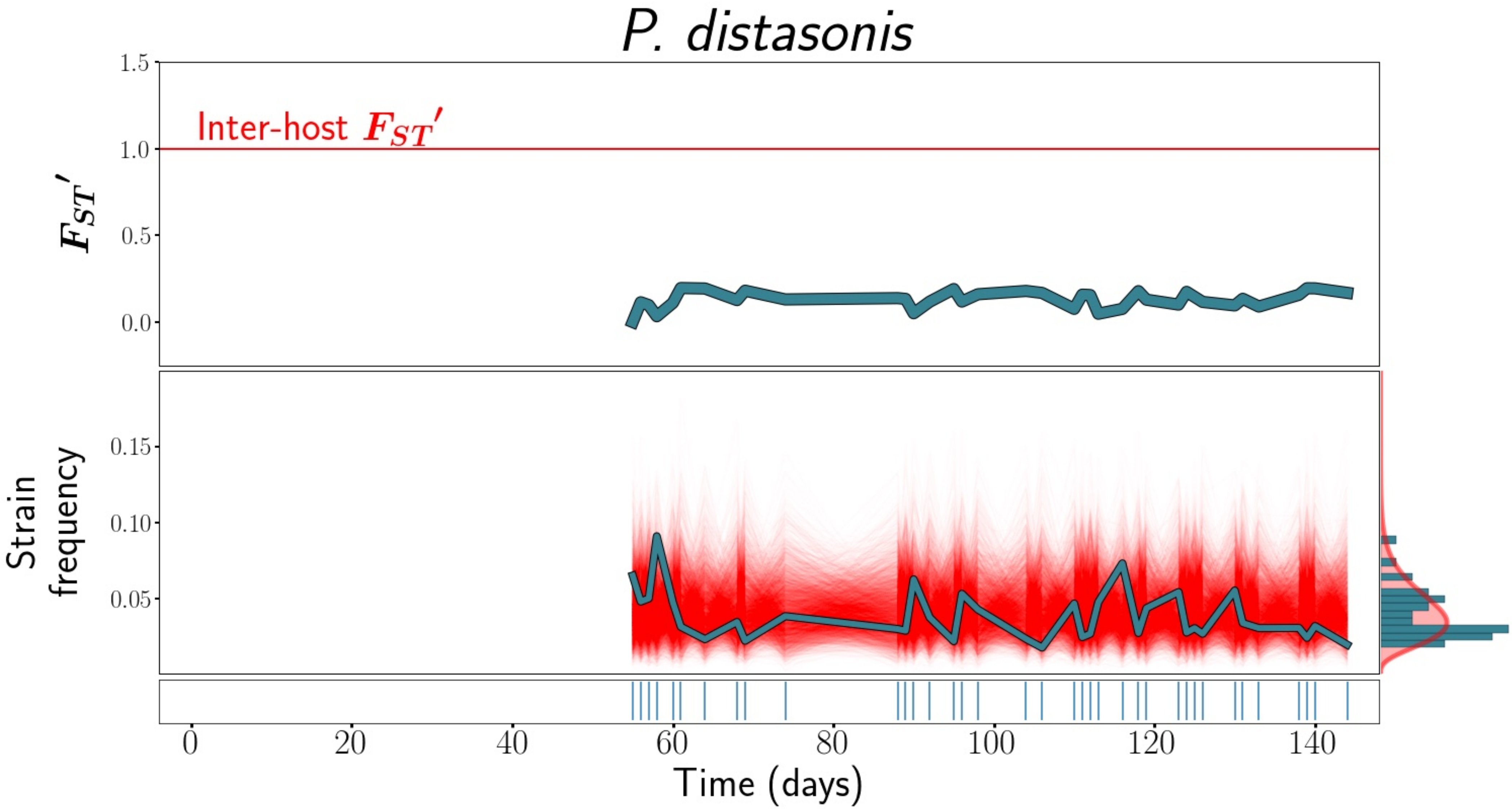

P. distasonis
Inter-host $F_{ST}'$
$F_{ST}'$
1.5
1.0
0.5
0.0
Strain frequency
0.15
0.10
0.05
0
20
40
60
80
100
120
140
Time (days)
